# Generalized Biomolecular Modeling and Design with RoseTTAFold All-Atom

**DOI:** 10.1101/2023.10.09.561603

**Authors:** Rohith Krishna, Jue Wang, Woody Ahern, Pascal Sturmfels, Preetham Venkatesh, Indrek Kalvet, Gyu Rie Lee, Felix S Morey-Burrows, Ivan Anishchenko, Ian R Humphreys, Ryan McHugh, Dionne Vafeados, Xinting Li, George A Sutherland, Andrew Hitchcock, C Neil Hunter, Minkyung Baek, Frank DiMaio, David Baker

**Affiliations:** Department of Biochemistry, University of Washington, Seattle, WA 98105, USA; Institute for Protein Design, University of Washington, Seattle, WA 98105, USA; Paul G. Allen School of Computer Science and Engineering, University of Washington, Seattle, WA 98105, USA; Graduate Program in Biological Physics, Structure and Design, University of Washington, Seattle, WA 98105, USA; School of Biosciences, University of Sheffield, Sheffield, S10 2TN, UK; School of Biological Sciences, Seoul National University, Seoul 08826, Republic of Korea; Howard Hughes Medical Institute, University of Washington, Seattle, WA 98105, USA

## Abstract

Although AlphaFold2 (AF2) and RoseTTAFold (RF) have transformed structural biology by enabling high-accuracy protein structure modeling, they are unable to model covalent modifications or interactions with small molecules and other non-protein molecules that can play key roles in biological function. Here, we describe RoseTTAFold All-Atom (RFAA), a deep network capable of modeling full biological assemblies containing proteins, nucleic acids, small molecules, metals, and covalent modifications given the sequences of the polymers and the atomic bonded geometry of the small molecules and covalent modifications. Following training on structures of full biological assemblies in the Protein Data Bank (PDB), RFAA has comparable protein structure prediction accuracy to AF2, excellent performance in CAMEO for flexible backbone small molecule docking, and reasonable prediction accuracy for protein covalent modifications and assemblies of proteins with multiple nucleic acid chains and small molecules which, to our knowledge, no existing method can model simultaneously. By fine-tuning on diffusive denoising tasks, we develop RFdiffusion All-Atom (RFdiffusionAA), which generates binding pockets by directly building protein structures around small molecules and other non-protein molecules. Starting from random distributions of amino acid residues surrounding target small molecules, we design and experimentally validate proteins that bind the cardiac disease therapeutic digoxigenin, the enzymatic cofactor heme, and optically active bilin molecules with potential for expanding the range of wavelengths captured by photosynthesis. We anticipate that RFAA and RFdiffusionAA will be widely useful for modeling and designing complex biomolecular systems.

---

The deep neural networks AlphaFold2 (AF2)(*1*) and RoseTTAFold (RF)(*2*) enable high-accuracy prediction of protein structures from amino acid sequence information alone. However, in nature, proteins rarely act alone; they form complexes with other proteins in cell signaling, interact with DNA and RNA during transcription and translation, and interact with small molecules both covalently and noncovalently during metabolism. Modeling such general biomolecular assemblies composed of polypeptide chains, covalently modified amino acids, nucleic acid chains, and arbitrary small molecules remains an outstanding challenge; one approach is to model the protein chains using AF2 or RF, and then successively add in the non-protein components using classical docking methods(*3–9*) but systematically evaluating and optimizing such procedures is not straightforward. RF has been extended to model both protein and nucleic acids by increasing the size of the residue alphabet to 28 (20 amino acids, four DNA bases, and four RNA bases) with RoseTTAFold nucleic acid (RFNA) (*10*), but general biomolecular system modeling is a more challenging problem given the great diversity of possible small molecule components. An approach capable of accurately predicting the three-dimensional structures of biomolecular assemblies starting only from knowledge of the constituent molecules (and not their 3D structures) would have broad impact on structural biology and drug discovery, and also open the door to deep learning-based design of protein-small molecule assemblies.

We set out to develop a structure prediction method capable of generating 3D coordinates for all atoms of a biological unit, including proteins, nucleic acids, small molecules, metals, and chemical modifications (Figure 1A). The first obstacle we faced in taking on the broader challenge of generalized biomolecular system modeling was how to represent the components. Existing protein structure prediction networks represent proteins as linear chains of amino acids, and this representation can be readily extended to nucleic acids. However, many of the small molecules that proteins interact with are not polymers, and it is unclear how to model them as a linear sequence. A natural way to represent the bonded structure of small molecules is as graphs whose nodes are atoms and whose edges represent bond connectivity. This graph representation is not suitable for proteins, as they contain many thousands of atoms, and hence, modeling whole proteins at the atomic level is computationally intractable. To overcome this limitation, we sought to combine a sequence-based description of biopolymers (proteins and nucleic acids) with an atomic graph representation of small molecules and protein covalent modifications.

**Figure 1.**
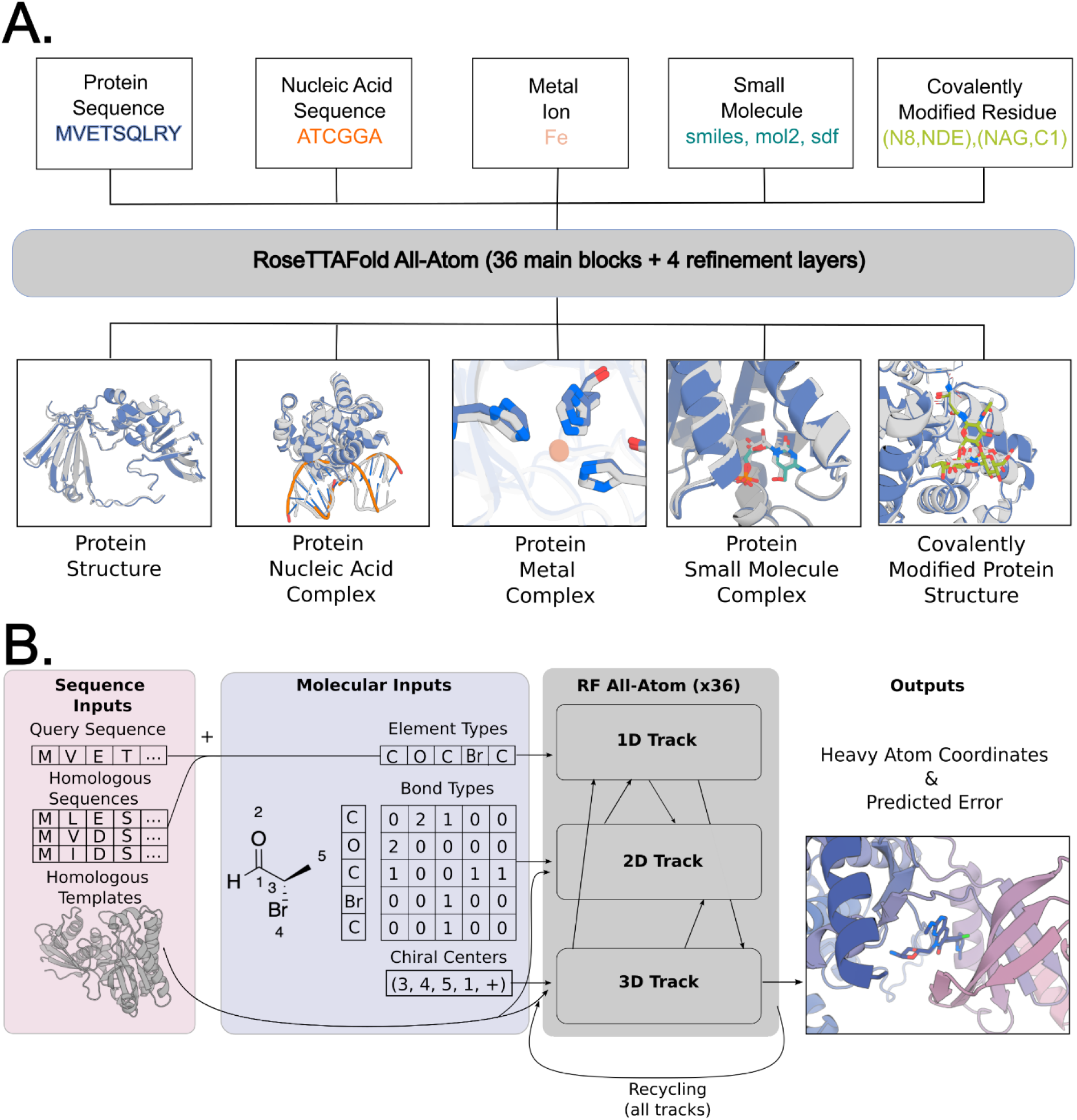
General biomolecular modeling with RoseTTAFold All-Atom. **A)** RFAA takes input information about the molecular composition of the biomolecular assembly to be modeled, including protein amino acid and nucleic acid base sequences, metal ions, small molecule bonded structure, and covalent bonds between small molecules and proteins. **B)** Processing of molecular input information. Small molecule information is parsed into element types (46 possible types), bond types, and chiral centers. Covalent bonds between proteins and small molecules are provided as a separate token in the bond adjacency matrix. The three-track architecture mixes 1D, 2D, and 3D information and predicts all-atom coordinates and model confidence.

## Generalizing Structure Prediction to All Biomolecules

We modeled the network architecture after the RoseTTAFold2 (RF2) protein structure prediction network, which accepts 1D sequence information, 2D pairwise distance information from homologous templates, and 3D coordinate information and iteratively improves predicted structures through many hidden layers(*11*). We retain the representations of protein and nucleic acid chains from RF2 and represent arbitrary small molecules as atom-bond graphs. To the 1D track, we input the chemical element type of each non-polymer atom; to the 2D track, the chemical bonds between atoms; and to the 3D track, information on chirality [whether chiral centers are (r) or (s)]. For the 1D track, we supplement the 20 residue and eight nucleic acid base representation in RFNA with 46 new element type tokens representing the most common element types found in the Protein Data Bank (PDB) (Table S6). For the 2D track atom-bond embedding, we encode pairwise information about whether bonds between pairs of atoms are single, double, triple, or aromatic bonds. These features are linearly embedded and summed with the initial pair features at the beginning of every recycle of the network, allowing the network to learn about bond lengths, angles, and planarity. Since the 1D and 2D representations in the network are invariant to reflections, we encode stereochemistry information in the third track by specifying the sign of angles between the atoms surrounding each chiral center (Fig S1); at each block in the 3D track the gradient of the deviation of the actual angles from the ideal values (with respect to the current coordinates) is computed and provided as an input feature to the subsequent block (Figure 1B).

Unlike proteins and nucleic acid sequences, molecular graphs are permutation invariant, and hence, the network should make the same prediction irrespective of small molecule element token order. In AF2 and RF2, the sequence order of amino acids and bases is represented by a relative position encoding; for atoms, we omit such an encoding and leverage the permutation invariance of the network’s attention mechanisms. We also modify the coordinate updates: in AF2 and RF, protein residues are represented by the coordinates of the C*α* and the orientation of the N-C*α*-C rigid frame (or the P coordinate and the OP1-P-OP2 frame orientation in RFNA) and along the 3D track the network generates rotational updates to each frame orientation and translational updates to each coordinate. To generalize this in RFAA, heavy atom coordinates are added to the 3D track and move independently based only on a predicted translational update to their position. Thus, immediately after input, the full system is represented as a disconnected gas of amino acid residues, nucleic acid bases, and freely moving atoms, which is successively transformed through the many blocks of the network into physically plausible assembly structures. For the loss function to guide parameter optimization, we develop an all-atom version of the Frame Aligned Point Error (FAPE) loss introduced in AF2 by defining coordinate frames for each atom in an arbitrary molecule based on the identities of its bonded neighbors and, as with residue based FAPE, successively aligning each coordinate frame and computing the coordinate error on the surrounding atoms (Figure 2A; for greater sensitivity to small molecule geometry, we upweight contributions involving atoms; see Supplemental Methods). In addition to atomic coordinates, the network predicts atom and residue-wise confidence (pLDDT) and pairwise confidence (PAE) metrics to enable users to identify high-quality predictions. A full description of the RFAA architecture is provided in the Supplemental Methods.

**Figure 2.**
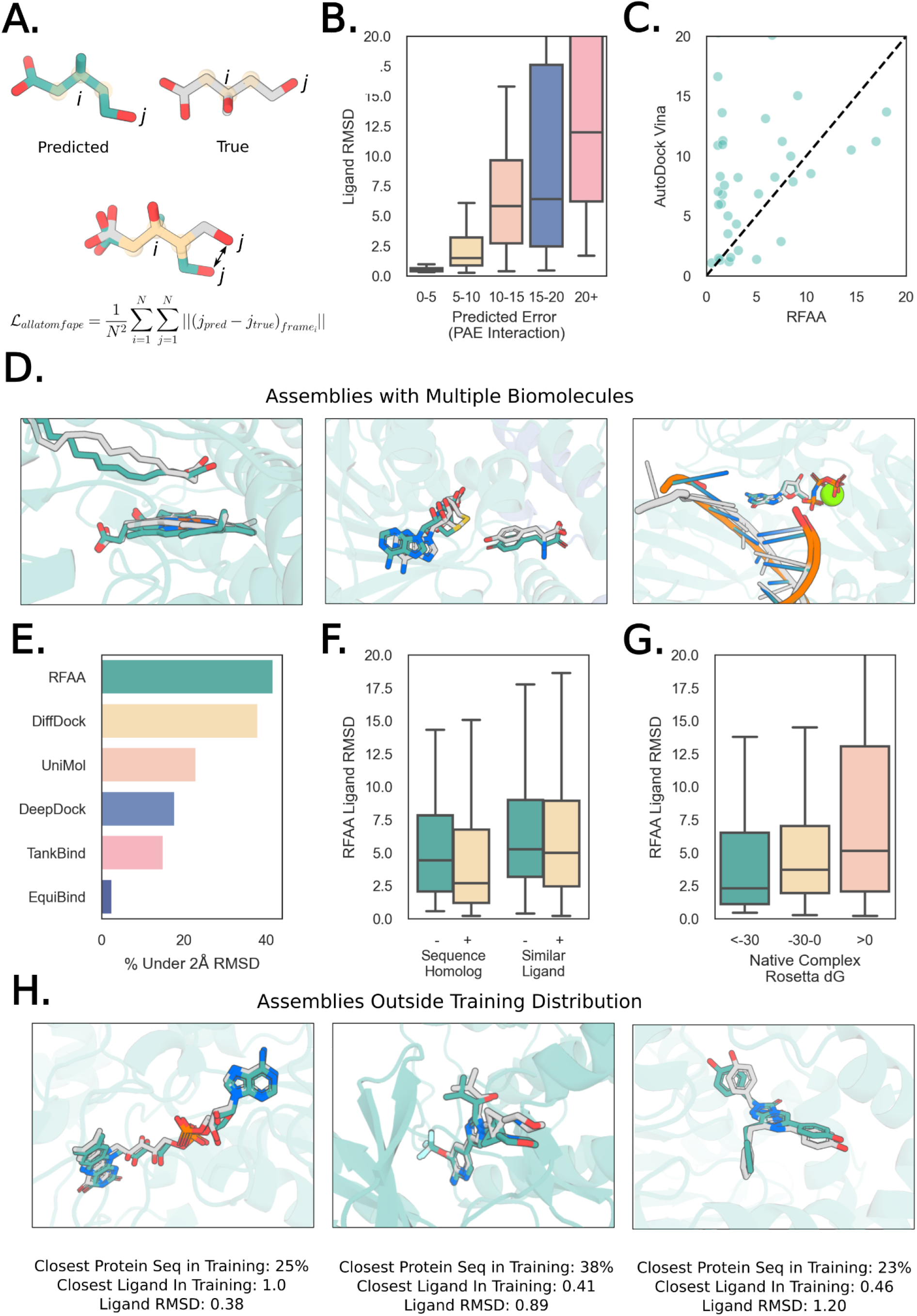
RoseTTAFold All-Atom can accurately predict protein-small molecule complex structures. All panels: Predicted protein structure (aligned to native): transparent teal, predicted ligand conformation: teal, native ligand conformation: gray. All boxplots cut off at 20Å for clarity.**A)** Every “atom” node is assigned a local coordinate frame based on the identities of its neighbors. To compute the main loss in the network, we align each atom’s coordinate frame in the predicted and true structures and measure the error over all the other atoms. **B)** Model accuracy correlates with error predictions. Computed for CAMEO targets (05/20/23-7/29/23; 261 protein-small molecule interfaces). Ligand RMSD was computed by CAMEO organizers. **C)** RFAA outperforms AutoDock Vina on CAMEO targets (Week 8/12/23-09/02/23; 149 protein-small molecule interfaces). Both servers have to model the protein, find pockets for all ligands present in the solved structure, and the correct docks for all ligands. Ligand RMSD for both servers was computed by CAMEO organizers, AutoDock Vina server set up by CAMEO organizers. **D)** Three examples of successful predictions with multiple biomolecules. From left to right: novel fatty acid decarboxylase (PDB ID: 8d8p; from CAMEO) with a heme cofactor and a lipid substrate, a dimeric tyrosine methyltransferase (PDB ID: 7ux6; CASP15 Target: T1124) with an s-adenosyl homocysteine and tyrosine interaction and a DNA polymerase (PDB ID: 7u7w). **E)** Comparison to other deep learning-based docking methods. In this case, each method was applied in their respective training regime. For RFAA this means only having sequence and minimal atomic graph inputs, but for other methods, this involves providing the bound crystal structure. Ligand RMSD was computed using PoseBusters suite, and a single example present in our training set was removed for all methods in comparison. **F)** Comparison of RFAA predictions on recently solved PDBs that are novel compared to the training set (Homolog >30% sequence identity, Similar Ligand >0.5 Tanimoto Similarity). Each set is clustered based on sequence/ligand similarity, and the lowest Ligand RMSD cluster representative is chosen for each (to answer the question of what is the best the network can do on these inputs). **G)** Comparison of RFAA prediction accuracy to Rosetta ΔG energy estimates for the native complex (over 940 cases that were successfully processed by Rosetta). RFAA makes more accurate predictions for native complexes with low Rosetta energy, suggesting that the network has learned principles that correlate with physics, not just whether the structure was seen in training. **H)** Three examples of successful predictions with low similarity to the training set. From left to right: a putative L-amino acid oxidase (PDB ID: 7eme), complex of DLK bound to an inhibitor (PDB ID: 8ous), a *Renilla* luciferase bound to an azacoelenterazine (non-native substrate; PDB ID: 7xqr).

## Training RFAA

From the PDB, we curated a protein-biomolecule dataset including protein-small molecule, protein-metal, and covalently modified protein complexes, filtering out common solvents and crystallization additives. Following clustering (30% sequence identity) to avoid bias towards overrepresented structures, we obtained 121,800 protein-small molecule structures in 5,662 clusters, 112,546 protein-metal complexes in 5,662 clusters, and 12,689 structures with covalently modified amino acids in 1,099 clusters for training. To help the network learn the general properties of small molecules rather than features specific to the molecules in the PDB, we supplemented the training set with small molecule crystal structures from the Cambridge Structural Database(*12*). Each training example is sampled uniformly from the set of organic non-polymeric molecules, and the network predicts the coordinates for the asymmetric unit given atomic graph information. To further help the network learn about general atomic interactions, we take advantage of the commonalities between atomic interactions within proteins and many of the atomic interactions between proteins and small molecules, and augment the training data by inputting portions of proteins as atoms rather than residues (a process we term *atomization*). We atomize randomly selected subsets of three to five contiguous residues, by deleting the sequence and template features and providing instead atom, bond, and chirality information for the atoms in those residues. For example, an alanine would be replaced by five atom tokens (one for each heavy atom). Since the atoms are still part of the polypeptide chain, we provide the relative position of the atom tokens with respect to the other residue tokens by adding an extra bond token that corresponds to an “atom-to-residue” bond and develop a positional encoding to account for atom-residue bonds (Supplemental Methods). To increase prediction accuracy on biological polymers, we train the network on protein monomer, protein complex, and protein-nucleic acid complex examples as previously described(*10*, *11*). All examples were cropped to have 256 tokens during the initial stages of training and 375 tokens during fine-tuning. The progress of training was monitored using independent validation sets consisting of 10% of the protein sequence clusters (see Supplementary Information Table 4).

Given the much broader set of possible inputs, RFAA can model full biomolecular systems unlike previous protein-only deep learning architectures(*13–15*). In the following sections, we describe the performance of RFAA on different structure modeling tasks. We adopted the philosophy that a single model trained on all available data over all modalities would have the greatest ability to generalize and be more accessible than a series of models specialized for specific problems–it is possible that better performance could be obtained by problem specific fine-tuning.

## Predicting Protein-Small Molecule Complexes

There has been considerable recent effort in developing deep learning methods for ligand docking into known crystal structures(*5*, *16–20*). While these methods have shown promising accuracy, they generally require *a priori* knowledge of the bound conformation of the protein and are also unable to model additional atomic contexts such as cofactors, metal ions, or covalent modifications. We reasoned that RFAA could be particularly useful for two difficult modeling challenges: the general protein-ligand docking problem while allowing for flexibility in the target and docking multiple small molecules or nucleic acid chains simultaneously.

To enable blind testing of RFAA prediction performance, we enrolled an RFAA server in the blind CAMEO ligand docking evaluation, which carries out predictions using a series of servers on all structures submitted to the PDB each week and evaluates their performance(*21–23*). While not all predictions are accurate, we use the network’s predicted pairwise error between protein chains and small molecule chains (PAE Interaction) to identify accurate predictions; across CAMEO targets, 43% of cases are predicted with high confidence (PAE Interaction < 10), and 77% of those high-confidence structures are predicted with < 2Å ligand RMSD (Figure 2B). One of the other servers is an implementation of a leading non-deep learning protein small molecule docking method AutoDock Vina by the CAMEO organizers that predicts the protein structure by homology modeling(*24–29*), runs AutoDock to dock the small molecule, and ranks the poses using the Vina scoring function (*9*, *24*). RFAA consistently outperformed the other servers in CAMEO on protein-small molecule modeling; for example, on cases modeled by both the RFAA and the AutoDock Vina servers, RFAA models 32% of cases successfully (< 2Å ligand RMSD) compared to 8% for the Vina server (Figure 2B; the Vina performance by an expert would likely be considerably improved because of the complexities of fully automatic multiple step modeling pipelines). The most common failure mode is the placement of small molecules in the correct pockets but not in the correct orientation (Figure S3; for further exploration of failure modes, see Supplemental Methods).

One strength of RFAA compared to previous methods is that the network is able to jointly predict interactions between proteins and multiple non-protein ligands in a single forward pass. Figure 2D shows three examples of recently solved structures with three or more components for which RFAA predictions had <2Å ligand RMSD (when the proteins are aligned). These cases include a novel fatty acid decarboxylase (PDB ID: 8d8p) with a heme cofactor and a lipid substrate, a dimeric tyrosine methyltransferase (PDB ID: 7ux6) with an s-adenosyl homocysteine and tyrosine interaction, and a DNA polymerase (PDB ID: 7u7w) with a bound DNA, non-hydrolyzable guanine triphosphate and magnesium ion (Figure 2D; the network received no examples of higher order assemblies containing proteins, small molecules, and nucleic acids during training). To our knowledge, no other current methods can model arbitrary higher-order biomolecular complexes, which can include multiple proteins, small molecules, metal ions, and nucleic acids.

An often used but not very stringent test of protein-ligand docking methods is the ability of a method to dock a small molecule with a protein target given the crystal structure of the backbone and side chains of the protein in complex with the small molecule (in real-world problems, such a “bound” crystal structure is never available). We compared RFAA to other deep learning based docking tools when they are provided with more privileged information on a set of recent PDBs curated in (*30*) (the test structures are outside of the training sets of all the methods; the other methods were not enrolled in CAMEO, so we could not carry out a blind test). RFAA predicts 42% of complexes successfully compared to DiffDock, which predicts 38% of complexes successfully (Figure 2D; RFAA predicts the protein backbone and side chains in addition to the small molecule dock, whereas DiffDock receives the crystal structure of the protein from the bound complex as input). In cases where both the bound protein structure and the pocket residues are provided, physics-based methods such as AutoDock Vina outperform RFAA (52% vs 42%), which has the much harder task of predicting both the protein backbone and sidechain details and the dock from sequence alone (Figure S4A).

We investigated whether additional biomolecular context can improve protein structure prediction. In cases where RFAA predicts ligand placement with high confidence and RF2 has high confidence (PAE Interaction <10 and pLDDT>0.8 respectively), RFAA makes higher accuracy protein structure predictions than RF2 (Fig S3A), indicating that training with ligand context can improve overall protein prediction accuracy. Some examples of shifts predicted by RFAA but not by RF2 include domain movements, subtle backbone movements, and flipping of side chain rotamers to accommodate the ligand in the pocket (Figure S3B-C).

During training, RFAA should be learning general features of protein-small molecule interactions, but it could also be memorizing likely binding modes of molecules to different sequence families. To evaluate the ability of the model to generalize to new cases, we assembled a dataset of recent PDB entries with small molecules bound that were deposited after the cutoff date for our training set, and predicted full structure models for all 5,421 complexes (1,681 protein sequence clusters at 30% sequence identity). The network performs better for clusters with overlap with the training set, but does generalize to novel clusters (41% vs. 23% success rate)(Figure 2F). We observe a similar pattern for ligand clusters (across 3,261 ligand clusters); while the network makes more accurate predictions for ligands seen in training, it also can make accurate predictions on ligands that are not similar to those in training (<0.5 Tanimoto similarity; 20% vs. 16% success rate) (Figure 2F). Training on more extensive datasets will likely be necessary to generate consistently accurate predictions for new protein-small molecule complexes on par with the accuracy deep networks can achieve on protein systems alone.

To determine the extent to which the network is learning general principles of protein-small molecule interactions, we investigated the correlation between prediction accuracy and physically based correlates of protein-small molecule interaction affinity. We found that predictions for protein-small molecule complexes with high predicted affinity (by Rosetta ΔG)(*31*, *32*)) were more accurate than predictions for complexes predicted to bind weakly (Figure 2G; 50%, 25%, and 22% success rates for <-30, -30-0, and >0 Rosetta Energy Units respectively), further suggesting that RFAA is learning general principles rather than just memorizing global protein-small molecule interactions patterns.

## Predicting Structures of Covalent Modifications to Proteins

Many essential protein functions, such as receptor signaling, immune evasion, and enzyme activity, involve covalent modifications of amino acid side chains with sugars, phosphates, lipids, and other molecules (*33–36*). RFAA models such modifications by treating the residue and chemical moiety as atoms (with the corresponding covalent bond to the atom token in the residue) and the rest of the protein structure as residues (Figure 3A).

**Figure 3.**
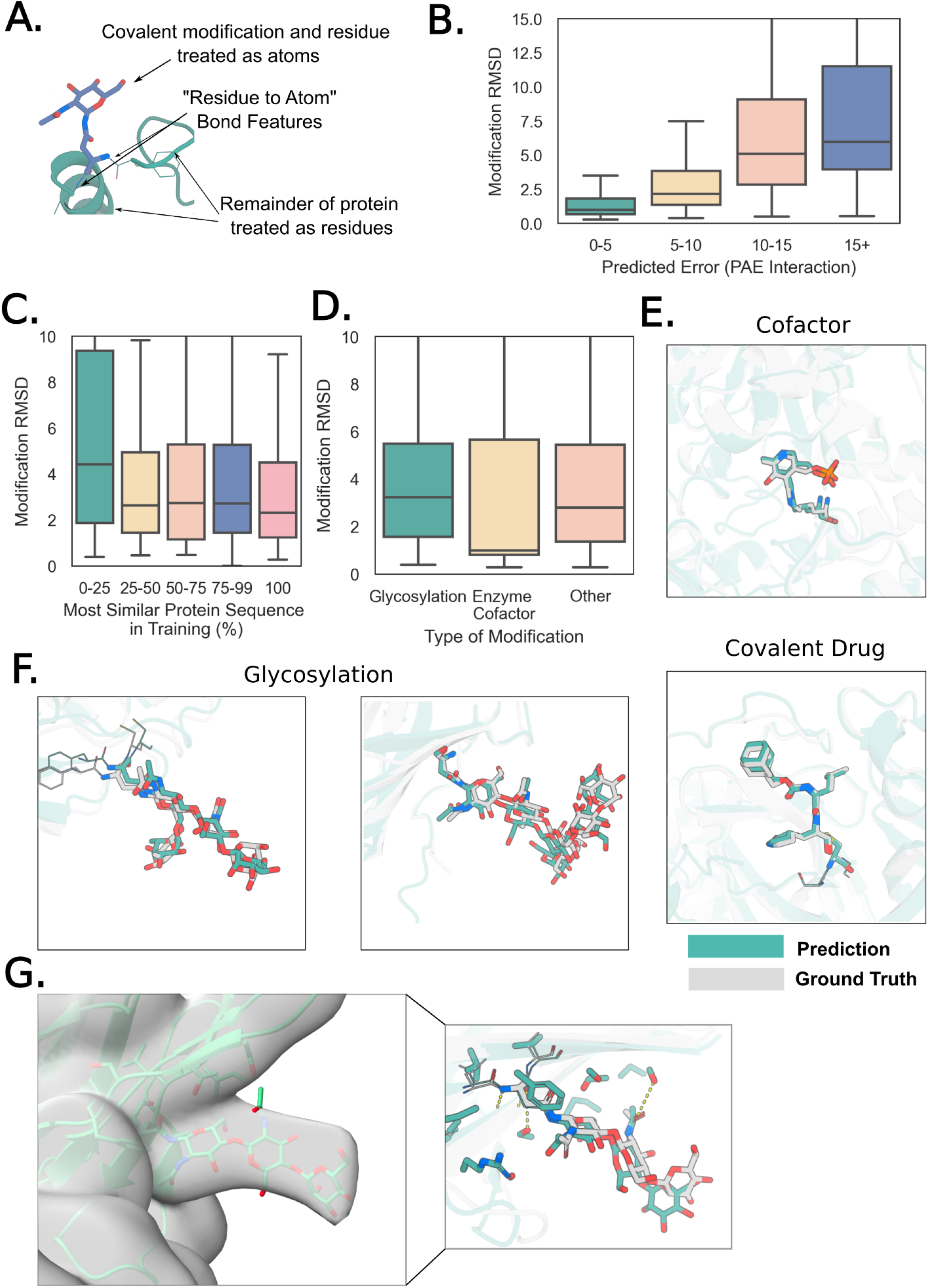
Accurate prediction of protein covalent modifications. All panels: transparent teal: predicted protein structure, transparent gray: native structure, teal: predicted covalent modification, gray: native covalent modification. **A)** Schematic describing how RFAA models covalent modifications to proteins. The chemical moiety that modifies the residue and the residue are modeled as atom nodes, and the rest of the protein is modeled as residues (with MSA and template inputs). **B)** Model accuracy correlates with predicted error on a set of 938 recently solved structures with covalent modifications. Modification RMSD is computed by aligning the protein structure within 10Å and computing RMSD over the modified residue and chemical modification. Boxplot cut off at 15Å for clarity. **C)** Comparison of sequence identity to training set and model accuracy. Models are generally accurate even with low sequence homology to the training set. **D)** Comparison of model accuracy for different types of covalent modifications. **E)** *Top*: Example of successfully predicted covalently linked enzyme cofactor (PDB ID: 7ny2), which is a structure of a human branched chain amino acid aminotransferase (BCAT-1). *Bottom*: example of a covalently bound drug candidate (PDB ID: 7lkt), which is SARS-CoV-2 3CL protease bound covalently to an inhibitor with a conformationally constrained cyclohexane moiety. **F)** Accurate predictions of glycans on the N-acetylglucosamine-1-phosphotransferase (GNPT) gamma subunit (PDB ID: 7s69), human sperm TMEM ectodomain (PDB ID: 7ux0). **G)** *Left:* Fit of predicted model into EM density for a glycan on the IL-27 beta subunit (PDB ID: 7u7n; EMDB: 26382) *Right:* zoom in of interactions between predicted sidechains and predicted glycan. The network accurately predicts hydrogen bonding between a serine in the protein and the second monosaccharide in the glycan chain.

We benchmarked the performance of the network in covalent modification structure prediction on 931 recent entries in the PDB (post-May, 2020), and found that the network made accurate predictions (Modification RMSD<2.5Å) in 46% of cases (where Modification RMSD is defined as RMSD of the modified residue and chemical modification when the rest of the protein is aligned). As in the protein-small molecule complex case, confident predictions tend to be more accurate: 60% of structures are predicted with high confidence (PAE Interaction <10), and 63% of those predictions are accurate (<2.5Å modification RMSD) (Figure 3B). While the network makes slightly more accurate predictions on cases with sequence similarity (>25% identity) to proteins in the training set, there are still many cases (27.5%) that do not have sequence overlap to the training set that are predicted with high accuracy (Figure 3C). RFAA models interactions with covalently bound cofactors and covalently bound drugs with median RMSDs of 0.99Å and 2.8Å respectively (Figure 3D-E).

Prediction of glycan structure has applications in therapeutics, vaccines, and diagnostics(*37–39*). RFAA can accurately model carbohydrate groups introduced by glycosylations with a median RMSD over our test set of 3.2Å (Figure 3D). RFAA successfully predicts glycans on the N-acetylglucosamine-1-phosphotransferase (GNPT) gamma subunit (PDB ID: 7s69), human sperm TMEM ectodomain (PDB ID: 7ux0) and the IL-27 beta subunit (PDB ID: 7u7n), which have low sequence homology (<30%) to our training set (Figure 3F-G) and have multiple monosaccharides and different branching patterns. The latter structure was resolved by cryo-EM and the RFAA predicted glycan model fits well in the density (Figure 3G). The network is able to make accurate predictions of glycan interactions even when the sequences were distant from the sequences in the training set and on glycans with chains up to seven monosaccharides (Figure S5).

It is difficult to compare to other methods because, to our knowledge, previous deep learning based tools do not model covalent modifications to proteins. Accurate and robust modeling of covalent modifications in predicted structures should contribute to the understanding of biological function and mechanism.

## *De Novo* Small Molecule Binder Design

The design of proteins that bind small molecules is a grand challenge in protein design. Previous efforts have involved docking molecules into large sets of native or expert-curated protein scaffold structures(*40*)-(*41*). Recent work has shown diffusion approaches can generate proteins in the context of a protein target that bind with considerable affinity and specificity (*42*).

However, current deep learning based generative approaches do not explicitly model protein-ligand interactions, so they are not directly applicable to the small molecular binder design problem. In RFdiffusion, a heuristic attractive-repulsive potential encouraged the formation of pockets with shape complementarity to a target molecule, but the approach was unable to model the details of protein-small molecule interactions, and none of the designs were experimentally validated(*42*). A general method that can generate protein structures around small molecules and other non-protein targets to maximize favorable interactions could be broadly useful.

We reasoned that RFAA could enable protein design in the context of non-protein biomolecules following fine-tuning on structure denoising. We developed a diffusion model, RFdiffusion All-Atom (RFdiffusionAA), by training a denoising diffusion probabilistic model (DDPM) initialized with the RFAA structure-prediction weights to denoise corrupted protein structures conditioned on biomolecular contexts using the protein-small molecule dataset described above (Figure 4A). Native structures are noised through progressive shrinking and the addition of 3D Gaussian noise to the C*α* coordinates and Brownian motion on the manifold of rotations. While some protein diffusion models are trained unconditionally and incorporate conditional information through forms of guidance (*43*),(*44*), we train an explicitly conditional model that learns the distribution of proteins conditioned on biomolecular substructure. During training, substructures of the native complexes, ‘motifs’, are provided to the model as context. In design settings, we typically know the conformation of the small molecules we want to bind, so the ligand conformation is always included in the motif. Furthermore, to enable the inclusion of specific protein functional motifs when desired, we also train the network to scaffold a variety of discontiguous protein motifs both in the presence and absence of small molecules. To generate proteins, we initialize a Gaussian distribution of residue frames with randomized rotations around a fixed small molecule motif, and at each denoising step t, predict the fully denoised X_0_ state, and then update all coordinates and orientations (for residues) by taking a step towards this conformation while adding noise to match the distribution for X_t-1_. Similarly to RFdiffusion, we investigated the use of auxiliary potentials to influence trajectories to make more contacts between small molecules and binders. Generally, we found the use of such potentials unnecessary, though they can yield tighter interfaces for larger small molecules, as we demonstrate for FAD (Figure 4C).

**Figure 4.**
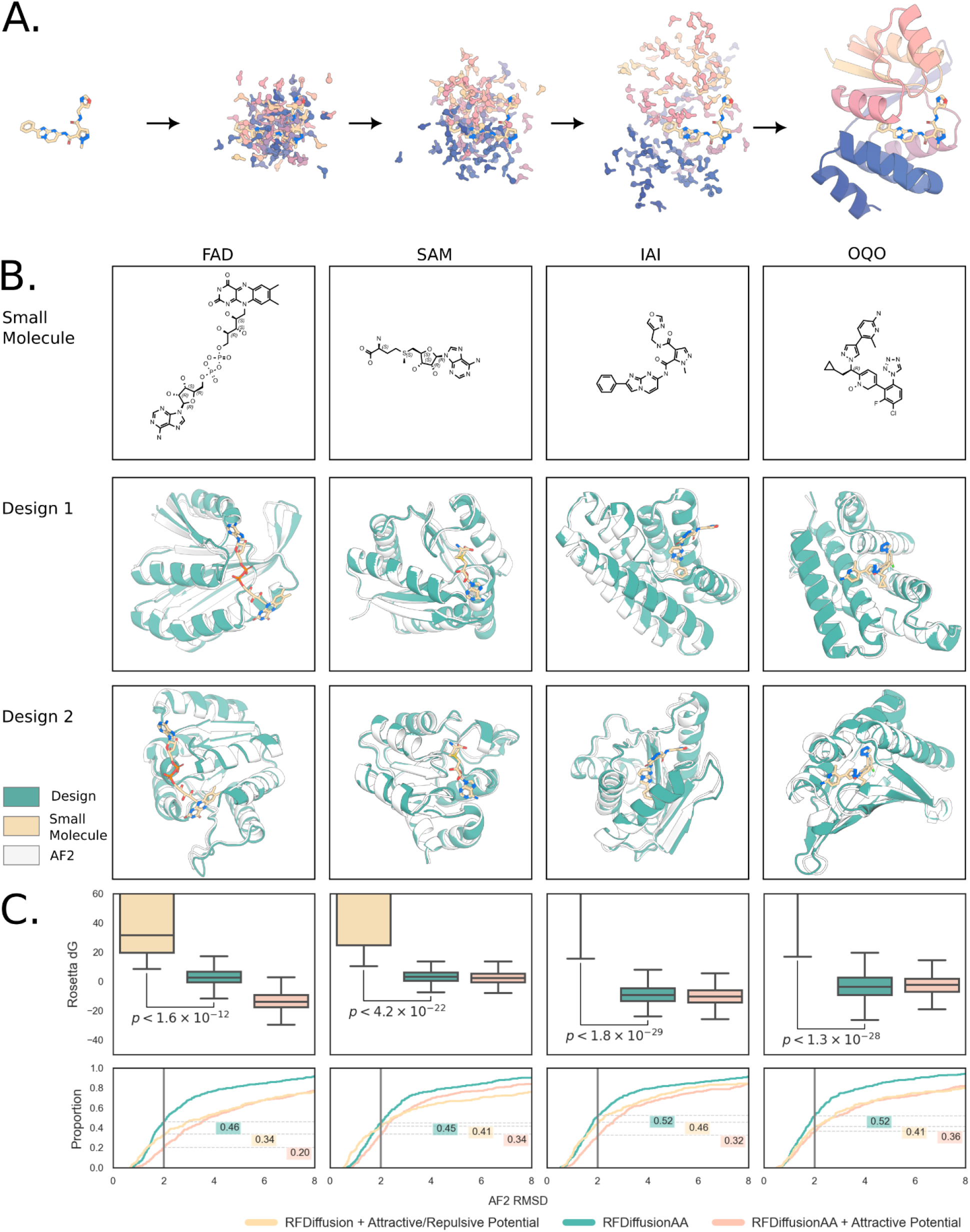
Small molecule binding protein design with RFdiffusion All-Atom. **A)** Schematic depicting the denoising of a residue gas into a small molecule binder. **B)** Binder design models for four representative small molecules (top row) are nearly identical to the AF2 pose prediction from single sequence alone. Two models are shown (second and third rows) to illustrate design diversity (for more extensive evaluation of diversity, see Figure S6). **C)** Comparison of protein-only RFdiffusion with the substrate modeled implicitly using an attractive/repulsive potential against RFdiffusionAA with and without attractive potential. *Top*: GALigandDock evaluated binding ΔG (minimum of eight LigandMPNN sequences). In all cases, RFdiffusionAA produces protein-ligand interfaces with a lower computed ΔG than RFdiffusion. *Bottom*: proportion of designs with RMSD to AF2 prediction less than the value specified by the x-axis (minimum of eight LigandMPNN sequences). The fraction of designs with less than 2Å RMSD to AF2 are highlighted; in all cases, RFdiffusionAA produces more self-consistent designs than RFdiffusion.

We evaluated RFdiffusionAA *in silico* by generating protein structures in the context of four diverse small molecules. Starting from random gaussian residue distributions surrounding each of the small molecules, iterative denoising yielded coherent protein backbones with pockets complementary to the small molecule target. We assign each backbone a sequence using an updated version of ProteinMPNN that can accept atomic context as conditional information (LigandMPNN)(*45*). Following sequence design using LigandMPNN, we score the designs using Rosetta GALigandDock (*31*) energy calculations to evaluate the quality of the small molecule interface and AF2 predictions to evaluate the extent the sequence encodes the designed structure. The *in silico* binding energy evaluations of RFdiffusionAA binders are far better (p<1.56E-12 in each small molecule case) than those obtained using a heuristic attractive/repulsive potential with protein-only RFdiffusion, the latter of which produces no interfaces with negative Rosetta ΔG as compared to 52% and 72% for RFdiffusionAA and RFdiffusionAA with an auxiliary contact potential respectively (Figure 4B). The standard method for *in silico* assessment of protein generative modeling is to measure the RMSD between the generated protein backbone and AF2’s prediction of that backbone from sequence alone (*42*)-(*46*). RFdiffusionAA’s small molecule binders have high consistency with AF2 predictions made from a single sequence, with at least 45% of structures having AF2 backbone RMSD < 2 Å in all small molecule binder design cases (Figure 4C). For each small molecule, RFdiffusionAA generates diverse protein structural solutions to the binding problem; two examples are shown for each case in Figure 4 (see Figure S6 for more detail on the diversity of the designs). Two of the small molecules are quite distinct from any of the molecules in the training set (Tanimoto similarity <= 0.5; IAI: 0.50, OQO: 0.46); the design metrics for the binders to these models were similar to those of previously seen ligands, suggesting that the model has considerable ability to generalize not only in protein topology and structure but also in interactions with non-protein targets. In 99% of cases, the most structurally similar protein in the training dataset does not possess the same ligand (see Figure S7 for more details on the novelty of the designs).

## Experimental Characterization of Designed Binders

We designed binders for three diverse small molecules: one with no protein motif, one with a single residue protein motif, and one with a four residue protein motif, produced the proteins in *E. coli*, and measured binding experimentally.

Digoxigenin is the aglycone of digoxin, a small molecule used to treat heart diseases with a narrow therapeutic window(*47*), and digoxigenin-binding proteins could help reduce toxicity(*48*). Previous attempts to design digoxigenin-binding proteins relied on protein scaffolds with experimentally determined structures and prespecified binding pockets and interacting motifs(*49*). However, this approach is not generally applicable since ideal scaffolds and specific binding interactions may not be available or known for arbitrary small molecules. We used RFdiffusionAA to design digoxigenin-binding backbones without any prior assumption about the protein-ligand interface or backbone structure (Figure 5A). Sequences were fitted to these backbones using LigandMPNN and Rosetta FastRelax (*50*). 4,416 designs selected based on consistency with AF2 predictions and Rosetta metrics (Supplemental Methods) were cloned into yeast and screened for binding using fluorescence-activated cell sorting (Supplemental Methods). Three hits with enriching binding signals were further characterized by protein purification and fluorescence polarization (FP). The tightest binder has a 10 nM *K*_d_ (Figure 5A) to digoxigenin and is stable at temperatures up to 95℃.

**Figure 5.**
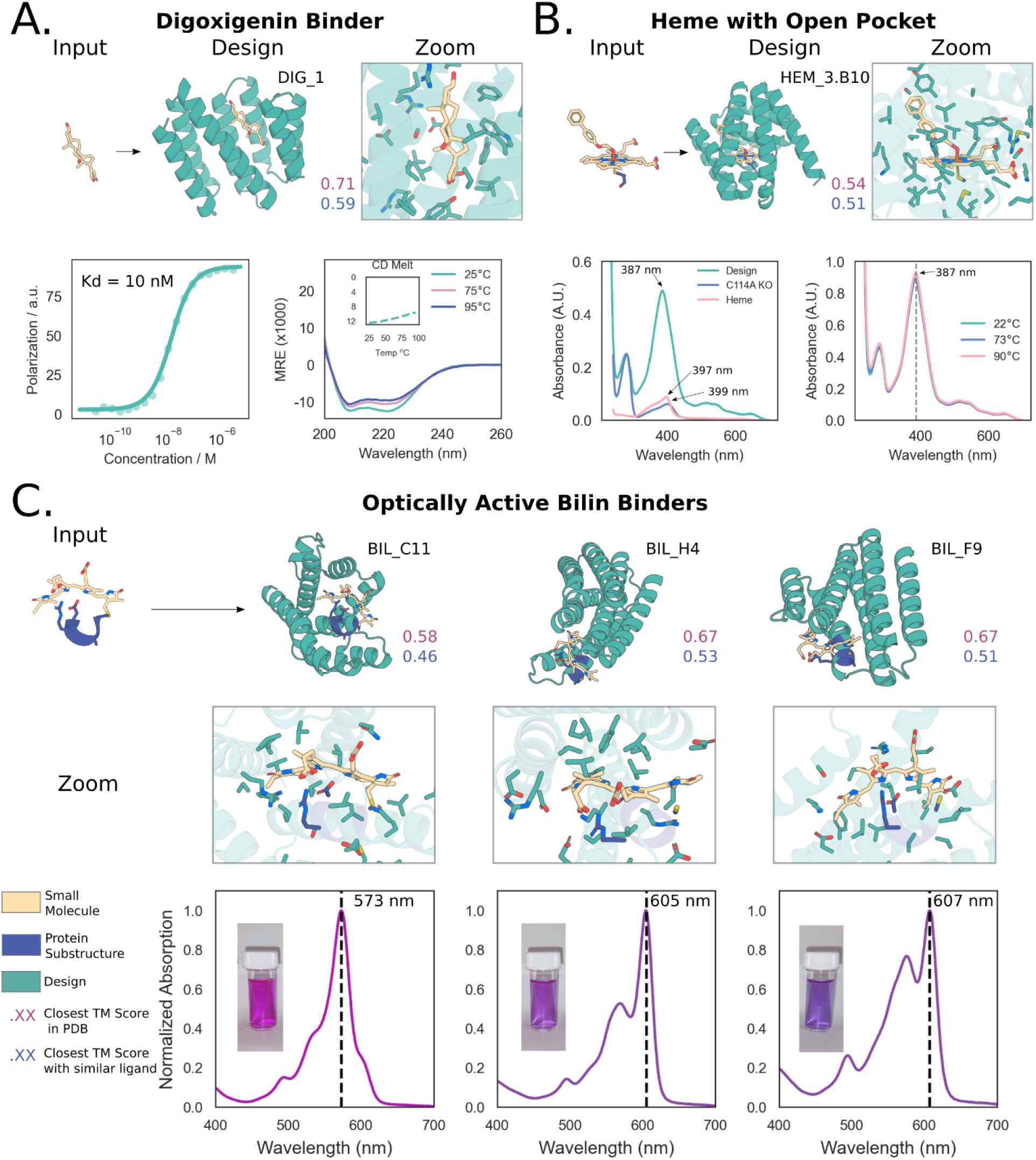
Experimental characterization of RFdiffusionAA designed binders. All panels: input ligand shown in yellow, input protein motif shown in blue, and diffused protein shown in teal. Purple text: Closest TM Score to any protein in the training set, Blue text: Closest TM Score to any protein with a similar ligand bound in the training set (Tanimoto >0.5). **A)** Characterization of Dioxigenin binder design. *Row 1*: (From left to right) Input motif to RFdiffusionAA, designed protein, zoom in view of binding site sidechains. *Row 2*: Fluorescence Polarization (FP) measuring binding affinity (*K*_d_ = 10nM), CD trace (26uM protein concentration; inlay CD Melt showing intensity at 220 nm across a broad range of temperatures). **B)** Characterization of Heme binding designs. *Row 1*: (From left to right) Input motif to RFdiffusionAA, designed protein, zoom in view of binding site. *Row 2*: UV-Vis spectra of designed protein matches expected spectra for penta-coordinated heme and mutating cysteine to alanine abolishes binding, designed protein retains heme binding at temperatures up to 90°C. **C)** Characterization of bilin binding designs. (*Row 1*, left to right) Input motif to RFdiffusionAA, three designs with different predicted structural topologies. (*Row 2*, left to right) Zoom in view of binding site for each design (*Row 3*, left to right) Normalized absorption spectra for the three designs shown. Designs have a range of maximum absorption wavelengths and hence different colors in solution (inset).

Heme is a versatile enzymatic cofactor for a wide range of oxidation reactions and oxygen transport (cytochrome P450 and hemoglobin are two notable examples), with catalytic function enabled by pentacoordinate iron binding and an open substrate pocket (*51*, *52*) Designed heme-binding proteins with these features have considerable potential as a platform for the development of new enzymes (*53*). We diffused proteins around heme with the central iron coordinated by a cysteine and placeholder molecule just above the porphyrin ring to keep the axial heme binding site open for potential substrate molecules. Of 168 designs selected based on AF2 predicted confidence (pLDDT), backbone RMSD to design, and RMSD of the predicted cysteine rotamer to the design, 135 were well expressed in *E. coli*, and 96 had UV/Vis spectra consistent with CYS-bound heme (as judged by the Soret maximum wavelength)(*54*). We further purified 45 of the designs, and found that 38 were monomeric and retained heme-binding through size exclusion chromatography (SEC). Mutating the putative heme-coordinating cysteine residue to alanine led to a notable change in the Soret features upon *in vitro* heme-loading (Figure 5B). The majority of the designs exhibit remarkable thermostability, retaining their heme binding at temperatures above 85 ℃, and do not unfold at temperatures up to 95 ℃ (Figure 5B and Supplementary Information Figures 9-11).

Bilins are brilliantly colored pigments that play important roles across diverse biological kingdoms. When bilins are constrained by protein scaffolds, such as phycobiliproteins in the megadalton phycobilisome antenna complexes of cyanobacteria and some algae(*55*), their absorption features narrow, their extinction coefficients increase, and their fluorescence is dramatically enhanced. We utilized the CARD motif recognised by the CpcEF bilin lyase for covalent attachment of bilins to the native cyanobacterial CpcA phycobiliprotein (*56*), (*57*). Starting from a panel of 94 RFdiffusionAA designs and using phycoerythrobilin (PEB) as the chromophore, we identified nine bilin-binders based on visible pigmentation or fluorescence in a whole cell screen (a 9.6% hit rate) that have quite diverse structures from each other and to CpcA (Figure S8A). From the initial round of screening, we purified three promising designs (C11, H4, and F9) for biochemical and spectroscopic analysis. For each variant, we observed significant spectral shifts (absorption maxima at 557, 605, and 607 nm) relative to the CpcA-PEB control (absorption maxima 573 nm) (Figure 5C, S8B). The degree of red shifting largely correlates with the strength of negative coulombic potential within 5 Å of the chromophore (Figure S9). The three designs have fluorescence quantum yield (FΦ) values of 17%, 38%, and 57% relative to CpcA-PEB, which we arbitrarily set as 100% (Figure S8C), much higher than obtained previously with expert designed scaffolds (maquette proteins) with the CARD motif (FΦ values of 2-3%), which displayed limited bilin incorporation and less pronounced spectral enhancements(*58*). Bilins with enhanced conformational restrictions typically display higher fluorescence yields, and the predicted bilin-binding pocket of H4 imposes the least conformational restriction on the bilin, whereas the C11 binding site restricts the conformation of the bilin to the greatest extent of the three designs.

The 50/46 nm range in absorption/emission covered by just one design round utilizing a single chromophore raises the exciting prospect of tailoring the spectral profiles of designed biliproteins by manipulating the conformational flexibility of the bilin and the protein microenvironment. This could enable building novel antenna complexes to capture light over a wider range of the UV-visible spectrum to enhance photosynthetic energy capture and conversion (*59*), and the design of new fluorescent reporter probes with defined excitation/emission maxima.

The experimental validation of digoxigenin, heme and bilin binders demonstrates that RFdiffusionAA can readily generate novel proteins with custom binding pockets for diverse small molecules that bind their target molecules. Unlike prior methods which rely on redesigning existing scaffolds, RFdiffusionAA builds proteins from scratch around the target compound, resulting in highly shape-complementary binding pockets and reducing the need for expert knowledge. The ability of RFdiffusionAA to generalize is highlighted by the sequence and structural dissimilarity between the designs and proteins in the PDB that bind related molecules (related meaning Tanimoto similarity > 0.5); the most similar protein in the PDB that binds a related molecule has a TMscore of 0.59 for the highest affinity digoxigenin binder, less than 0.62 for all the characterized heme binders, and less than 0.52 for the bilin binders (Figure S10). In all cases there is no detectable sequence similarity to any known protein.

## Discussion

RoseTTAFold All-Atom (RFAA) demonstrates that a single neural network can be trained to accurately model a wide range of general biomolecular assemblies containing a wide diversity of non-protein components. RFAA can make high-accuracy predictions on protein-small molecule complexes, with 32% of CAMEO targets predicted under 2Å RMSD, and of covalent modifications to proteins, predicting 46% of recently solved covalent modifications under 2.5Å RMSD. RFAA goes beyond any previous method we are aware of in generating accurate models for complexes of proteins with two or more non-protein molecules (small molecules, metals, nucleic acids, etc.). These new capabilities do not come at the expense of performance on the classic protein structure prediction problem: RFAA achieves similar protein structure prediction accuracy as AF2 (median GDT of 85 vs. 86) and protein-nucleic acid complex accuracy as RFNA (median allatom-LDDT of 0.74 vs. 0.78) (Figure S11).

Several observations suggest that RFAA has learned general principles about protein-small molecule complexes and not simply memorized the training set. First, the network is able to make high-accuracy predictions for proteins and ligands that differ considerably from those in the training dataset (Figure 2F). Second, prediction accuracy is higher for more tightly binding ligands (Figure 2G) suggesting the network has learned aspects of the physical chemistry of protein-small molecule interactions. Third, our RFdiffusionAA generated bilin, heme, and digoxigenin binders are very different from proteins which bind these compounds in the PDB.

While immediately useful for protein-small molecule binder design and for modeling complex biomolecular assemblies for which there are few or no alternative methods available, the accuracy of RFAA will need to be further increased to have a big impact on drug discovery. The primary factor limiting accuracy is the relatively small size of the training set; whereas there are over 21,000 distinct protein-only structure clusters in the PDB, there are only 6,016 distinct sequence clusters with protein-small molecule complexes. It is likely, however, that a considerably larger number of protein-small molecule crystal structures have been solved in industry but are not publicly available–if it were possible to access such structures to train more accurate and robust versions of RFAA (and other similar networks) considerable public good in the form of improved medicines could result.

## Acknowledgments

We thank Luki Goldschmidt and Kandise VanWormer for maintaining the computational and wet lab resources at the Institute for Protein Design. We thank Joe Watson and David Juergens for helpful conversations during the development of the method and providing the figure color scheme used in this work, Guangfeng Zhou for help with GALigandDock and Madison Kennedy for reading earlier drafts of the manuscript. We also thank Xavier Robin for helping set up our CAMEO server and providing the Vina and AD4 servers for us to benchmark our results and Martin Buttenschoen for providing metadata from their PoseBusters benchmark results.

## Funding

We thank Microsoft for generous donation of Azure Compute Credits, and Perlmutter grant NERSC award BER-ERCAP0022018 for access to the Perlmutter high performance computing resources. This work was supported by gifts from Microsoft (R.K., P.S., D.B.), the Howard Hughes Medical Institute (D.B),, the New Faculty Startup Fund from Seoul National University (M.B.), the Schmidt Futures program (J.W., R.M., F.D.), the Open Philanthropy Project Improving Protein Design Fund (J.W., I.K, G.R.L.), grant no. INV-010680 from the Bill and Melinda Gates Foundation (W.A.), the Audacious Project (P.V.), the Washington State General Operating Fund supporting the Institute for Protein Design (P.V.), Defense Threat Reduction Agency (DTRA) (G.R.L.), Faculty of Science PhD studentship from the University of Sheffield (F.S.M-B), Federal funds from the National Institute of Allergy and Infectious Diseases, National Institutes of Health, Department of Health and Human Services, under Contract No.: 75N93022C00036 (I.A.), Amgen (I.R.H.), Juvenile Diabetes Research Foundation International (JDRF) grant # 2-SRA-2018-605-Q-R (X.L.), Helmsley Charitable Trust Type 1 Diabetes (T1D) Program Grant # 2019PG-T1D026 (X.L.), Defense Threat Reduction Agency Grant HDTRA1-19-1-0003 (X.L.), Bill and Melinda Gates Foundation #OPP1156262 (X.L.), Synergy award 854126 from the European Research Council (G.A.S., C.N.H), Royal Society University Research Fellowship (award no. URF\R1\191548) (A.H.), Human frontiers science program grant RGP0061/2019 (F.D.).

## Author Contributions

Designed the research: R.K., I.A., M.B., F.D. and D.B. Developed RFAA architecture and training regimen: R.K., J.W. and F.D. Evaluated RFAA on different structure prediction tasks: P.S., R.K., I.R.H. and R.M. Developed RFdiffusionAA: W.A. Generated designs for digoxigenin binders: P.V and G.R.L. Performed experiments for digoxigenin binders: P.V, G.R.L, D.V. and X.L. Generated designs and performed experiments for heme binders: I.K. Generated designs for bilin binders: W.A. Performed experiments for bilin binders: F.S.M-B. Contributed code and ideas: I.A., G.A.S., M.B. and F.D. Offered supervision throughout the project: D.B., A.H. and C.N.H. Wrote the manuscript: R.K, J.W., W.A and D.B. All authors read and contributed to the manuscript.

---

**Figure S1.**
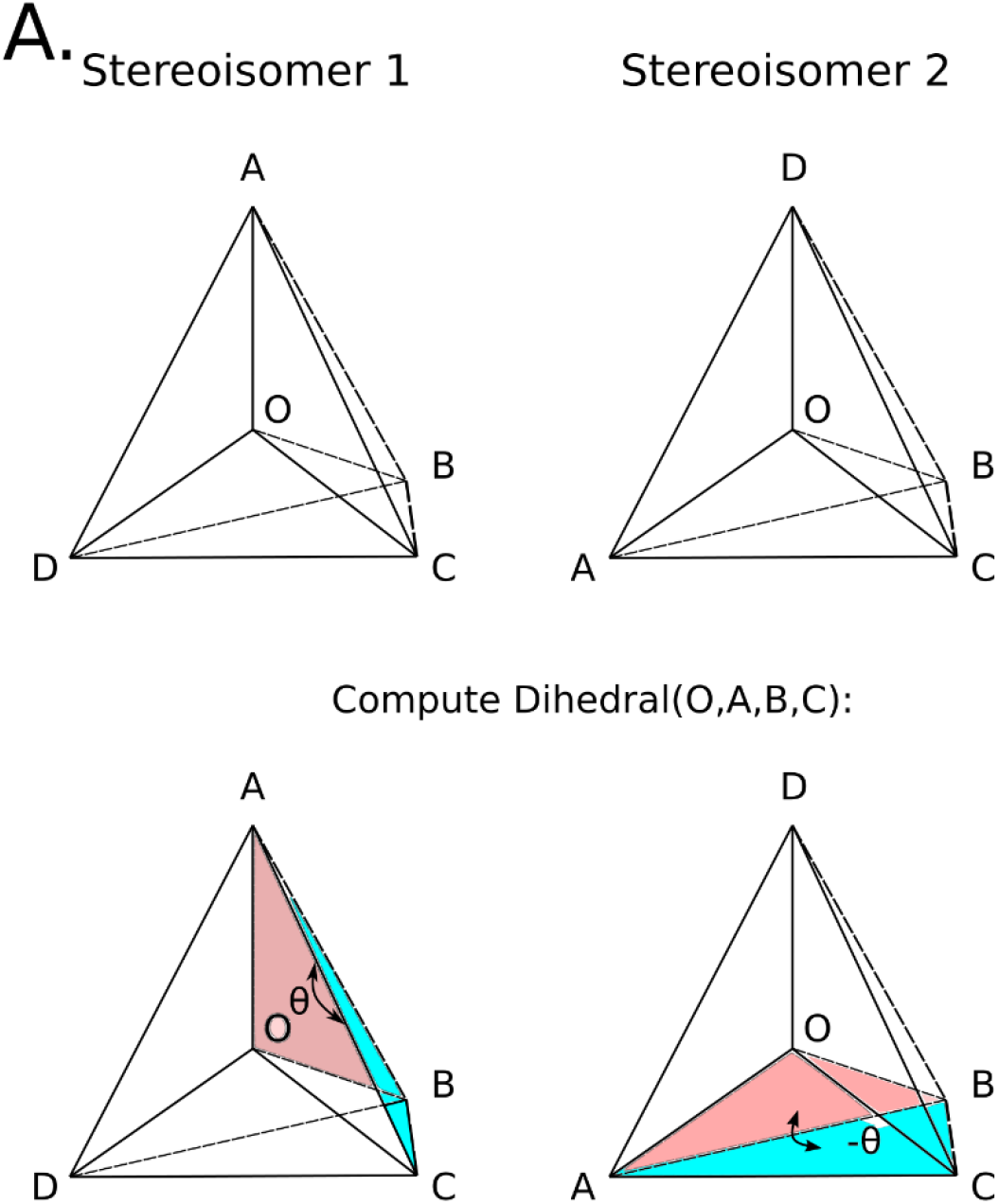
Depiction of chirality input angles. **A)** Chirality is encoded in the network by computing a set of dihedral angles of planes where the first plane starts with the chiral center. *First row:* 3D depiction of a chiral center with tetrahedral geometry with two different chiralities. *Second row:* Depiction of which angles are computed for each center. In practice, we compute the angles for all unique pairs of planes in the center that are explicitly modeled (hydrogens are implicit), measure their error from the ideal tetrahedron in the unit sphere, and pass the gradients of the error in predicted angles with respect to the predicted coordinates into the subsequent blocks as vector input features in the SE(3)-Transformer which breaks the symmetry over reflections present in the rest of the network and allows the network to iteratively refine predictions to match ideal tetrahedral geometry.

**Figure S2.**
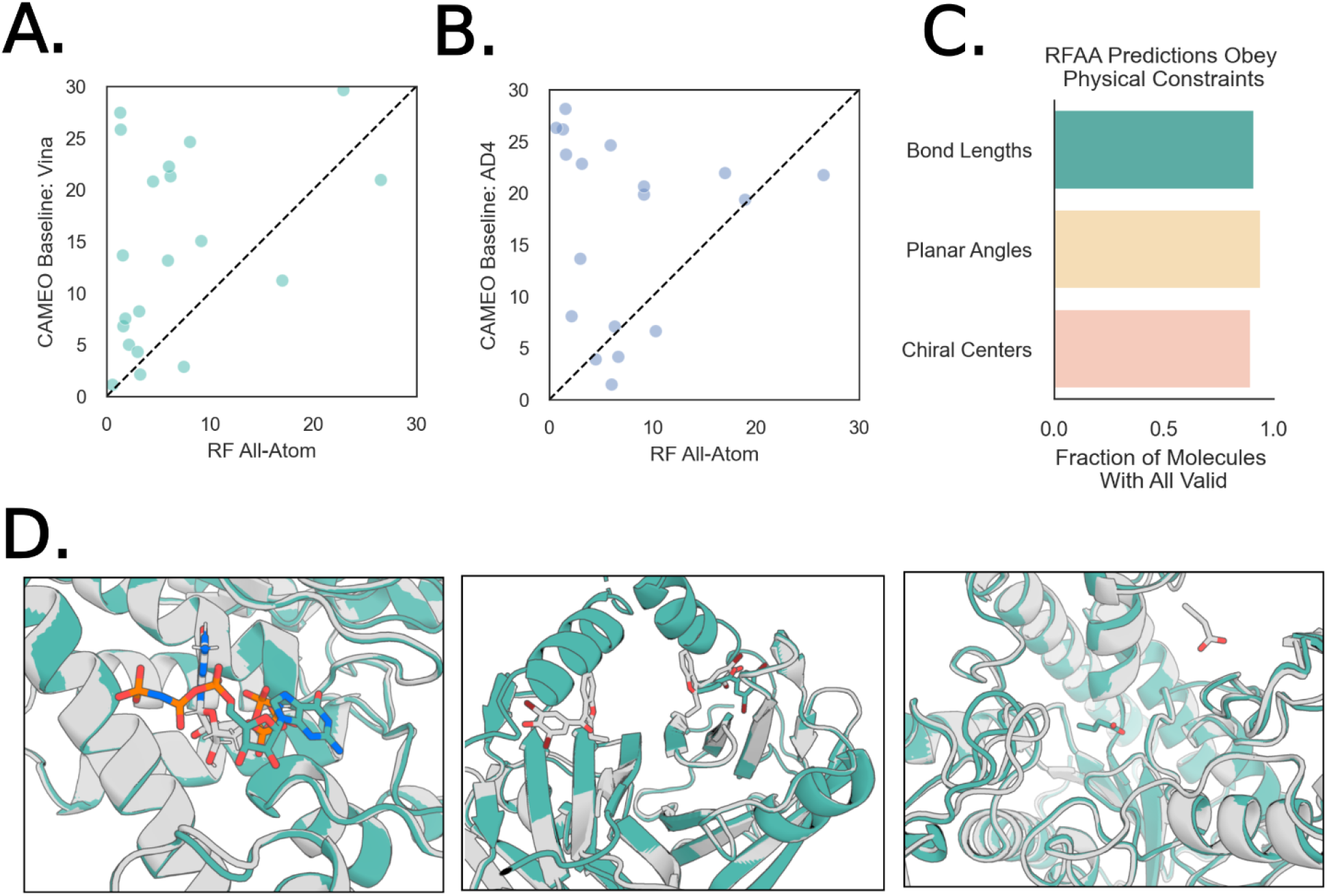
Additional results from the CAMEO ligand-docking challenge. **A)** Pointwise comparison of targets predicted both by RFAA and the CAMEO’s baseline “Vina” server. RFAA predicts lower RMSD docks for a substantial number of ligand-protein complexes. **B)** The same comparison against CAMEO’s “AD4” server. Again, RFAA outperforms the baseline server on the majority of the targets. **C)** RFAA preserves important structural properties of ligands in its predicted poses, such as accurate bond lengths between bonded atoms, planarity of aromatic rings and direction of chiral centers. **D)** Native structures are shown in gray and predicted structures are shown in teal. Some examples of high RMSD poses predicted by RFAA in the CAMEO challenge. From left to right: 1) The model predicts the correct global dock but orients the model incorrectly within the pocket (PDB ID: 7xql). 2) The model predicts an unresolved region as forming a pocket that interferes with the crystal dock of the ligand (PDB ID: 8ii2). 3) The model fails to predict the correct binding pocket of a small ligand, preferring to bury it deep into the protein (PDB ID: 8hwp).

**Figure S3.**
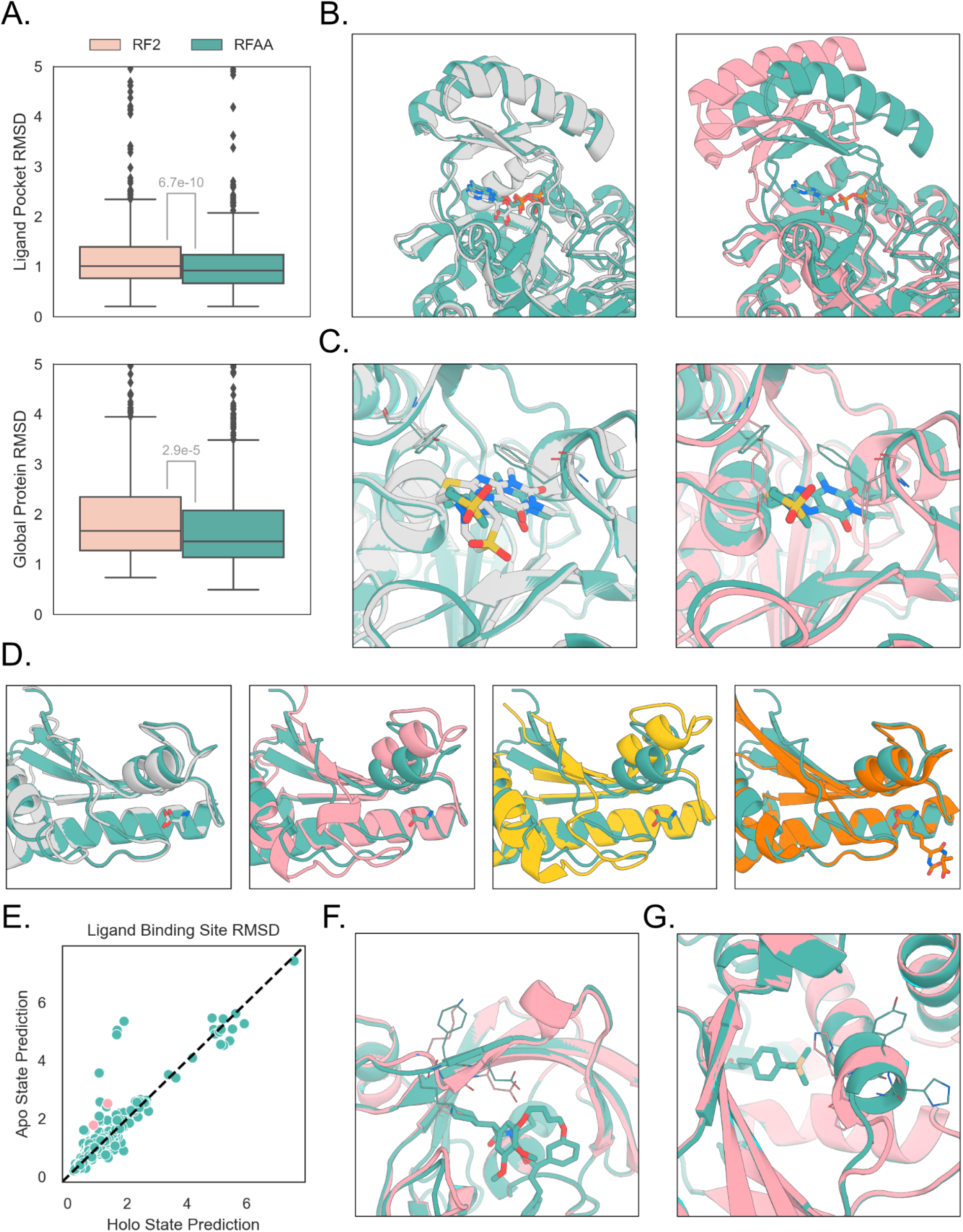
Comparisons to RoseTTAFold2 on predicting protein structures of ligand binding proteins. **A)** Both RFAA and RF2 were used to make predictions on a held-out set of ligand-binding proteins in the PDB (filtered by PAE < 10 and PLDDT > 0.8, respectively). RFAA makes better predictions than RF2, globally and on the pocket residues that bind to the ligand in the crystal structure (p-value < 0.05 paired t-test). **B)** Ligand-aware protein folding allows better positioning of relative protein domains. The crystal structure is gray, the RFAA prediction is green, and the RF2 prediction is pink (PDB ID: 7kct). **C)** Ligand-aware protein folding enables more accurate side-chain predictions in a binding pocket (PDB ID: 7kg7). **D)** Structure prediction for the PDB entry 7rjj, along with the closest sequence match in the model’s training set (yellow, PDB ID 5m38, seq ID 73.1%) and the sequence match in the model’s training set with the most similar bound ligand (orange, PDB ID 5U1H, seq ID 40.2%). RFAA can use information from less similar proteins seen in training to make accurate predictions with small molecule context. **E)** Binding site RMSD of apo and holo predictions of protein chains relative to the holo state crystal structure of the protein. The pink points are depicted in panels F and G. **F)** Prediction of the FK1 domain of FKBP51 with (teal) and without (pink) the binding partner 35-(E) (PDB entry 7b9z). The model demonstrates an understanding of the conformational shift upon binding in the β_3a_ loop, represented in the PDB and our training set (e.g., see PDB entries 3o5e, 6saf). **G)** Prediction of a tRNA ligase (PDB entry 7ckg) with and without binding partner TCQ. The model shifts residues in a neighboring helix to better form a binding pocket for the ligand.

**Figure S4.**
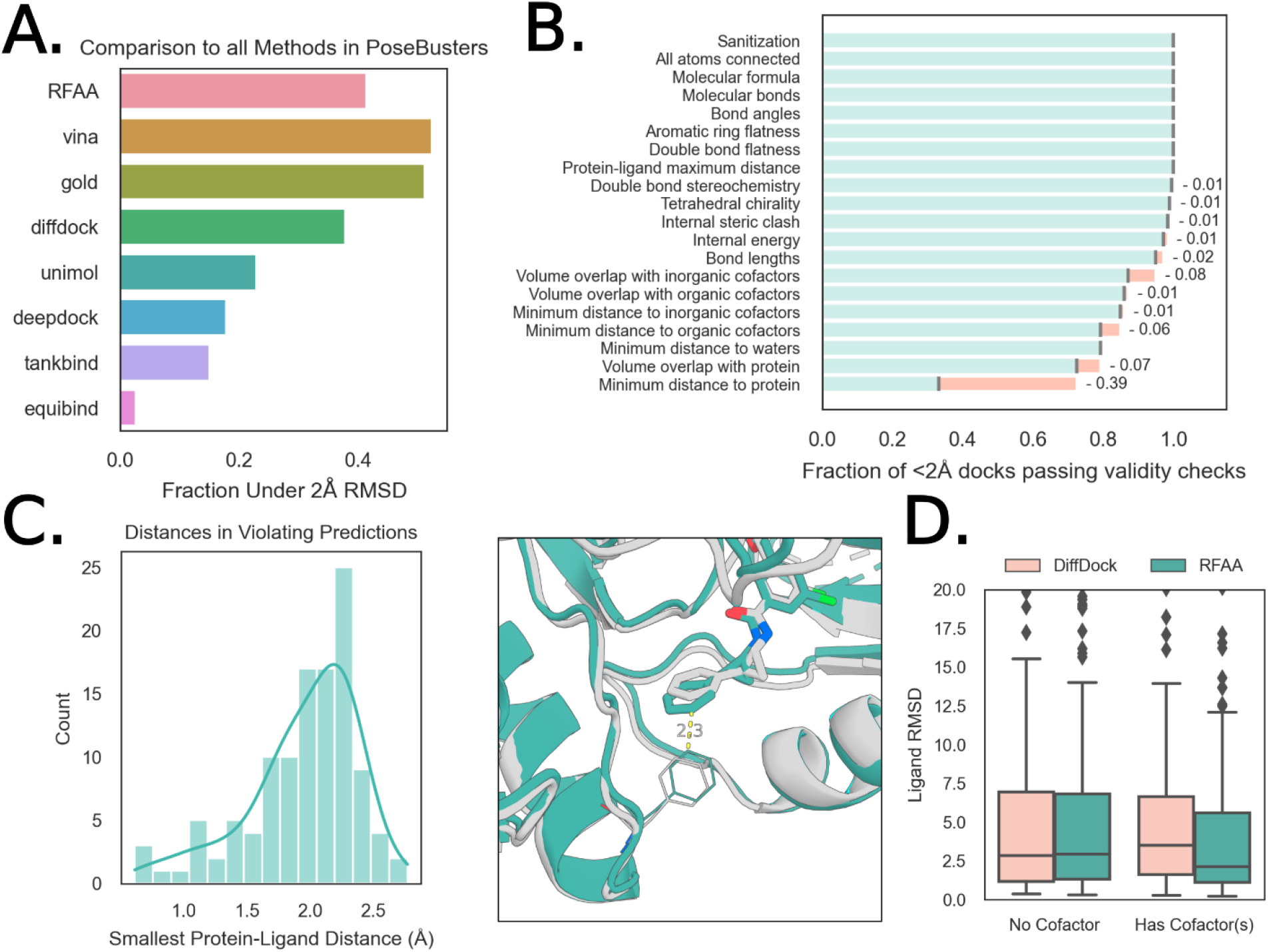
Analysis of predictions of recent PDB protein-small molecule complexes. **A)** Comparison to other methods on PoseBusters benchmark set, which attempts to assess the validity of predicted structures from deep learning networks. Each model is used in its “training regime,” which means that Vina and Gold receive the bound crystal structure and a bounding box for the pocket. RFAA only receives the protein sequence and basic atomic graph information for the small molecule. **B)** Waterfall plot showing results from running the PoseBusters filters on predicted structures from RFAA. Most structures are physically plausible, with most violations occurring in the “minimum distance to protein” metric. **C)** Distribution of shortest distance to protein in violating predictions. Most “violating” distances are between 2.0 and 2.5 Å. The second panel shows an example of a violating distance at 2.3Å. **D)** Comparison of RFAA and DiffDock on cases with and without cofactors. In cases with cofactors, RFAA outperforms because of its ability to simultaneously model multiple molecules.

**Figure S5.**
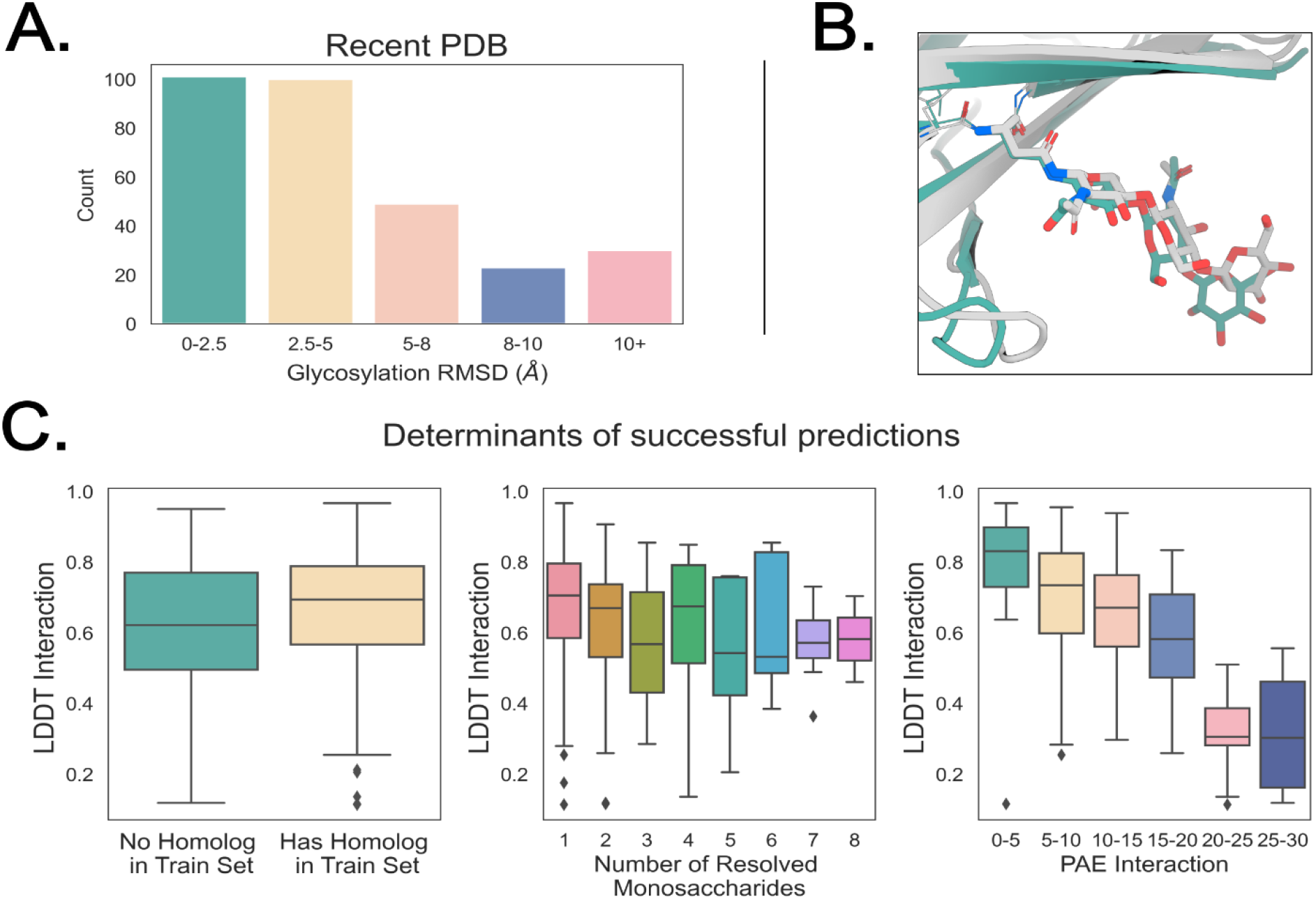
Analysis of predictions of glycosylated proteins. **A)** Histogram of RMSDs of predictions made on recently solved glycoproteins. **B)** Example of successfully predicted glycoprotein (PDB ID: 7u7n), the IL-27 quaternary signaling complex (Native structure in gray, predicted structure in teal). **C)** Determinants of successful predictions from left to right. LDDT Interaction is computed by measuring the all-atom LDDT between protein residues and the residue and modification atoms. From left to right: Having a homolog in the training set (>30% sequence similarity) does not seem to be a large indicator of successful predictions. While predictions of shorter glycans are more accurate, longer glycans can still be successfully modeled. The model’s error prediction accurately identifies high-accuracy predictions.

**Figure S6.**
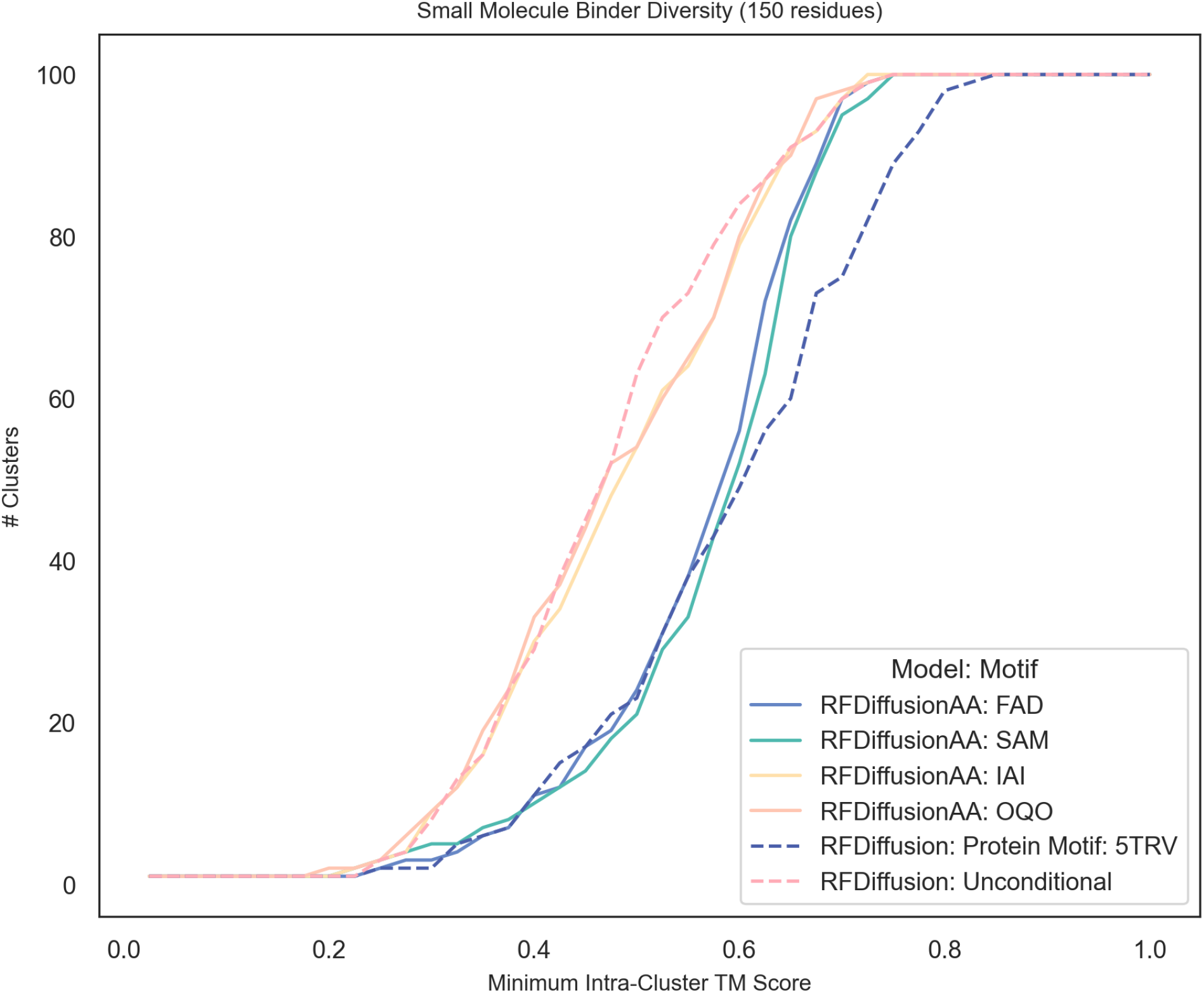
Diversity of small molecule binders generated by RFdiffusionAA. To assess diversity among the generated binders for a given task we perform an all-by-all TMAlign of 100 unfiltered designs. We then perform agglomerative clustering at TM-score thresholds from 0 to 1, such that every design within a cluster has a TM-score to every other design in that cluster less than that threshold and report the number of clusters. Left shifted curves correspond to more clusters at less stringent clustering criteria and thus more diverse designs. We observe that for ligands that appear frequently in the training set [FAD, SAM], the binders generated are less diverse than those designed against ligands with low similarity to any ligand in the training set [IAI, OQO] as the network has come to recognize some of the canonical binding modes of common ligands. To contextualize the magnitude of this difference in diversity we show the same analysis using RFdiffusion when generating unconditional samples vs. RFdiffusion generating motif-conditional samples. RFAA designs have comparable diversity to conditional samples from RFdiffusion.

**Figure S7.**
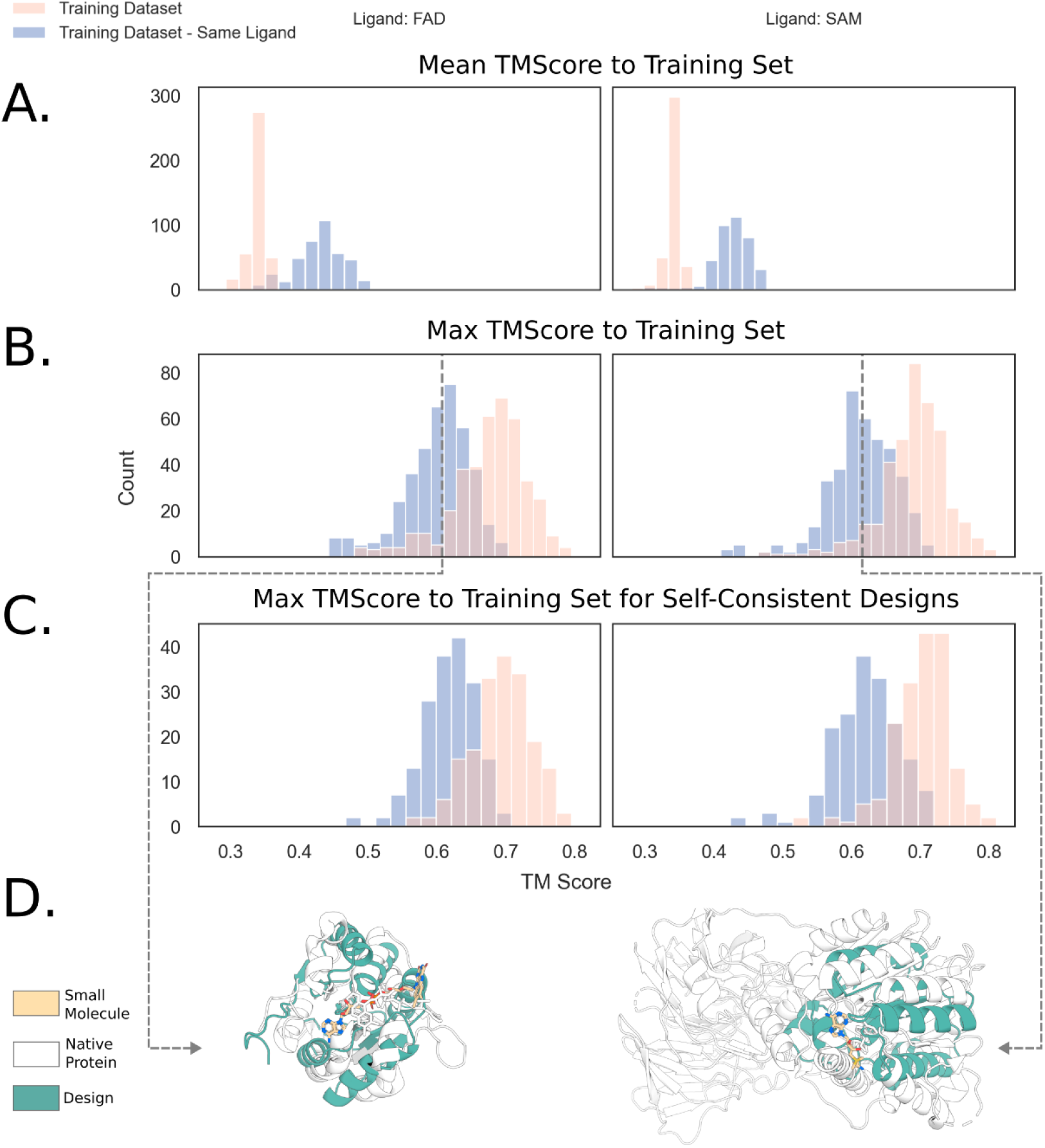
Novelty of binders generated by RFdiffusionAA. **A)** Mean TM-score to all hits in the training dataset for each of 400 designs. Designs against an already-seen ligand are more similar on average to proteins in the training dataset that possess the ligand than to the training dataset as a whole. **B)** Maximum TM-score to the training dataset for each of the 400 designs. The most similar protein to a design in the entire training dataset is substantially more similar than any protein in the training dataset which possesses the ligand, i.e. designs made against an already-seen ligand are not memorized examples from the training dataset for that ligand. **C)** Designs filtered to only those that are self consistent (AF2 RMSD < 2Å), to show that the novelty demonstrated in (B) is not owed to backbones that would not fold. **D)** Teal: The design with median TM-score to the training set with the same ligand bound, White: Closest PDB structure with the same ligand bound.

**Figure S8.**
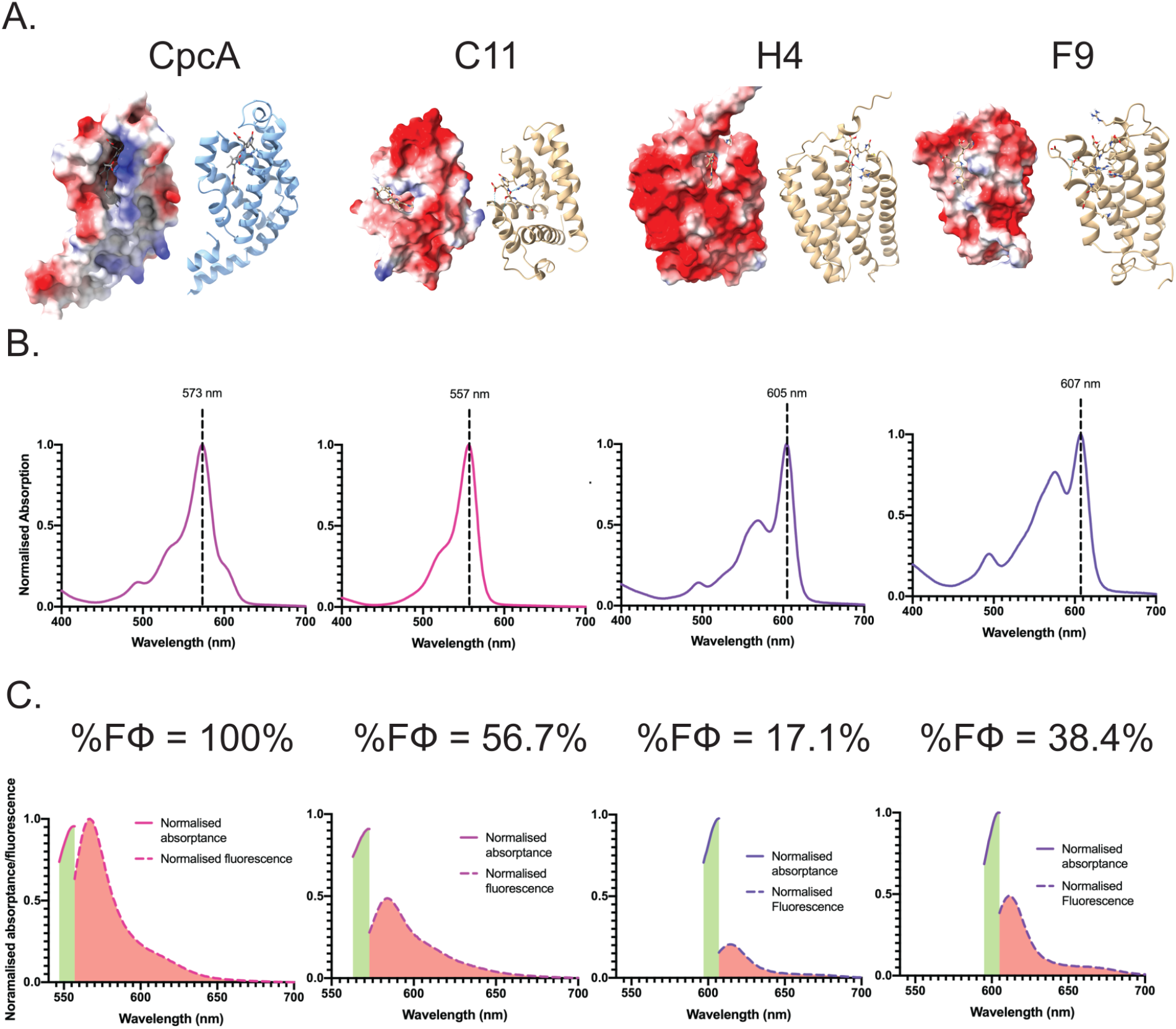
Designed biliproteins. **A)** Molecular topologies and electrostatic space-filling models; molecular topologies of the positive control (CpcA) and three RFdiffusionAA designs (C11, F9, and H4) and their electrostatic space-filling models (blue = positive; red = negative). **B)** Normalised absorption spectra of CpcA, C11, F9, and H4 with images of the colored purified protein solutions inset. **C)** Normalised absorptance and emission profiles of CpcA (100%) and the designed proteins (C11, H4, and F9 - 57.6%, 17.1%, and 38.4%, respectively) - red area under the graph, and the region excited during experiments, expressed as absorptance - green area under the graph.

**Figure S9.**
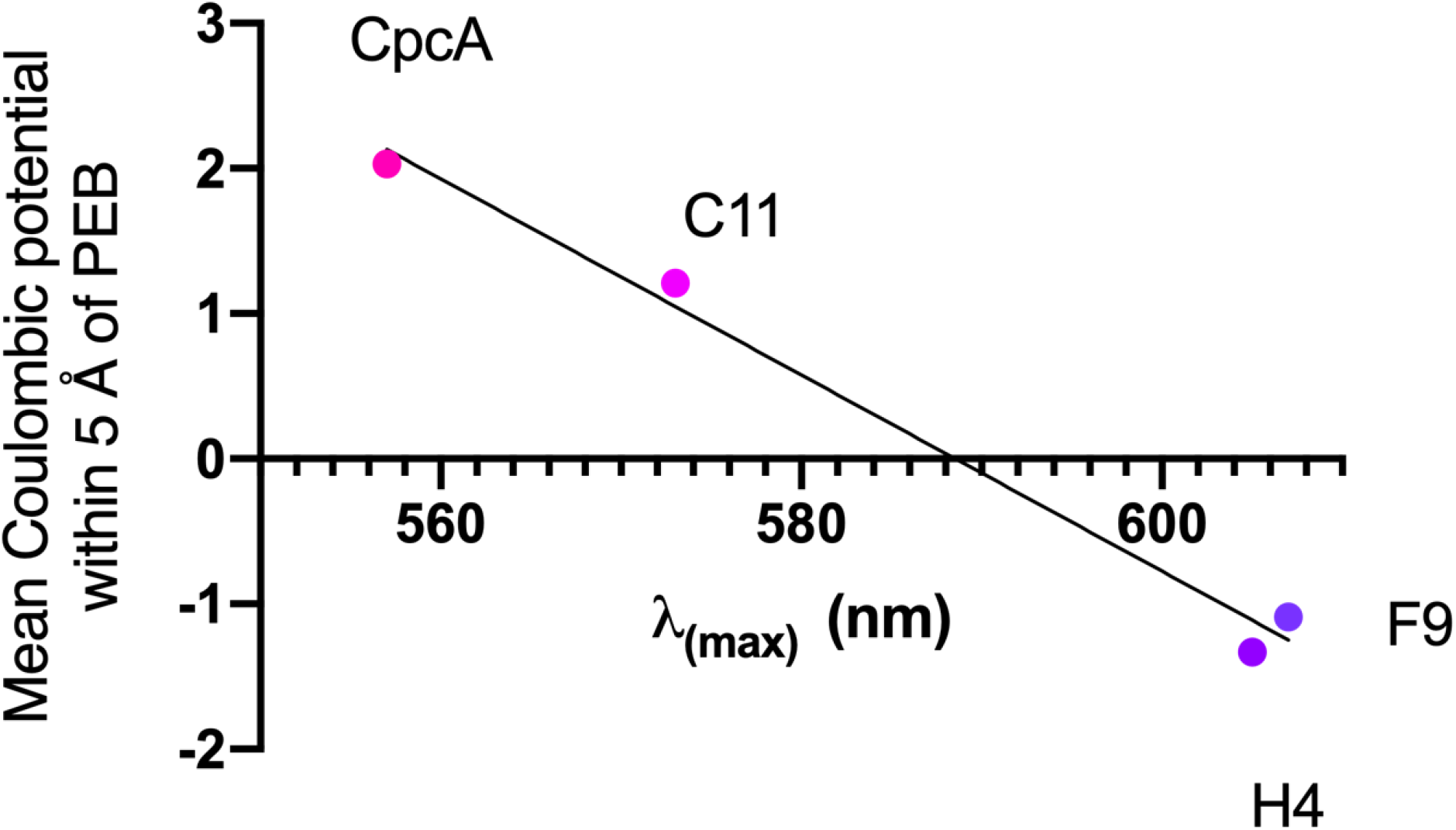
Correlation between coulombic charge surrounding bilin and the maximal absorption wavelength.

**Figure S10:**
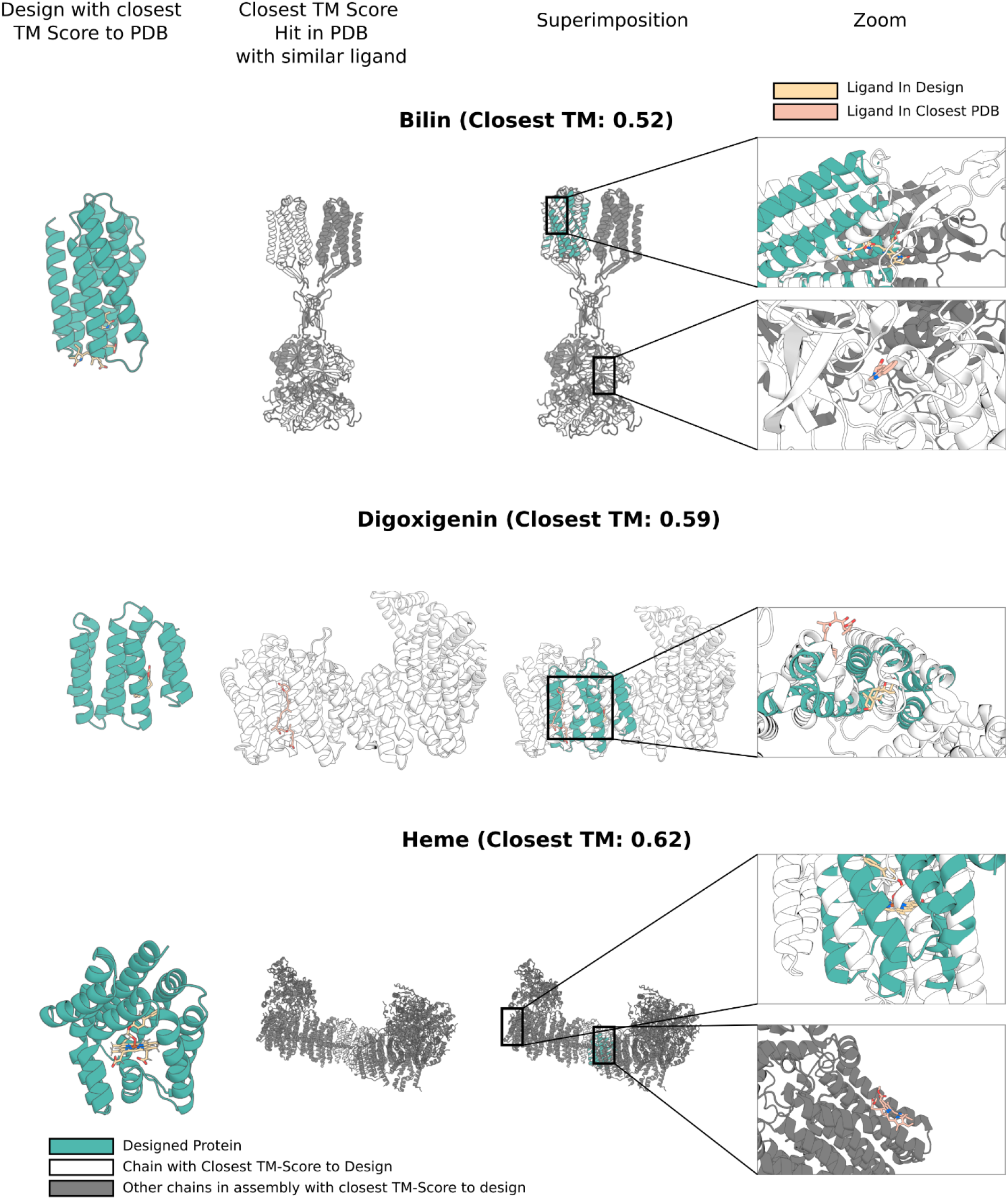
Novelty of experimentally characterized proteins. For each design case, we found the experimentally successful design with the highest TM score to a protein chain in the training set. For all hits in the training set, we measure whether a similar ligand (Tanimoto > 0.5) is present in the PDB entry containing the TM align hit. For each design task, we show the designed protein (teal), the entry with the closest TM score and bound to a similar ligand, the superimposition of the design into the closest TMalign hit and finally a zoom into the binding site of the designed interface (ligand in yellow) and interface with the similar ligand in the training set (ligand in pink). In all cases, the closest entry with a similar ligand in the training set by TM score is below 0.62 and the binding mode of the similar ligand is quite different compared to the binding mode for the designed protein.

**Figure S11.**
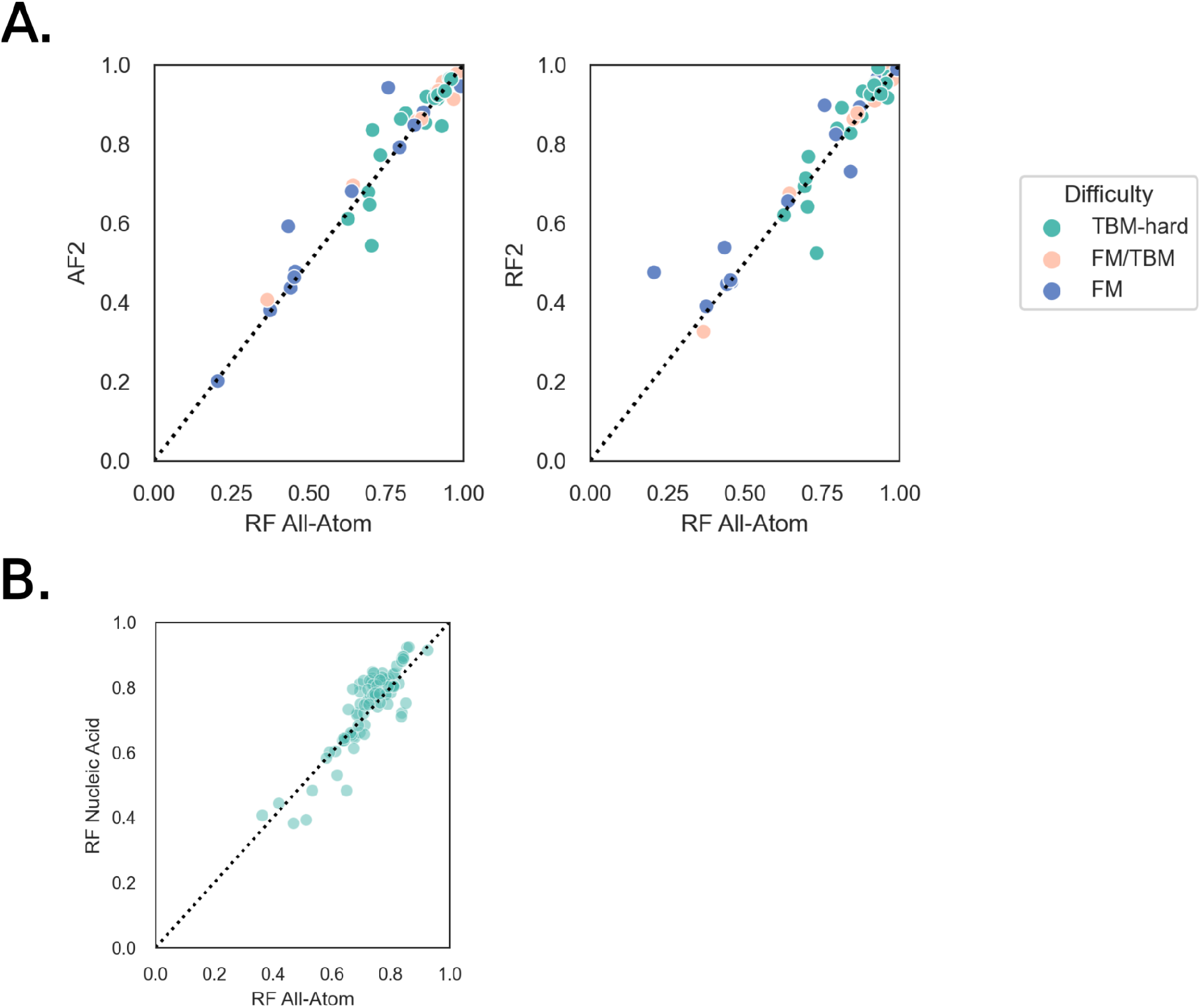
Comparisons to AlphaFold2 and RoseTTAFoldNA on protein structure prediction and protein-nucleic acid interface prediction. **A)** Comparison of RFAA to AF2 and RF2 on protein monomer structure prediction based on a set of 126 structures from CASP14 (TBM-hard, FM/TBM, and FM). Each prediction is scored on GDT to the native structure and colored by difficulty. **B)** Comparison of RFAA to RFNA on protein-nucleic acid complex prediction on a dataset of 89 recently solved structures that were not in the training set of either method.

## Notes

### Competing Interest Statement

The authors have declared no competing interest.

