## Supplemental Methods for "Generalized Biomolecular Modeling and Design with RoseTTAFold All-Atom"

October 9, 2023

### 1 Dataset Curation

The datasets used to train RosettaFold All-Atom can be broadly grouped into three categories: protein-only, nucleic acid, and small molecule datasets. In this section, we describe the curation of each dataset. During training, protein sequence clusters are sampled within each dataset (further discussion of how often each dataset is sampled in Section 6). Multiple Sequence Alignments (MSAs) and templates were generated as in RoseTTAFold2 (RF2) [1]. For MSAs, hhblits [2] was ran at successive e-value cutoffs until 10000 unique sequences with >50% coverage were found. For templates, HHsearch [3] was ran to find a maximum of 500 templates with probability >5%.

#### 1.1 Protein-Only Datasets

Datasets with protein-only examples come from both the Protein Data Bank (PDB) [4] and the AlphaFold2 (AF2) [5] distillation set.

##### 1.1.1 PDB Monomers and Protein Complexes

Similar to RF2 [1], we train on protein monomer and protein complexes structures deposited into the PDB before April 30, 2020 with resolution below 4.5Å. For each chain, we find all contacting chains in all bioassemblies and featurize pairs of homomeric and heteromeric proteins by cropping around the interface. In cases where heteromeric complexes are from the same organism, we provide paired MSAs.

##### 1.1.2 AlphaFold 2 Distillation Data

We train RFAA on a set of UniRef50 structures predicted by AlphaFold2 in [6]. We follow RF2 and augment our training data with structures with AF2 mean pLDDT >70. We apply backbone losses for all structures but only apply sidechain losses for residues with AF2 pLDDT >90. The structures were predicted with only MSAs and no templates and we do not use templates for these examples in training.

#### 1.2 Nucleic Acid Datasets

We follow RoseTTAFold nucleic acid (RF-NA) [7] train on protein-nucleic acid complexes and RNA structures. The training setup is identical to the training set for RF-NA.

#### 1.3 Small Molecule Datasets

This section deals with data that involves non-polymer small molecules (or non-linear polymers such as sugars), generally bound to a protein. Each item in the various small molecule datasets is represented by a selected subset of chains from a PDB entry. We build the dataset by iterating through each entry in the PDB with a bound, non-polymer chain. For each non-polymer chain, we first discard that entry if it is considered "non-biological" (see Section 1.3.1). Otherwise, we treat that chain as a *query ligand* of interest and compute a list of every chain in that entry that contacts it (see Section 1.3.2). We maintain a running list of transformations operate on each chain's coordinates to place it into a global reference frame: this is necessary for symmetric assemblies in the PDB where a single chain may represent multiple copies of identical molecules in different, symmetric positions. In all cases, hydrogens are not modeled. Finally, we compute MSAs and templates for each protein chain contacting each query ligand in the same manner as in Section 1.1.

This definition of dataset item implies that a single PDB entry can comprise of multiple items in the dataset corresponding to different query ligands. For example, the PDB entry '5nag' corresponds to two entries in our dataset: one for each bound ligand (FAD and 8R5). At training time, we crop each entry around the query ligand (further description in Section 3.3), resulting in different contexts for the two query ligands.

We also note here that we only include in our small molecule datasets those PDB entries that are present in either the PDBBind [8] or BioLip [9] datasets.

##### 1.3.1 Ligand Filtering

We find that many of the non-polymer entities in PDB entries are *non-biological*: that is, they represent solvents or crystallization additives rather than binding partners found in a biological context. We curated a list of 3-letter component identifiers (which have up to 3 letters) from the PDB corresponding to such non-biological molecules based on the BioLip database [9] and our own manual inspection as shown in Table 1. For example, GOL is the three letter code for glycerol, a common solvent.

We remove all non-biological from our training set as they generally represent non-specific binding partners and/or molecules held in place by a crystal lattice rather than by protein interactions. We note that the list in Table 1 may not be exhaustive, but it filters out the most common examples we observed in the PDB.

##### 1.3.2 Contacting Chains

A protein chain is considered "in contact" with the query ligand if at least 5 C$\alpha$ atoms of that protein chain are within 30Å of any atom in the query ligand or if any atom in the protein chain is within 5Å of any atom in the query ligand. This definition of contacting protein chain allows for a broad biological context in which to predict the query ligand of interest.

A non-polymer chain is considered "in contact" with the query ligand if at least one atom of that chain is within 5Å of the query ligand, or if all atoms are within 30Å of the query ligand. The model thus learns

Supplementary Methods Table 1: Nonbiological Molecules

| |
|---|
| NUC, ZN, CA, MG, III, MN, FE, CU, SF4, FE2, CO, FES, GOL, NA, CL, K, CU1, GOL, XE, NO2, EDO, NI, BR, CD, O, CS, NO, TL, HG, UNL, KR, SR, RB, F, AG, AR, U, AU, MO, SE, GD, YB, VX, SM, LI, RE, N, W, OS, HO, PI, EDO, PG4, OGA, SO4, HEZ, FEO, CL, DMS, ACT, MPD, GOL, NH2, CUA, SIW, PGW, IOD, BR, 3NI, ZRW, 78M, UNX, MES, CCN, PO4 |

Supplementary Methods Table 2: Selected Metals

| |
|---|
| LA, NI, 3CO, K, CR, ZN, CD, PD, TB, YT3, OS, EU, NA, RB, W, YB, HO3, CE, MN, TL, LI, MN3, AU3, AU, EU3, AL, 3NI, FE2, PT, FE, CA, AG, CU1, LU, HG, CO, SR, MG, PB, CS, GA, BA, SM, SB, CU, MO, CU2 |

to predict not only the relative positions of a ligand and its protein binding partners, but also associated cofactors in the same or nearby binding pockets.

###### 1.3.3 Covalent Modification Dataset

We separate query ligands that are covalently bonded to a protein into a distinct and separate dataset. For each such query ligand, we filtered out those that involved a covalent bond between an oxygen atom on the ligand and an oxygen atom on the protein, as well as protein-ligand fluorine-flourine bonds, covalent bonds to hydrogen atoms, and all cases where the length of the protein-ligand covalent bond was less than 1Å. We find that such covalent protein-ligand bonds are usually between a protein and a non-biological small molecule, as described in Section 1.3.1.

###### 1.3.4 Metal Ion Dataset

We also separate query atoms that represent metal ions into a distinct dataset. In order to avoid expanding the vocabulary set of our model too greatly and potentially introducing non-specific metal binding sites into the dataset, we only train on the metals whose 3-letter PDB codes are listed in Table 2.

###### 1.3.5 Small Molecule Datasets

Every remaining query ligand can be grouped into one of the following groups: multi-residue, multi-protein assembly, and single-protein assembly. We describe the three sub-datasets here:

1. **Protein/Small Molecule Complex**: Dataset containing all single residue small molecules that only interact with a single other protein chain. However, the query ligand in each item of this dataset lies on a single residue in the PDB entry.
2. **Protein/Multi-Residue Ligand Complex**: Dataset containing all "multi-residue" ligands. Such ligands exist as multiple residues (or a single residue with multiple bioassembly transforms) in their respective cif file, and usually represent sugar chains or small peptides.
3. **Protein/Small Molecule Assembly**: Dataset containing non-protein biomolecular context (small molecules, covalent modifications, multi-residue ligands, metal ions) and $>1$ protein chain. This

| **Dataset** | **Sequence Clusters** | **Examples** |
|---|---|---|
| Protein Monomer | 21,648 | 301,934 |
| AF2 Distillation Set | 1,036,080 | 3,605,951 |
| Protein Heteromer | 13,755 | 183,821 |
| Protein Nucleic Acid | 1,235 | 17,240 |
| RNA | 1,449 | 6,522 |
| Protein Small Molecule | 5,662 | 121,800 |
| Protein Metal Complex | 5324 | 112,456 |
| Protein Multi-Residue Ligand | 613 | 4,775 |
| Protein Small Molecule Assembly | 2,564 | 43,838 |
| Covalent Modification | 1,099 | 12,689 |

Supplementary Methods Table 3: Number of Protein Sequence Clusters In Each PDB Training Dataset

| **Dataset** | **Sequence Clusters** | **Examples** |
|---|---|---|
| Protein Monomer | 1,206 | 47,094 |
| Protein Homomer | 682 | 25,284 |
| Protein Heteromer | 1792 | 24,454 |
| Protein Nucleic Acid | 79 | 1,272 |
| RNA | 153 | 720 |
| Protein Small Molecule | 354 | 18,110 |
| Protein Multi-Residue Ligand | 51 | 302 |
| Protein Small Molecule Assembly | 163 | 6,165 |
| Covalent Modification | 80 | 1,834 |

Supplementary Methods Table 4: Number of Protein Sequence Clusters In Each PDB Validation Dataset

dataset is distinct from 1. because we generate paired MSAs when there are two or more distinct (heteromeric) protein chains in a complex.

###### 1.3.6 Dataset Clustering

We cluster each entry in each dataset by their *primary protein partner*. For protein-only datasets, this is either the protein monomer, or in the case of hetero-oligomers, we arbitrarily select the first chain that appears in the PDB entry as the primary chain. For nucleic acid and small molecule datasets, we designate as the primary protein partner the protein chain with the greatest number of atoms within 5Å of the query nucleic acid/ligand. All items in each database are then clustered using MMSeqs2 [10] using the default hyper-parameters and subsequently used for dataset sampling at training time.

###### 1.3.7 Final Statistics

After all the filters were applied, we kept chains that have resolution $< 4.5$Å and that were deposited in the PDB before April, 30th, 2020 for the purposes of held-out evaluation on PDB entries from 2021 onward. 10% of protein sequence clusters were held out of training for validation during training. The final number of items and clusters in each dataset is shown in Table 3. The number of validation items and clusters in each dataset is shown in Table 4.

##### 1.4 CSD Dataset

In addition to molecules in the PDB, we augment our training dataset with small molecule crystal structures from the Cambridge Structure Database (CSD v5.43; November 2021) [11]. We filter structures based on the following metrics: 1) resolution $< 5$Å, 2) not polymeric, 3) greater than 5 atoms resolved, 4) less than 100 atoms resolved, and 5) ability to be parsed by OpenBabel [12]. We sample molecules with equal probability and separate a validation set that does not have Tanimoto score $>0.75$ to any molecule in training.

#### 1.5 Negative Datasets

Following RF2 and RF2NA, we use a set of "negative" interactions to help the network focus on relevant features that constitute binding. Briefly, for proteins this means showing examples where we randomly pair chains and only assess a loss on each individual chain and for nucleic acids it involves mutating bases that make essential contacts for the formation of the complex.

Supplementary Methods Table 5: Element Tokens in RFAA

| |
|---|
| Al, As, Au, B, Be, Br, C, Ca, Cl, Co, Cr, Cu, F, Fe, Hg, I, Ir, K, Li, Mg, Mn, Mo, N, Ni, O, Os, P, Pb, Pd, Pr, Pt, Re, Rh, Ru, S, Sb, Se, Si, Sn, Tb, Te, U, W, V, Y, Zn |

### 2 Modeling Arbitrary Biological Inputs

Architectures for modeling both protein structures and nucleic acid structures have been previously described. In this section, we describe the necessary changes to such architectures to model small molecules, covalent modifications to protein structures and arbitrary non-canonical amino acids.

#### 2.1 Expanded Input Sets

The most significant architectural change from existing protein structural networks is the expanded input features that RFAA takes in, which we describe in the following sections.

##### 2.1.1 New Tokens

The original RoseTTAFold architecture had 22 tokens: 20 amino acids, 1 unknown and 1 gap token. The RF-Nucleic Acid expanded this token set by 10, adding 8 distinct tokens for the 4 DNA and 4 RNA bases, and 2 tokens for unknown DNA base and unknown RNA base, respectively. The RFAA architecture includes 46 additional tokens representing individual atoms with the element types shown in Table 5, a token for deprotonated histidine (unused in practice, left in for legacy reasons) and an unknown atom token for a total token count of 80.

##### 2.1.2 Bond Connectivity

When predicting structures of arbitrary molecules, it is important for the network to know the bond connectivity of those molecules. To provide this information to the network, we pass in the bond connectivity of input molecules as a 2D bond adjacency matrix as an input. We designate 7 bond types representing single bond, double bond, triple bond, aromatic bond, residue-residue (or base-base), residue-ligand atom bond and other bond type. In practice, the "other" bond type is not used but exists for historical reasons. The residue-residue "bond" type exists to be able to provide bond features for protein and nucleic acid inputs, so that the input bond matrix is always of the same dimension as the input. Residue-atom bonds exist in order to do a process we call residue *atomization*, which is used to model ligands that are covalently bonded to a residue and arbitrary non-canonical amino acids. This process is described further in Section 3.4.

##### 2.1.3 Chiral Features

Another key bias in more generalized biomolecular modeling is chirality. Aforementioned features like atom types and bond connectivities are not sufficient to specify the chirality of input molecules, so we provide chiral features explicitly to the network at each chiral center. In this work, we only deal with tetrahedral chirality and leave more complicated forms of stereochemistry to future work.

For each tetrahedral chiral center, we enumerate all sets of three heavy atom neighbors in all orders. For each ordering, we compute the pseudo-dihedral angle between those four points (center and 3 heavy atoms) and note whether that angle is positive or negative, which determines the chirality of the center

Supplementary Methods Table 6: Bond Types in RFAA

| **Bond type** | **Encoding** |
|---|---|
| No bond | 0 |
| Single | 1 |
| Double | 2 |
| Triple | 3 |
| Aromatic | 4 |
| Protein residue-residue or nucleic acid base-base | 5 |
| Residue-atom | 6 |
| Other | 7 |

uniquely. For ideal tetrahedral geometry, the magnitude of the dihedral angle is $\arcsin(1/\sqrt{3})$. See Section 4.5.2 for further details.

We compute the difference between the predicted dihedral angle and the ideal dihedral angle of the chiral center, and pass the coordinate-wise gradients of the difference as an input to the 3D track of the network. This representation of chirality provides direct signal to the network on how to update atomic coordinate positions in order to obey the ideal tretrahedral geometry. This process is described further in Section 4.5.

###### 2.1.4 Atom Frames

Key to the success of AF2 and RF2 is the frame aligned point error loss [5]. This involves aligning N-C$\alpha$-C backbone frames of predicted structures to true structures and then measuring the error of all the other predicted atoms with respect to the true structure in that alignment. This loss has attractive properties for biomolecules such as not being invariant to reflections which allows the network to predict correct chirality. We construct *canonical* frames for each atom in small molecules comprising of atoms and their bonded neighbors. We achieve this by iterating through all bonded triplets of atoms and assigning each triplet a priority based on the bonded atoms, depicted in Table 7. The process for constructing canonical frames from a ligand is outlined below:

1. Construct a graph where each node is an atom and each edge is a bond.
2. For each atom in the graph:
   (a) Enumerate through all paths of length three containing that atom. If there exist paths such that the given atom is in the center, exclude all other paths.
   (b) Compute frame priorities for each atom in each path and make a list of frame priorities in increasing order.
   (c) For each such path of length three, sort them by lexicographic atom frame priority, e.g. for two paths A and B, path A will appear before path B if and only if either the lowest frame priority in path A is less than the lowest frame priority in path B, or they are equal and the second lowest frame priority in path A is less than the second lowest in path B, and so on.
   (d) The first path in the lexicographic order is chosen as the frame for this atom, and the order of the frame is determined by frame priority in increasing order.

This process deterministically computes a local coordinate frame for each atom in arbitrary molecules (with at least 3 atoms). If a frame has an unresolved atom, it still is assigned as the canonical frame but is not used in loss calculation. The usage of the atom frames is further discussed in Sections 4.5, 5.5 and

5.4. Importantly, these features are constructed from the bond graph so they maintain the permutation invariance of the inputs to the network.

| **Element Type** | **Priority** |
|---|---|
| K | 0 |
| Li | 1 |
| Ca | 2 |
| Mg | 3 |
| Be | 4 |
| Y | 5 |
| Tb | 6 |
| U | 7 |
| V | 8 |
| W | 9 |
| Mo | 10 |
| Cr | 11 |
| Re | 12 |
| Mn | 13 |
| Os | 14 |
| Ru | 15 |
| Fe | 16 |
| Pr | 17 |
| Ir | 18 |
| Rh | 19 |
| Co | 20 |
| Pt | 21 |
| Pd | 22 |
| Ni | 23 |
| Au | 24 |
| Cu | 25 |
| Hg | 26 |
| Zn | 27 |
| Al | 28 |
| B | 29 |
| Pb | 30 |
| Sn | 31 |
| Si | 32 |
| C | 33 |
| Sb | 34 |
| As | 35 |
| P | 36 |
| N | 37 |
| Te | 38 |
| Se | 39 |
| S | 40 |

| | |
|---|---|
| O | 41 |
| I | 42 |
| Br | 43 |
| Cl | 44 |
| F | 45 |

Supplementary Methods Table 7: Frame Priorities of Atoms in RFAA

###### 2.1.5 Positional Encodings

To break permutation symmetry for sequences, we use a signed relative positional encoding for protein sequences and nucleic acid bases [5]. Atomic graphs require permutation symmetry so we do not provide a relative positional encoding. For atomic inputs, we provide a separate embedding that measures the shortest distance between any pair of atoms in the bond graph described in Section 2.1.2. We develop a generalization of these two embeddings for cases where atom nodes are bonded to residues to encode the distance between an atom and its closest bonded residue. Further details are provided in Section 4.1.

### 3 Data Pipeline

Our data pipeline involves taking raw data from cif files and formatting the data into input tensors for the network. We first will describe the inputs to the network and then go through details of how each dataset is preprocessed. We use OpenBabel to parse the ideal sdf files for each PDB but do not use any chemical quantities computed by it (just element types, bond types and whether an atom is chiral center). We find this to be preferable because OpenBabel can process all ideal sdf files provided by the PDB so we do not exclude examples because of parsing errors. We compute the direction of the chiral center as discussed in Section 4.5.2.

#### 3.1 Inputs for RFAA

Remaining features such as MSAs and templates are handled identically for proteins to RF2. The coordinate dimension, 36, reflects the maximum amount of heavy atoms and hydrogens possible in a residue or base. The small molecule tokens are appended to the first sequence in all the MSA features and the remaining MSA sequences are initialized with gap tokens. Small molecules receive empty template features which are concatenated block diagonally to the protein features. A detailed description of the inputs are shown in Table 8.

| **Input (dimension)** | **Description** |
|---|---|
| *msa_masked* ($N_{num\_clusters}$, L, 164) | Clustered MSA with some portions of the sequences masked. For atom nodes, the first sequence has its respective atom tokens and then remaining sequences are filled with gap tokens. (80 raw msa, 80 cluster statistics, 2 insertions/deletions, 2 Nterm/Cterm) |
| *msa_full* ($N_{num\_sequences}$, L, 80) | Full MSA clipped at 1024 sequences. |
| *seq* (L, 80) | First row of the MSA. In this case, the protein sequence and any atom tokens, including mask tokens. |

| | |
|---|---|
| *idx*<br>(L) | Residue index of each residue in the input. This input must be provided for atom nodes but has no semantic meaning (it is unused by the network). |
| *bond_feats*<br>(L, L, 7) | Pairwise bond adjacency matrix. Pairs of residues are either single, double, triple, aromatic, residue-residue, residue-atom or other. |
| *dist_matrix*<br>(L, L) | Minimum amount of bonds to traverse between two nodes. This is 0 between all protein nodes. |
| *chirals*<br>($L_{num\_chiral\_centers}$, 5) | All orderings of 4 atoms around a chiral center (first four dimensions) and the ideal pseudo-dihedral angle formed by that ordering of atoms (fifth dimension). |
| *atom_frames*<br>($L_{num\_atoms}$, 3, 2) | Indices that form frames for each atom node in the input. The second dimension represents that there are three atoms in each frame. The third dimension represents an offset in the node dimension because atom frames go across nodes and the absolute index in the atom dimension. |
| *t1d*<br>($N_{num\_templates}$, L, 80) | 1D template feature. First, 79 represent the "sequence" (residue/atom types) of the templated structure. Last dimension represents residue wise template confidence. |
| *t2d*<br>($N_{num\_templates}$, L, L, 64) | 2D template information which gives the binned distances and angles between frames (N-C$\alpha$-C for proteins, designated atom frame for atoms) |
| *alpha_t*<br>($N_{num\_templates}$, L, 30) | Sidechain torsion angles from templates (10 angles x sin, cos and whether the angle exists in the structure for each residue) |
| *msa_prev*<br>($N_{num\_clusters}$, L, $C_m$) | Recycled MSA features. $C_m$=256 (number of 1D channels) |
| *pair_prev*<br>(L, L, $C_p$) | Recycled pair features. $C_p$=192 (number of 2D channels) |
| *state_prev*<br>(L, $C_s$) | Recycled state features. $C_s$=32 (number of 3D $\ell 0$ channels) |
| *xyz_prev*<br>(L, 36, 3) | Recycled XYZ coordinates. On first iteration, this is set to the coordinates from the first template. If no templates, coordinates are initialized at the origin with random noise (between -2.5 and 2.5Å) applied. |
| *sc_torsions_prev*<br>(L, 30) | Recycled predicted sidechain torsion angles. |

Supplementary Methods Table 8: Inputs to RFAA

#### 3.2 Featurization of Symmetric Permutations

When featurizing multimers (empirically in the homomer and small molecule assembly datasets), there are often identical chains that, if swapped, result in the identical complex. We want the gradient of the loss to push the network towards the relabeled complex that is closest to the prediction so during preprocessing we track which chains can be swapped so that we can deconvolute which permutation to apply the loss on during training. We use the same scheme to account for permutation swaps of atoms in small molecule structures.

#### 3.3 Featurization of Protein Small Molecule Complexes

A similar preprocessing procedure was followed for the small molecule protein complex dataset, the multichain residue ligand dataset, the covalent modification dataset and the protein-small molecule assembly dataset (multiple contacting protein chains). Each training example centers around a single query molecule in a specific bioassembly. Based on the details of the bioassembly features for the nearest chains (both protein and other small molecules) are constructed. There are two types of biomolecular contexts that are sampled stochastically. First, metal ions in the presence of other small molecules are sampled stochastically because often is is not a priori known when solving a structure whether there will be a metal crystallized in the pocket. Second, if there modified residues present in the cif file, they are featurized as an atomized residues rather than their canonicalized version.

Due to memory restrictions, we then perform a cropping procedure to select a subset of nodes to represent a training example. The cropping procedure samples a random atom on the query molecule and computes the distance to all other atoms or C$\alpha$ atoms in proteins. It then selects the top n_crop nodes to include in the crop. In our early experiments we found that sometimes, there were protein chains that were either too short or far away from the ligand in euclidean space that were included in the crop with insufficient context to be predicted accurately. In these cases, the gradient was dominated by the incorrect prediction of those protein chains and not on the correct docking of the small molecule. To remedy this, after finding the top n_crop nodes, we iterate through all the protein chains that were in the crop and remove any chains with $<$10 contacts to other nodes in the crop or with less than 8 residues. After these chains are removed, we noticed that certain molecules in the crop also did not have sufficient contacts to be docked so we iterate through all the molecules in the crop and remove any molecules that have $<$4 contacts to a protein chain. The exact logic is shown in 1.

It is evident from this cropping process that full subunits in large symmetric assemblies could be cropped out. Since we compute potential symmetric relabeling of chains before cropping (see Section 3.2), certain chains that were computed as potential symmetric relabelings are no longer valid (specifically because the small molecule context should drive the network to predict a specific interface when a symmetric oligomer could have multiple distinct protein-protein interfaces). After cropping, we reiterate through the precomputed symmetric permutations and remove those that are no longer possible given the chains that were removed during cropping.

#### 3.4 Featurization of Atomized Protein Examples

Training examples for *atomized* proteins first are featurized identically to protein monomer examples (except the stochastic homomer featurization is turned off, see Section 6). After cropping, a number of residues is sampled from Uniform(3,5) and that number of (fully resolved) contiguous residues is chosen for atomization. If there are not enough valid residues in the crop to *atomize*, we treat the example as a monomer example. We then take that selection of residues, featurize them using all the small molecule features (atom tokens, bond features, chirality inputs). We also provide bond tokens to indicate bonds between the first N token in the *atomized* region to the previous residue and the last C token to the following residue. Finally, the MSA and template information for these residues is removed from the input features so the network must learn how to generate their structures and poses from the atomic

**Algorithm 1** Cropping for SM Complex Datasets

**function** CROP_SM_COMPL($xyz, query\_mol, n_{crop}$)
  $atom_{query} \leftarrow Uniform(atoms_{query\_mol})$
  $d = ||xyz_{atom_{query}} - xyz_j||$ ▷ distances between query atom and all C$\alpha$s
  keep $\leftarrow n\_lowest\_values(n = n_{crop}, d)$
  **for** chain=0... $N_{sm\_chains}$ **do**
    **if** any(keep *in* chain) **then**
      keep += all(atoms *in* chain) ▷ do not crop ligands
    **end if**
  **end for**
  keep_atoms $\leftarrow$ atoms *in* keep
  **for** chain=0... $N_{protein\_chains}$ **do**
    keep_chain $\leftarrow$ keep *in* chain
    $d_{ca} = ||xyz_{keep\_chain} - xyz_{keep\_atoms}||$
    **if** Count(keep_chain) $< 8$ OR Count($d_{ca} < 4$) $< 10$ **then**
      keep -= keep_chain ▷ remove chains with few contacts to small molecule
    **end if**
  **end for**
  keep_residues $\leftarrow$ residues *in* keep
  **for** chain=0... $N_{sm\_chains}$ **do**
    keep_atom_chain $\leftarrow$ keep *in* chain
    $d_{ar} = ||xyz_{keep\_atom\_chain} - xyz_{keep\_residues}||$
    **if** Count($d_{ar}$ <4) $< 4$ **then**
      keep -= keep_atom_chain ▷ remove small molecules with no contacts to proteins
    **end if**
  **end for**
  **return** keep
**end function**

information. Practically, we precompute the atoms, bonds and chiralities of each atom in each residue and convert the features as shown in Algorithm 2. Symmetric swaps of sidechain atoms are accounted for in the same manner as Section 3.2.

**Algorithm 2** *Atomization* of Protein Residues

**function** ATOMIZE_PROTEIN(residue_indices, seq, *protein_features)
  atoms $\leftarrow$ CONCAT(atoms_in_residues(seq[residue_indices]))
  bonds $\leftarrow$ BLOCK_DIAGONAL(bonds_in_residues(seq[residue_indices]))
  bonds += atomized_peptide_bonds ▷ add peptide bonds between adjacent atomized residues
  chirals $\leftarrow$ chirals_per_residue(seq[residue_indices])
  frames = get_atom_frames(atoms, bonds)
  DELETE *protein_features[residue_indices]
**end function**

#### 3.5 Featurization of Covalently Bound Ligands

Covalently bound ligands are preprocessed very similarly to small molecule complexes. The covalent modifications are modelled just as other ligands would be. The residue that has the covalent bond to the ligand is atomized using the same method as Section 3.4. The atom in the ligand and the atom in the *atomized* residue is provided in the bond features (eg. single bond between atom i from modification and atom j in atomized residue). All other featurization (protein MSA, templates etc) remains the same as other datasets.

#### 3.6 Featurization of Metal Ions

Metal Ions are provided to the network as a single atom ligand. The only difference is that since metal ions only have a single atom, they do not have their own canonical frame. In these cases, the network does not receive a frame input and there is no loss calculated with respect to the frame of the ion (there are still gradients from the error of the placement of the ion with respect to the other frames in the structure).

#### 3.7 Featurization of CSD Small Molecule Crystals

Asymmetric units of crystal lattices from the CSD are featurized identically to small molecules that are bound to proteins. The network is then tasked with predicting the atomic coordinates of the molecule. Molecules with less than 5 atoms, greater than 100 atoms, polymers or resolution $>5$Å are discarded from the dataset. Remaining molecules that could be parsed by OpenBabel were used to train the network.

### 4 Algorithm Details

The architecture of RFAA is similar to RF2. The network accepts 1D, 2D and 3D inputs and treats structure prediction as a graph inference problem where the objective is to find edges in the graph where edges are distances in euclidean space. The outputs of the model are predicted coordinates, confidence measures and the latent embeddings from the three tracks. There are 3 main stages of the network, the embedding stage, the simulator stage and the refinement layers. Similar to AF2 and RF2, the network employs recycling where latent features (shown in Algorithm 3) from previous forward passes to the network are provided to future iterations. Here, we will mainly cover the places where our implementation diverges from RF2. Importantly, the RFAA architecture has three main features that are passed throughout the network, *msa*, *pair* and *state*. *msa* are features from the 1D track, *pair* are features from the 2D track and *state* are node-wise features from the 3D track. The outputs of the network include *xyz* and *sc_torsions* which are the predicted backbone coordinates and predicted side chain torsion angles respectively (which can be used to construct the full atom coordinates). For clarity, intermediate outputs of the network are in italics and preprocessed features are in plaintext.

Similar to RF2, the first four blocks process the *msa_full* features. In these layers, the network uses

---

**Algorithm 3** Recycling Iterations

msa, pair, state = **0**, **0**, **0**
xyz, sc_torsions = Uniform(-2.5,2.5), **0** ▷ or template structure if available
**for** i=0... $N_{cycle}$ **do**
    *msa, pair, state, xyz, sc_torsions* = RFAA(*input_features, *msa, pair, state, xyz, sc_torsions*)
**end for**

---

global column attention over the full MSA instead of column attention and biased row attention used in the 36 main blocks. This portion of the network is meant to extract extra information for cases with deep MSAs. Similar to all blocks in RFAA, there are predicted structures generated for each block and the features from those bias the future interactions of the network. The implementation of these are unchanged from RF2 (except the additional atomic context is also processed), the intuition being that these layers are mainly for MSA processing that is not applicable to atomic graphs.

The next 32 layers are the main blocks of the network. They process the *msa_masked* features along with the biomolecular context. The main block logic is shown in Algorithm 5. Each block in RFAA has five steps, first the computation of the structure bias from the previous block, second the *msa* feature update, third the update of the *pair* features based on the *msa* features, fourth the update of the *pair* features and finally the update of the 3D features. The full algorithm is shown in Algorithm 4, which has 83M parameters that are optimized through training.

#### 4.1 1D Embeddings

The 1D embeddings are very similar to those in RF2. For the *msa_full* blocks, they are identical. The msa_full and seq input features are embedded and the embedding of the *seq* features are added to every row of the *msa* features. The *msa_masked* embedding slightly diverges from RF2 (6). The *msa_masked* embedding begins with the exact same operations as the *msa_full* embedding that construct the initial *msa* features. Then, the initial *pair* and *state* features are constructed. The *pair* features are constructed by embedding the *seq* input with two sets of weights and then computing the outer product of the two embeddings.

At this point the relative positional encoding is added (in the *msa_full* blocks the attentions are column-wise so breaking row-wise symmetry is less important). Since atoms nodes in small molecules are permu-

**Algorithm 4** RFAA Forward Pass

**function** RFAA(msa_masked, msa_full, seq, idx, bond_feats, dist_matrix, chirals, atom_frames, t1d, t2d, alpha_t, msa_prev, pair_prev, state_prev, xyz_prev, sc_torsions_prev, $N_{full\_blocks}$=4, $N_{main\_blocks}$=32, $N_{ref\_blocks}$=4)
    $msa_{full}$ = MSA_Full_Embed(msa_full)
    $msa_{cluster}, pair, state$ = MSA_Cluster_Embed(msa_masked, seq, idx, bond_feats, dist_matrix)
    $pair$ += Bond_Embed(bond_feats)
    $msa_{cluster0}, pair, state$ += Recycle_Embed(msa_prev, pair_prev, state_prev, xyz_prev, sc_torsions) ▷ Only update first row of MSA representation
    $pair, state$ += Template_Embed(t1d, t2d, alpha_t, *pair*, *state*)
    **for** i=0... $N_{full\_blocks}$ **do**
        Stopgrad(xyz)
        $msa_{full}, pair, state, xyz, sc\_torsions$ = Full_Block(msa_full, *pair*, *state*, *xyz*)
    **end for**
    **for** i=0... $N_{main\_blocks}$ **do**
        Stopgrad(xyz)
        chiral_grads = calc_chiral_grads(*xyz*, chirals)
        $msa_{cluster}, pair, state, xyz, sc\_torsions$ = Main_Block($msa_{cluster}, pair, state, xyz$, idx, bond_feats, dist_matrix, chiral_grads, atom_frames)
    **end for**
    **for** i=0... $N_{ref\_blocks}$ **do**
        Stopgrad(xyz)
        chiral_grads = calc_chiral_grads(*xyz*, chirals)
        clash_l1_grads, clash_l0_grads = calc_clash_grads(*xyz*, *sc_torsions*)
        $state, xyz, sc\_torsions$ = Ref_Block($msa_{cluster}, pair, state, xyz$, idx, bond_feats, dist_matrix, chiral_grads, clash_l1_grads, atom_frames, clash_l0_grads)
    **end for**
    **return** $msa_{cluster}, pair, state, xyz, sc\_torsions$
**end function**

**Algorithm 5** RFAA Main Block

**function** MAIN_BLOCK(*msa_cluster*, *pair*, *state*, *xyz*, idx, bond_feats, dist_matrix)
    $str\_bias$ = RBF($||xyz_{C\alpha} - xyz_{C\alpha}||$)
    rel_pos, bond_dist = Positional_Encoding(idx, bond_feats, dist_matrix)
    $str\_bias$ += Linear(rel_pos) + Linear(bond_dist)
    $msa_{cluster}$ = 1D_Update($msa_{cluster}$, *pair*, *state*, *str_bias*)
    *pair* = Aggregation(*msa_cluster*, *pair*)
    *pair* = 2D_Update(*pair*, *str_bias*, *state*)
    $xyz, state, sc\_torsions$ = 3D_Update($msa_{cluster}$, *pair*, *state*, *xyz*, idx, bond_feats, dist_matrix, atom_frames)
    **return** $msa_{cluster}, pair, state, xyz, sc\_torsions$
**end function**

tation invariant, we remove the positional encoding for these nodes. When considering *atomized* residues which are in the polypeptide chain, we must break permutation symmetry for these atoms. We intended to provide the network both information about where the atoms are in the polypeptide chain and the relative distances within the bond graph between atoms (and between atoms and residues). We devised two encodings, the first is the signed relative position encoding (capped between -32 and 32) similar to RF2 and the second is a relative bond graph distance (capped at 8). For each atom in the bond graph we compute the shortest path through the bond graph to all other atoms. For cases where atoms are bonded to residues, we count the atom-residue bond as an additional bond in the shortest distance feature and then provide the closest residue's relative positional offset. Both of these features are linearly embedded and summed (Algorithm 7). Finally, the *state* features are initialized by embedding the *seq* inputs.

**Algorithm 6** Clustered MSA Features Embedding

**function** MSA_CLUSTER_EMBED(msa_masked, seq, idx, bond_feats, dist_matrix)
    $msa_{cluster}$ = Linear(msa_masked)
    $seq$ = Linear(seq)
    $msa_{cluster0}$ += $seq$
    $seq_i, seq_j$ = Linear(seq) ▷ Different weights
    $pair$ = Outer_Product($seq_i, seq_j$)
    rel_pos, bond_dist = Positional_Encoding(seq, idx, bond_feats, dist_matrix)
    $pair$ += Linear(one_hot_encode(rel_pos))
    $pair$ += Linear(one_hot_encode(bond_dist))
    $state$ = Linear(seq)
    **return** $msa_{cluster}, pair, state$
**end function**

**Algorithm 7** Generalized Positional Encoding

**function** POSITIONAL_ENCODING(idx, bond_feats, dist_matrix)
    rel_pos = clip($idx_i - idx_j$, -32, 32)
    bond_dist = clip(dist_matrix, 8)
    **if** atom_residue_bond **then** ▷ Atomized protein or covalent modification
        closest_termini = argmin(dist_matrix[:, terminal_N, terminal_C])
        closest_residues = bonded_residue(bond_feats, closest_termini)
        rel_pos[atomized_nodes] ← rel_pos[closest_residues] ▷ add relative position of closest residue for each atomized nodes
    **end if**
    **return** rel_pos, bond_dist
**end function**

#### 4.2 Bond Embedding

We reasoned that bonds provide very straightforward constraints on pairwise distances between atoms. We linearly embed the one-hot encoded bond graph and sum it with the *pair* features to allow the network to predict accurate bond lengths and angles.

**Algorithm 8** Bond Embedding

**function** BOND_EMB(bond_feats)
    $bond\_emb$ = Linear(one_hot_encode(bond_feats))
    **return** $bond\_emb$
**end function**

##### 4.3 1D Track Update

The 1D track update is identical to the 1D track update in RF2. The *pair* features are passed through a LayerNorm and then an embedding of the radial basis function (64 bins, 0.5 Å increments, standard deviation=0.5) of the distances from the predicted coordinates is added to *pair* features. A LayerNorm is applied to the *state* features and then it is embedded. The first row in the *msa* embedding (representing the query sequence embedding) is updated with the embedded state features. The *msa* embedding is then updated with row attention with bias from the *pair* features and then column-wise attention is computed to update the *msa* features [5, 13, 14].

##### 4.4 2D Track Update

The 2D track update is identical to the update in RF2. The *pair* features are updated using the TriangleMultiplicationOutgoing and TriangleMultiplicationIncoming updates with 0.25 probability of dropout for each. We then apply structure biased axial attention as in RF2. To form the structure bias, the distances between C$\alpha$ (or P for nucleic acids and atom for atom nodes) coordinates predicted at the previous block are binned with gaussian blur using a radial basis function (64 bins, 0.5 Å increments, standard deviation=0.5). This binned distribution is embedded to match the number of channels in the pair dimension. To gate which distances should bias the pair features, the *state* features are embedded with two separate sets of weights and the outer product of the embeddings is computed. The outer product goes through another linear embedding, followed by a *sigmoid* activation function. This final value is used as a gate and multiplied by the binned distance distribution which represents the coordinate bias that will be applied. Row and column attention are computed for the pair feature and the coordinate bias is added to the attention weights. Finally, a FeedForward layer updates the final pair representation from the block.

We did not change the functional form of the 2D update from RF2 but we expected that the bottleneck to learning would be the 2D update which has to learn many new types of interactions about arbitrary biomolecules. We expected that a more rich pair representation and more attention heads would give the network enough capacity to learn the new interactions that were shown in the training set following intuition from [5] and [15]. We increased the number of pair channels to 192 (from 128 in RF2) and the number of attention heads per update to 6 (from 4 in RF2).

##### 4.5 3D Track Update

The 3D track update is where our implementation diverges the most from RF2. The goal of the 3D track update is to use the features from the 1D and 2D tracks and predict the coordinates of the complex. Similar to RF2, we sought to use the SE(3)-Transformer architecture which guarantees equivariance over the group of rotations and translations by projecting features into a basis formed by the spherical harmonics [16, 17]. The SE(3) Transformer is a graph neural network which takes in node features for every node and edge features for each connected pair of nodes and aggregates features across them. This architecture uses tensor products to mix $\ell 0$ (scalar features), $\ell 1$ features (vector features) and higher order $\ell$ features and can predict arbitrary $\ell$ features for each node. In this section, we will describe how we formulated $\ell 0$ and $\ell 1$ features for each node that is modelled and edge features between nodes. The full algorithm is shown in Algorithm 9.

**Algorithm 9** RFAA Structure Update

**function** 3D_UPDATE($msa_{cluster}, pair, state, xyz, idx, bond\_feats, dist\_matrix, chiral\_grads, atom\_frames$)
  $seq$ = LayerNorm($msa_{cluster0}$) ▷ Query sequence latent embedding
  $pair, state$ = LayerNorm($pair$), LayerNorm($state$)
  $node\_feats$ = Linear(CONCAT([$seq, state$]))
  $node\_feats$ += FeedForward($node\_feats$)
  $node\_feats$ = LayerNorm($node\_feats$)
  rel_pos, bond_dist = Positional_Encoding(idx, bond_feats, dist_matrix)
  rbf_feat = rbf($||xyz_{C\alpha} - xyz_{C\alpha}||$) ▷ Or P or atom coordinate
  $edge\_feats$ = Linear(CONCAT([$pair$, rbf_feat, rel_pos, bond_dist]))
  $edge\_feats$ += FeedForward($edge\_feats$)
  $edge\_feats$ = LayerNorm($edge\_feats$)
  $frame_{ij}, frame_{kj} = frame_i - frame_j, frame_k - frame_j$ ▷ Construct frames from N-C$\alpha$-C, O-P-O or atom_frames
  l1_feats = CONCAT([$frame_{ij}, frame_{kj}, chiral\_grads$]
  geometric_feats = $xyz_i - xyz_j$
  $state, coord\_update$ = SE3_Transformer(node_feats, edge_feats, l1_feats, geometric_feats)
  $quaternion\_values, T = coord\_updates$
  $R$ = construct_rotation_matrix_from_quaternion($quaternion\_values$) ▷ Set R = Identity matrix for "atom" nodes
  $xyz = R \circ xyz + T$
  $sc\_torsions$ = SC_pred($msa_{cluster0}, state$)
  **return** state, xyz, sc_torsions
**end function**

##### 4.5.1 Constructing $\ell 0$ features

The first row of the *msa* features (the features corresponding to the query sequence) and the *state* features are passed through a LayerNorm. These values are concatenated and then processed by a shallow network consisting of a Linear embedding, a FeedForward Layer and a LayerNorm. These features are all scalar values and represent the $\ell 0$ node features.

##### 4.5.2 Providing the Network Chirality Inputs

To provide the network chirality features, we aimed to construct a geometric representation of chirality that does not involve learning human made concepts such as (r) and (s) (as the network would need to learn how to implicitly order substituents and determine whether they were presented in the clockwise or counterclockwise order). We observed that set of atoms in chiral centers form planes and the angle between the frames are different depending on the stereochemical identity of that center. We then construct an ideal tetrahedron on the unit sphere with centroid at the origin (where the chiral center would be), which composes of four points:

$$v_1 = (\sqrt{\frac{8}{9}}, 0, -\frac{1}{3})$$
$$v_2 = (-\sqrt{\frac{2}{9}}, \sqrt{\frac{2}{3}}, -\frac{1}{3})$$
$$v_3 = (-\sqrt{\frac{2}{9}}, -\sqrt{\frac{2}{3}}, -\frac{1}{3})$$
$$v_4 = (0, 0, 1)$$

We can then compute the dihedral angle between the planes formed by $(v_1, v_2, v_3)$ and $(v_2, v_3, v_4)$:

$$\begin{aligned}
&\text{Dihedral}(a, b, c, d) = \text{atan2}( \\
&\quad [c-b] \cdot (([b-a] \times [c-b]) \times ([c-b] \times [d-c])), \\
&\quad \|c-b\|([b-a] \times [c-b]) \cdot ([c-b] \times [d-c])) \\
&\text{Dihedral}(v_1, v_2, v_3, v_4) = 1.23 \,\text{radians}
\end{aligned}$$

In the example above, a switch in stereochemistry would result in the following coordinates($v_3$ comes out of the plane and $v_4$ goes into the plane):

$$\begin{aligned}
v_1 &= (\sqrt{\frac{8}{9}}, 0, -\frac{1}{3}) \\
v_2 &= (-\sqrt{\frac{2}{9}}, \sqrt{\frac{2}{3}}, -\frac{1}{3}) \\
v_3 &= (0, 0, 1) \\
v_4 &= (-\sqrt{\frac{2}{9}}, -\sqrt{\frac{2}{3}}, -\frac{1}{3}) \\
\text{Dihedral}(v_1, v_2, v_3, v_4) &= -1.23 \,\text{radians}
\end{aligned}$$

The positions of these four atoms are sufficient to determine chirality. The dihedral angle between planes $(v_1, v_2, v_3)$ and $(v_2, v_3, v_4)$ will be positive in the first case and negative in the second. In practice in RFAA, we do not always have access to all four substituents of a chiral center (one of them could be a hydrogen which is not explicitly modelled). Despite this we do have sufficient information to determine the chirality of a given system given three points since we know the chiral center has coordinates: $o = (0, 0, 0)$. We can then construct planes consisting of $(o, v_1, v_2)$ and $(v_1, v_2, v_3)$, and compute the dihedral angle between them which will be either $\arcsin \frac{1}{\sqrt{3}}$ or $-\arcsin \frac{1}{\sqrt{3}}$ (0.6155 radians or -0.6155 radians).

$$\begin{aligned}
o &= (0, 0, 0) \\
v_1 &= (\sqrt{\frac{8}{9}}, 0, -\frac{1}{3}) \\
v_2 &= (-\sqrt{\frac{2}{9}}, \sqrt{\frac{2}{3}}, -\frac{1}{3}) \\
v_3 &= (0, 0, 1) \\
\text{Dihedral}(o, v_1, v_2, v_3) &= 0.6155 \,\text{radians}
\end{aligned}$$

To show that this angle has enough information to convey information about chirality, we invert the chirality of the system and compute the angle:

$$o = (0,0,0)$$
$$v_1 = (\sqrt{\frac{8}{9}}, 0, -\frac{1}{3})$$
$$v_2 = (-\sqrt{\frac{2}{9}}, \sqrt{\frac{2}{3}}, -\frac{1}{3})$$
$$v_3 = (-\sqrt{\frac{2}{9}}, -\sqrt{\frac{2}{3}}, -\frac{1}{3})$$
$$\text{Dihedral}(o, v_1, v_2, v_3) = -0.6155 \,\text{radians}$$

There are two pieces of information necessary to compute these angles: the two planes to consider (an ordering of four atoms) and the resultant ideal angle. It is not clear how to embed this information into a neural network such as RoseTTAFold (how do we embed the order of the indices used to compute the angles?) so we decided to take inspiration from physical modeling where gradients of energy functions with respect to coordinates are used to update structures. We decided to define a pseudo-energy term, differentiate it with respect to the previous predicted coordinates and then provide it to the subsequent block as a $\ell 1$ (vector) feature, the direction of the vectors on atoms in chiral centers breaks the symmetry with respect to reflections in the rest of the input features. The chiral energy function is defined as:

$$E_{chirality} = \frac{1}{N * M} \sum_{i=1}^{N} \sum_{j=1}^{M} (\theta^{i,j}_{\text{predicted}} - \theta^{i,j}_{\text{ideal}})^2$$

where $N$ is the number of chiral centers in the molecule, $M$ is the number of unique pairs of planes that can be constructed starting with a give chiral center that include only explicitly modeled atoms, and $\theta^{i,j}_{\text{predicted}}$ and $\theta^{i,j}_{\text{ideal}}$ are the predicted and ideal angles in chiral center $i$ and set of planes $j$.

The gradients of the chiral energy with respect to the coordinates are calculated using autodifferentiation in Pytorch [18].

$$chiral\_grads = \text{autograd}(E_{\text{chirality}}, xyz)$$

###### 4.5.3 Constructing *ℓ1* features

As shown with the chiral gradients, we can provide vector features for different inductive biases through *ℓ1* features. Inputs of the geometry of the previous N-C$\alpha$-C (or OP1-P-OP2) frames are necessary to make accurate updates of frame orientations. We provide these features as $\ell 1$ features representing the vectors between the N and C coordinates with respect to the C$\alpha$. These features are also provided for atom nodes with respect to their canonical frame (nodes without frames such as metal ions receive vectors full of 0s for this feature).

###### 4.5.4 Constructing Edge features

In the SE(3) Transformer, edge features are used to update nodes. Edges are directed so edge features $e_{i,j}$ are different than $e_{j,i}$. We begin by computing the sequence relative encoding and the atom distance features described in Algorithm 7. We then compute the radial basis function of the pairwise C$\alpha$ (or P or atom) distances from the previous predicted structure. The *pair* features, the sequence and atom separation features and the radial basis function features are concatenated and fed into a small network. This network consists of a Linear layers and a FeedForward Layer followed by a LayerNorm (same architecture as the network that process the $\ell 0$ *features*. The graph is constructed, with all nodes connected to all nodes (in both directions). The resulting graph, $\ell$ node features and edge features are processed by the SE(3) Transformer and predicts node-wise $\ell 0$ and $\ell 1$ features.

###### 4.5.5 Applying Structure Updates

The output $\ell 0$ features are the updated *state* features. The predicted $\ell 1$ features correspond to coordinate updates (in nanometers) and four values that are used to generate a quaternion as in AF2. The quaternion is converted into a rotation matrix and used to update the orientation of N-C$\alpha$-C and OP1-P-OP2 frames (the predicted quaternion values have no semantic meaning for atom nodes and the rotation matrices are explicitly set to the identity matrix so gradients are not computed). The first row of the *msa* representation and the *state* features are concatenated and fed into a small residual network and predict $\chi$ angles for protein residues and nucleic acid bases as in RF2 and RF2NA.

##### 4.6 Refinement Layers

The refinement layers are additional 3D update blocks that do not feed back into the main 1D and 2D updates. In RFAA there are 4 blocks with shared weights, which refine the final structure based on the 1D and 2D features. We make some slight changes from the 3D update in the main blocks. First, there is an additional $\ell 1$ feature. We compute an approximation of the Rosetta Leonard Jones potential (described in Section 5.7) and provide the network the gradient of the potential function with respect to each protein and nucleic acid frame atoms (and the singular atom for each atom node, it does not make sense to use the canonical frame in this case because each node only updates one atom). This is intended to provide the inductive bias to the network that atoms and protein residues should not be clashing. We also provide an extra $\ell 0$ feature which is the gradient of the LJ potential with respect to the $\chi$ angle predictions of the network at the previous block. Second, the graph is no longer fully connected. We expect that this stage of the network, the updates are reflective of local interactions and not global movements that would need a fully connected update. The changes are highlighted in Algorithm 10.

##### 4.7 Auxiliary Binding Head

The auxiliary binding head is a classifier network that classifies if two chains will bind or not. We trained the network to predict binding for protein complexes and protein-nucleic acid complexes as previously described.

##### 4.8 Auxiliary Error Prediction

Similar to the original RF, we predict the allatom LDDT of each residue and assess a cross entropy loss on the predictions (over 50 evenly spaced bins). We extend this to atoms by predicting the LDDT of the atom with respect to the other nodes present. We also predict two pairwise accuracy estimation metrics, predicted aligned error (pAE) and predicted distance error (pDE). Predicted aligned error (similar to AF2), aligns each frame (i) and computes errors over all C$\alpha$ (or P or atom) coordinates. For atom nodes, the canonical frames are aligned and the error is computed over all other nodes. We also developed pDE as an alternative which just predicted the unsigned distance error between any two nodes. We found empirically that pAE correlates better with accuracy so we do not report pDE statistics.

**Algorithm 10** RFAA Refinement Layer Update

**function** REFINEMENT_LAYER($msa_{cluster}, pair, state, xyz, idx, bond\_feats, dist\_matrix, chiral\_grads$, clash_l1_grads, $atom\_frames$, clash_l0_grads, $topk$)
  $seq$ = LayerNorm($msa_{cluster0}$) ▷ Query sequence latent embedding
  $pair, state$ = LayerNorm($pair$), LayerNorm($state$)
  $node\_feats$ = Linear(CONCAT([$seq, state$]))
  $node\_feats$ += FeedForward($node\_feats$)
  $node\_feats$ = LayerNorm($node\_feats$)
  $node\_feats$ = CONCAT([$node\_feats, clash\_l0\_grads$])
  rel_pos, bond_dist = Positional_Encoding(idx, bond_feats, dist_matrix)
  rbf_feat = rbf($||xyz_{C\alpha} - xyz_{C\alpha}||$) ▷ Or P or atom coordinate
  $edge\_feats$ = Linear(CONCAT([$pair$, rbf_feat, rel_pos, bond_dist]))
  $edge\_feats$ += FeedForward($edge\_feats$)
  $edge\_feats$ = LayerNorm($edge\_feats$)
  $frame_{ij}, frame_{kj} = frame_i - frame_j, frame_k - frame_j$ ▷ Construct frames from N-C$\alpha$-C, O-P-O or atom_frames
  l1_feats = CONCAT([$frame_{ij}, frame_{kj}, chiral\_grads$, clash_l1_grads]
  geometric_feats = $xyz_i - xyz_j$
  $state, coord\_update$ = SE3_Transformer(node_feats, edge_feats, l1_feats, geometric_feats, top_k= topk)
  $quaternion\_values, T = coord\_updates$
  $R$ = construct_rotation_matrix_from_quaternion($quaternion\_values$) ▷ Set R = Identity matrix for "atom" nodes
  $xyz = R \circ xyz + T$
  $sc\_torsions$ = SC_pred($msa_{cluster0}, state$)
  **return** state, xyz, sc_torsions
**end function**

### 5 Loss Functions

#### 5.1 Resolving Equivalent Atom Orderings

When predicting general biomolecular systems with systems at hybrid residue and atom resolutions, there are multiple symmetric atom labels that represent the same biomolecular system. First, certain protein sidechains contain 180° flips. Second, certain assemblies contain multiple identical protein chains which could be predicted in any order. Third, arbitrary assemblies might have multiple copies of the same small molecules that could be predicted in any order. Fourth, arbitrary small molecules can have symmetric chemical groups that could be reordered. We developed an algorithm which assigns the ordering that would minimize distance RMSD between a ground truth assembly and a predicted assembly. We handle symmetric swaps of sidechains similarly to AF2 and RF2, where the distances of each ambiguous atom to all non-ambiguous atoms is computed for the predicted and true structure and the closest atom naming assignment to the true structure for each permutation swap is assigned. The process for resolving the remaining symmetries is as follows: 1) Distances between C$\alpha$ coordinates in the predicted structure and C$\alpha$ distances in all valid permutation swaps of protein chains (see Section 3.2) are computed. Chains are assigned by finding the chain ordering that has the minimum difference between the distances in the predicted structure and distances in the true structure. 2) Given the anchor point of the protein ordering, every ligand is greedily assigned by the minimizing the difference between distances between atoms in the ligand to C$\alpha$ atoms in the predicted protein structure and true protein structure (based on the assignment in step 1). During the greedy assignment, all possible permutation swaps of atoms within each ligand are also considered.

##### 5.2 Masked Token Recovery

The masked token recovery loss is identical to the published RF2 [13, 19, 14]. In cases with small molecules or other biomolecules, we only mask protein tokens so there is no masked token recovery loss applied for other biomolecules.

##### 5.3 Torsion Angle Loss

The torsion angle loss is unchanged from RF2 and RF2NA. Atom nodes have no torsion angle loss applied, since none of those values have semantic meaning with respect to placing individual atoms.

##### 5.4 Distogram Loss

Following trRosetta [20] and RoseTTAFold [13], we apply a loss on predicted pairwise distance and orientations from the final pair representation. Several modifications were made to accomodate arbitrary biomolecules. First, we changed the binning of the distograms. For proteins in AF2 and RF2, the distogram loss is evenly spaced from roughly 2.5Å to 20Å. For small molecules, shorter distances (bond lengths, hydrogen bonds) are often more important so there are 30 evenly spaced distogram bins between 1Å and 4Å and then 30 more (coarser) bins between 4Å and 20Å. Three more angle losses representing inter-residue orientations are applied to a projection from the *pair* features. For atom nodes, the canonical frame is used to compute the three points for each angle. First, we compute a pseudo C$\beta$ atom for every frame (using default Rosetta params):

$$
\begin{aligned}
\vec{x} &= b - a \\
\vec{y} &= c - b \\
\vec{z} &= \vec{x} \times \vec{y} \\
d &= -0.57910144\vec{z} + 0.5689693\vec{x} - 0.5441217\vec{y} + b
\end{aligned}
$$

where a, b, c are three atom coordinates in a frame. While this coordinate ($d$) does not mean anything physically for nucleic acids or atom frames, we reasoned that this was a consistent geometric quantity so the network should be able to predict quantities derived from it given a local coordinate frame.

$D_{:,l,l'}$, $\Omega_{:,l,l'}$, $\Phi_{:,l,l'}$, $\Theta_{:,l,l'}$, together describe the orientation of residue $l$ relative to residue $l'$. The following loss consists of the cross entropy between the one-hot histogram of the known inter-residue distances and orientations and the corresponding distributions predicted by the model.

$$
\begin{aligned}
\mathcal{L}_{\text{2D}}(\text{logits}_d, \text{logits}_\omega, \text{logits}_\theta, \text{logits}_\phi, z_0) =& \text{CrossEntropy}(\text{logits}_{\text{dist}}, D) + \\
& \text{CrossEntropy}(\text{logits}_\omega, \Omega) + \\
& \text{CrossEntropy}(\text{logits}_\theta, \Theta) + \\
& \text{CrossEntropy}(\text{logits}_\phi, \Phi)
\end{aligned}
$$

where:

$$D \in \mathbf{R}^{[\mathbf{C_{dist}xLxL}]}; D_{b,l,l'} = \mathbb{1}[\text{bin}_{D,b}^{low} \leq \max(\|C_{\beta,l'} - C_{\beta,l'}\|_2, 18.5) < \text{bin}_{D,b}^{high}]$$
$$\Omega \in \mathbf{R}^{[\mathbf{C_{dist}xLxL}]}; \Omega_{b,l,l'} = \mathbb{1}[\text{bin}_{\Omega,b}^{low} \leq \text{Dihedral}(C_{\alpha,l}, C_{\beta,l}, C_{\alpha,l'}, C_{\beta,l'}) < \text{bin}_{\Omega,b}^{high}]$$
$$\Theta \in \mathbf{R}^{[\mathbf{C_{dist}xLxL}]}; \Omega_{b,l,l'} = \mathbb{1}[\text{bin}_{\Theta,b}^{low} \leq \text{Dihedral}(N_{\alpha,l}, C_{\alpha,l}, C_{\beta,l}, C_{\beta,l'}) < \text{bin}_{\Theta,b}^{high}]$$
$$\Phi \in \mathbf{R}^{[\mathbf{C_{phi}xLxL}]}; \Omega_{b,l,l'} = \mathbb{1}[\text{bin}_{\Phi,b}^{low} \leq \text{Planar}(C_{\alpha,l}, C_{\beta,l}, C_{\beta,l'}) < \text{bin}_{\Phi,b}^{high}]$$

and the bin edges for converting these angles and distances into a one-hot distribution are given by:

$$\text{bin}_{D,i} = \begin{Bmatrix} [0, d_{min}], & \text{if } i = 0 \\ [d_{min} + (d_{mid} - d_{min})\frac{i-1}{29}, d_{min} + (d_{mid} - d_{min})\frac{i}{29}], & \text{if } 1 < i < 30 \\ [d_{mid} + (d_{max} - d_{mid})\frac{i-30}{30}, d_{mid} + (d_{max} - d_{mid})\frac{i}{30}], & \text{if } 30 =< i < 60 \\ [d_{max}, \infty], & \text{if } i = 60 \end{Bmatrix}$$
$$\text{bin}_{\Omega,i} = \text{bin}_{\Theta,i} = [-\pi + \frac{2\pi i}{37}, -\pi + \frac{2\pi(i+1)}{37}]$$
$$\text{bin}_{\phi,i} = [\frac{\pi i}{19}, \frac{\pi(i+1)}{19}]$$

where $d_{min} = 1.2$, $d_{mid} = 4$, and $d_{max} = 20$

The formulas for computation of dihedral and planar angles are given by

$$\begin{aligned} \text{Dihedral}(a, b, c, d) = \text{atan2}( \\ &[c-b] \cdot (([b-a] \times [c-b]) \times ([c-b] \times [d-c])), \\ &\|c-b\|([b-a] \times [c-b]) \cdot ([c-b] \times [d-c])) \\ \text{Planar}(a, b, c) = \arccos(\frac{(a-b) \cdot (c-b)}{\|a-b\|\|c-b\|}). \end{aligned}$$

#### 5.5 All-Atom FAPE

Based on our experience training RF2 and RFNA, the FAPE loss introduced in the AF2 paper [5] was essential to producing accurate structures with the correct chirality. In RF2 and AF2, FAPE was applied only on the backbone frames for the intermediate generated structures and the full atom FAPE was applied on the final step and backpropagated through the whole network. Analogous to using N-C$\alpha$-C frames, we reasoned that we could use triplets of bonded atoms as frames, align the predicted and true atoms and compute the error over the other atoms.

The easiest way to do this to enumerate all triplets of bonded neighbors for each atom and apply a loss on the deviations on all atoms when aligning all the potential frames. For the "backbone" frame loss applied across all 40 (4 *msa_full* blocks, 32 main blocks and 4 refinement layers), we wanted to choose a single frame to apply a loss over for each atom node. We decided to implement an algorithm that deterministically chose a *canonical* frame for each atom node as described in Section 2.1.4. Each atom's *canonical* frame is used for the computation of FAPE. We believe that deterministic frame selection is important because it ensures FAPE is a stable optimization objective as if frames were stochastically sampled, the same predicted structures could have different FAPE values depending on the choice of

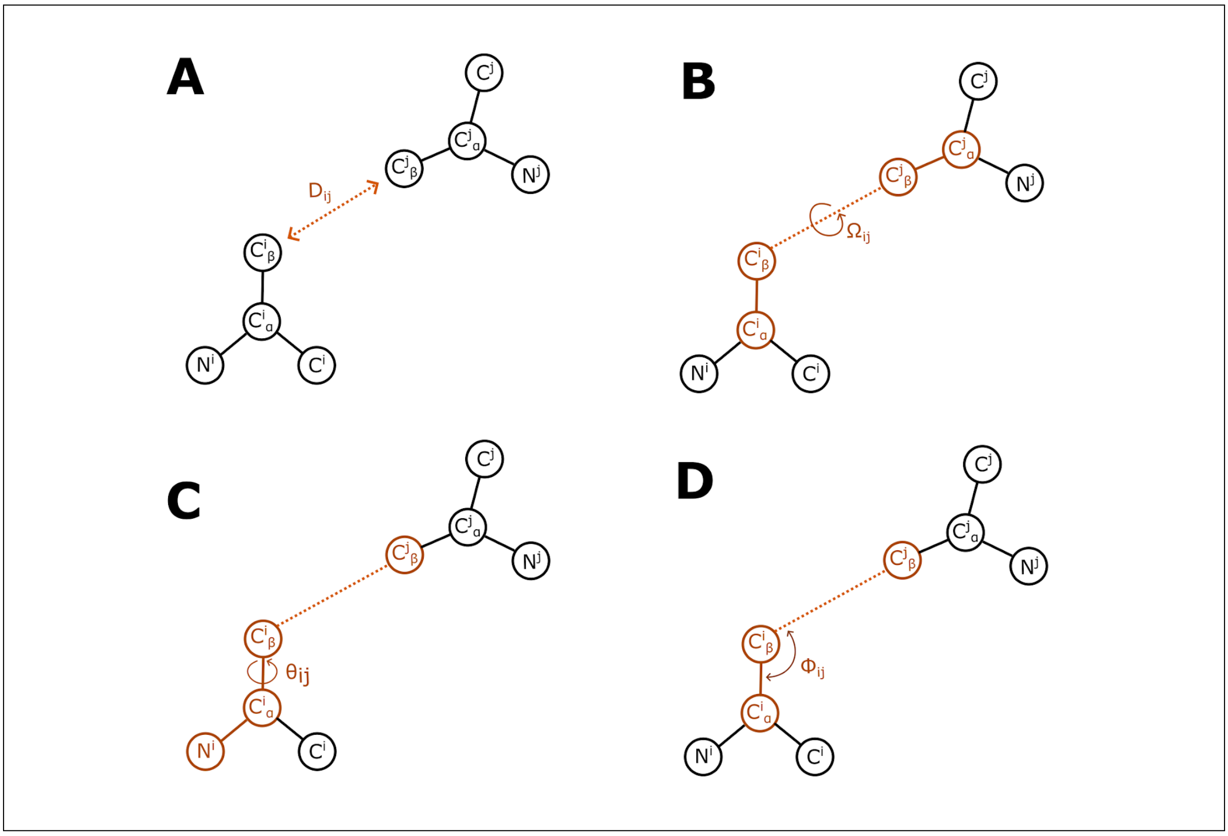

Supplementary Information Figure 1: Diagrams for how to compute the four inter-residue distance and dihedral degrees of freedom.

frames but did not test any alternatives. We also believe that deterministic frame selection is important for the frame orientation and predicted alignment error losses because the network has to predict quantities dependent on the chosen frame.

During early experiments, we found that the network was able to move groups of atoms closer to the desired pocket but did not make coherent ligand structures or interactions with the protein. We reasoned that there were two issues with our formulation of FAPE. First that the scale of distances that are important for small molecule structures are smaller than those that are important for proteins. Second, in a given training example, most nodes are protein residues so the errors of protein residues outweigh the errors of small molecules in the total loss calculation and the gradient to the weights of the network. We added an additional loss term that consisted of only the errors of the atom node coordinates with respect to the atom node frames with a normalization value of 4Å (compared to 10Å in AF2 and RF2). We also found that increasing the relative contribution of this term to the total loss helped balance the contributions of the protein errors and small molecule errors.

The second phenomenon (less contributions from atom nodes) also affected the contribution of the interchain FAPE values (errors of protein atoms with respect to atom frames and errors of atom coordinates with respect to protein frames). We chose to also add an auxiliary loss consisting of the interchain FAPE values and scaled its relative contribution to the total loss.

In the early training, we use an exponential schedule to weight backbone FAPE over the different layers of the network. During the fine-tuning stage of training, we equally weight FAPE across all layers. The

| Term | d_clamp | Z |
|---|---|---|
| $\mathcal{L}_{fapefullatom}$ | 10 | 10 |
| $\mathcal{L}_{fapebackbone}$ | 90% 30, 10% unclamped | 10 |
| $\mathcal{L}_{fapeintra-ligand}$ | 4 | 4 |
| $\mathcal{L}_{fapeinterproteinligand}$ | 90% 30, 10% unclamped | 10 |

Supplementary Methods Table 9: Parameters for different FAPE terms

following equation is used to weight backbone FAPE over all the intermediate layers of the network:

$$\mathcal{L}_{\text{fape backbone}} = \frac{1}{\sum_{i=0}^{I-1} \gamma^i} \sum_{i=1}^{I} \gamma^{I-i} \text{FAPE}(x^{(0)}, \hat{x}^{(0),i})$$

where $\hat{x}^{(0),i}$ is the $i^{\text{th}}$ structure block output and $\gamma$ is a scaling factor. The hyperparameters used for the different FAPE terms are shown in Table 9. We tuned these values in very early experiments and did not optimize them as we updated the network architecture so it is possible that better values exist for final performance.

To compute frames from an arbitrary set of three points we use a Gram-Schmidt process (identical to the rigid_from_3_points algorithm in AF2; referred to as Gram_Schmidt). Unlike AF2, we use this process on the predicted coordinates as well instead of using the chained set of rotations predicted by the network since small molecule nodes do not predict rotations. To compute FAPE, we first assemble a set of indices that comprise each frame, construct the 3 points for each frame, compute the associated rigid orientation of each frame in the global frame and apply the operation on each the predicted and true coordinates and measure the errors (shown in Algorithm 11).

**Algorithm 11** Compute All-Atom FAPE

**function** COMPUTE_GENERAL_FAPE(pred, true, atom_frames, Z, d_clamp)
  frames = construct_frames(atom_frames) ▷ Predefined frames for proteins/NA, atom_frames for atoms
  $pred_{frames}$ = gather(pred, frames) ▷ $N_{nodes}, M_{frames}$, 3 atoms,3 coordinates
  $true_{frames}$ = gather(true, frames) ▷ $N_{nodes}, M_{frames}$, 3 atoms,3 coordinates
  $T_{pred}^{-1}$ = Gram_Schmidt($pred_{frames}$)
  $T_{true}^{-1}$ = Gram_Schmidt($true_{frames}$)
  $x_{pred} = T_{pred}^{-1} \circ pred$
  $x_{true} = T_{true}^{-1} \circ true$
  $e_{ij} = \sqrt{(x_{pred} - x_{true})^2}$
  $\mathcal{L}_{fape} = \frac{1}{Z} mean_{ij}(clamp(e_{ij}, d\_clamp))$
  **return** $\mathcal{L}_{fape}$
**end function**

#### 5.6 Bond Geometry Losses

Generally, we found that the network learned accurate bond lengths and angles from the training set. We did notice that in some cases when the network was unsure on docking, it would produce unideal bond lengths and angles. We decided to place a small loss on bond lengths, angles, and distances within rigid groups (rings). We use a smoothed L1 Loss with beta=0.2. For bond lengths and rigid groups, we apply the loss on all pairwise distances (between two bonded atoms or atoms in a shared rigid group). For angles, we place the loss on distances predicted between every atom and atoms 2 bonds away in the bond graph. We reason that this in conjunction with the loss on bond lengths is sufficient to constrain the angles as well (similar ideas to those presented in [21]). We also apply this loss on bond lengths between atomized N atoms and the previous residue C and atomized C atoms and the following N. Losses on backbone geometry for proteins and nucleic acids are exactly the same as those in RF2. Bond lengths and angles are penalized compared their ideal values with a tolerance and the errors are normalized by the amount of violating predictions.

#### 5.7 Clash Loss

We found that initial trained versions of the network produced some clashes between protein residues and between protein residues and small molecule atoms. Following RF2, we intended to use a modified version of the Rosetta Leonard Jones potential (6-12 potential) [22] as an additional loss term to penalize the network from making clashing structures. Rosetta uses atom typing to determine the optimal parameters for the LJ potential and since our network operates on a larger set of atoms than have been tuned by the developers of Rosetta, we generated a set of parameters to use in our loss. These roughly correspond to the the mode of the parameters for each element type (most common parameter across all atom types for each element), parameters from UFF [23] where there are no available Rosetta parameters and default values for when there are no UFF or Rosetta parameters. We did not tune these values during model development.

| Element Type | LJ Radius | LJ Well Depth |
|---|---|---|
| Al | 1 | 0.1 |
| As | 1 | 0.1 |
| Au | 1 | 0.1 |
| B | 1 | 0.1 |
| Be | 1 | 0.1 |
| Br | 2.1971 | 0.1090 |
| C | 2.0067 | 0.0689 |
| Ca | 1 | 0.1 |
| Cl | 2.0496 | 0.1070 |
| Co | 1 | 0.1 |
| Cr | 1 | 0.1 |
| Cu | 1 | 0.1 |
| F | 1.6491 | 0.0750 |
| Fe | 1 | 0.1 |
| Hg | 1 | 0.1 |
| I | 2.36 | .111 |
| Ir | 1 | 0.1 |
| K | 1 | 0.1 |
| Li | 1 | 0.1 |
| Mg | 1 | 0.1 |
| Mn | 1 | 0.1 |
| Mo | 1 | 0.1 |
| N | 1.7854 | 0.1497 |
| Ni | 1 | 0.1 |
| O | 1.5492 | 0.1576 |
| Os | 1 | 0.1 |
| P | 2.1290 | 0.5838 |
| Pb | 1 | 0.1 |
| Pd | 1 | 0.1 |
| Pr | 1 | 0.1 |
| Pt | 1 | 0.1 |
| Re | 1 | 0.1 |
| Rh | 1 | 0.1 |
| Ru | 1 | 0.1 |
| S | 1.9893 | 0.3634 |
| Sb | 1 | 0.1 |
| Se | 1 | 0.1 |
| Si | 1 | 0.1 |
| Sn | 1 | 0.1 |
| Tb | 1 | 0.1 |
| Te | 1 | 0.1 |
| U | 1 | 0.1 |
| W | 1 | 0.1 |
| V | 1 | 0.1 |
| Y | 1 | 0.1 |
| Zn | 1 | 0.1 |

Supplementary Methods Table 10: LJ Loss Parameters for Atom Tokens

Supplementary Methods Table 11: Training Hyperparameters

| | Phase 1 | Phase 2 | Phase 3 |
|---|---|---|---|
| Crop Size | 256 | 256 | 375 |
| Batch Size | 64 | 64 | 128 |
| Learning Rate | 0.001 | 0.001 | 0.0002 |
| Schedule | No warmup. Decay Learning Rate by 0.95 every 5000 steps | No warmup. Decay Learning Rate by 0.95 every 5000 steps | No warmup. Decay Learning Rate by 0.95 every 5000 steps |
| Loss | $10 * \mathcal{L}_{fapefullatom} + 5 * \mathcal{L}_{fapebackbone} + 10 * \mathcal{L}_{fapeintra-ligand} + 2.5 * \mathcal{L}_{fapeinterproteinligand} + 1.0 * \mathcal{L}_{c6d} + 3.0 * \mathcal{L}_{msamask} + 10 * \mathcal{L}_{torsion} + 0.1 * \mathcal{L}_{plddt} + 0.5 * \mathcal{L}_{pae} + 0.5 * \mathcal{L}_{pde}$ | $10 * \mathcal{L}_{fapefullatom} + 5 * \mathcal{L}_{fapebackbone} + 10 * \mathcal{L}_{fapeintra-ligand} + 2.5 * \mathcal{L}_{fapeinterproteinligand} + 1.0 * \mathcal{L}_{c6d} + 3.0 * \mathcal{L}_{msamask} + +10 * \mathcal{L}_{torsion} + 0.1 * \mathcal{L}_{plddt} + 0.5 * \mathcal{L}_{pae} + 0.5 * \mathcal{L}_{pde}$ | $10 * \mathcal{L}_{fapefullatom} + 5 * \mathcal{L}_{fapebackbone} + 10 * \mathcal{L}_{fapeintra-ligand} + 2.5 * \mathcal{L}_{fapeinterproteinligand} + 1.0 * \mathcal{L}_{c6d} + 3.0 * \mathcal{L}_{msamask} + 10 * \mathcal{L}_{torsion} + 0.1 * \mathcal{L}_{plddt} + 0.5 * \mathcal{L}_{pae} + 0.5 * \mathcal{L}_{pde} + 0.5 * \mathcal{L}_{bind} + 0.02 * \mathcal{L}_{protein,NAgeom} + 0.02 * \mathcal{L}_{atomgeom} + 0.02 * \mathcal{L}_{clash}$ |
| top_k in Refinement Layers | 128 | 128 | 64 |
| Exponential Decay of FAPE over Str Layers | 0.99 | 0.99 | 1.0 |
| Optimizer Steps | ~35e3 | ~40e3 | ~15e3 |

### 6 Training Details for RFAA Structure Prediction

RFAA was trained in three phases which were intended to interrogate the question of whether a single model could represent all biomolecules. We iteratively added in different datatypes and tracked accuracy using the held out validation clusters. We observed that adding new datasets did not decrease accuracy on datasets that were already present in training. We anticipate that the network could be trained on all the datasets simultaneously and achieve the same accuracy but have not tested this rigorously. The first two phases were the main training phases and the final phase was a fine-tuning phase with different hyperparameters and loss functions.

#### 6.1 Training Hyperparameters

The first two stages of training are analagous to the initial training of RF2 and AF2 and the third stage is similar to the "fine-tuning" stage where violation losses are turned on, crop size increased and hyperparameters changed to refine the accuracy of the network. Exact hyperparameters used are shown in Table 11.

#### 6.2 Dataset Sampling

At the outset of this work, it was unclear to us whether it would be possible to train a single network to model all biomolecules in the PDB. We decided to add in new datasets as training progressed to assess whether the network could simultaneously learn features from all datasets. We restarted training

Supplementary Methods Table 12: Dataset Sampling Proportions

| **Dataset** | **Phase 1** | **Phase 2** | **Phase 3** |
|---|---|---|---|
| Protein Monomer | 0.09 | 0.07 | 0.12 |
| AF2 Distillation | 0.0 | 0.14 | 0.36 |
| Protein Complex | 0.17 | 0.11 | 0.055 |
| Negative PPI | 0.0 | 0.0 | 0.055 |
| Protein-Nucleic Acid Complex | 0.17 | 0.11 | 0.055 |
| Negative Protein-Nucleic Acid Complex | 0.0 | 0.0 | 0.055 |
| RNA | 0.09 | 0.06 | 0.05 |
| Protein-Small Molecule Complex | 0.37 | 0.20 | 0.11 |
| Protein-Metal Complex | 0.0 | 0.10 | 0.03 |
| Protein-Multi-residue Molecule Complex | 0.0 | 0.05 | 0.0275 |
| Covalently Modified Protein | 0.0 | 0.05 | 0.0275 |
| Protein-Small Molecule Assembly | 0.0 | 0.0 | 0.055 |
| CSD Crystal Structures | 0.03 | 0.03 | 0.03 |
| Atomized Protein Augmentation | 0.08 | 0.08 | 0.04 |

in three phases with different dataset proportions. We found generally adding new datasets did not decrease accuracy on the datasets that were already trained, although it is unclear whether we would have seen further improvements if we continued training without adding the new datasets.

Each dataset was sampled with a given probability and then each sequence cluster within that dataset is sampled with inverse probability to the number of examples in the cluster. Within the protein monomer clusters, if examples are homomers we stochastically sample featurizing them as monomers or homomers ($p_{\text{homomer features}} = 0.5$).

We do not believe that the exact values of the dataset samples are essential to recover the accuracy that we observe in the paper. The values were tuned with three guiding principles, 1) to balance out sequence clusters sampled early in training, 2) to show equal amount of positive and negative examples and 3) to bias towards the larger datasets towards the end of training to avoid overfitting on smaller datasets. The dataset sampling proportions are shown in Table 12.

### 7 Structure Prediction Inference Regimen

At inference time, we run the model without MSA corruption or template subsampling, instead preserving all MSA tokens and picking the top 4 searched templates for each entry. We run the model for 10 recycles unless otherwise specified. For every protein sequence, we build MSAs and templates using the standard MSA and template generation pipeline described in Section 1.1.

We define the *ligand RMSD* as follows: for each ligand target, we compute in the crystal structure every backbone atom that is 10Å of the any atom in the bound ligand. We then kabsch align the predicted and crystal protein structures on the aforementioned backbone atoms, and use the same transformation matrix to superimpose predicted and crystal pose ligands. The ligand RMSD is the resultant RMSD between predicted and true ligand positions. We use this metric throughout our evaluation of protein-ligand complex predictions. We note in figure legends when ligand RMSD is computed by an external tool (eg. for CAMEO evaluations and Posebusters).

The model makes predictions of its own error during training time in a 2D matrix called the predicted alignment error as desrcibed in 4.8. The $i, j$th entry is trained to be an estimate of the $\ell^2$ distance error between the the $j$th atom in the $i$th coordinate frame. We define the inter-chain protein-ligand predicted alignment error (PAE Interaction) as the mean of the predicted alignment error tensor between all protein residue frames and small molecule coordinates, and all small molecule frames and residue coordinates.

We do not perform any cropping at inference time, predicting full complexes (including residues/atoms that were not resolved in the true structure).

#### 7.1 CASP14 Protein Monomer Targets

We benchmarked the performance of RFAA against RF2 and AF2 using a subset of the CASP14 targets with experimentally resolved structures that were not removed during competition. TBM-easy targets were discarded and the final set of 42 proteins was composed of monomeric targets from TBM-hard, FM/TBM, and FM categories. Each of the three methods were run using default parameters with the same input multiple sequence alignments and no templates.

- AF2 (V1 weights), model_1, and 20 recycles
- RF2 (Apr23 weights), model 1, and 20 recycles
- RFAA (this work), model 0, and 20 recycles

GDT-TS was calculated based on experimentally solved structures provided by CASP using TM-score (accessed Jun, 2023) [24]. MSAs were generated as described in [25]. Briefly, query sequences were searched iteratively with HHblits [2] against uniclust30 (UniRef30_2020_01) database [26] with a gradient of E-value cutoffs and 95% sequence identity filtering. For targets with shallow MSAs, we converted the uniclust30 generated MSA into a seed HMM to search against JGI [27] with hmmsearch [28]. Homologous sequences from this search were aligned and combined with previous uniclust30 sequences.

#### 7.2 CAMEO Targets

The CAMEO BETA challenge (`https://beta.cameo3d.org/`) is an online, weekly, continuous evaluation of the most recent depositions into the PDB. We registered the RFAA model as a server for homomeric and ligand targets (excluding RNA, DNA and heteromeric targets for simplicity). The RFAA server searches for MSAs and templates for each protein sequence as described above. The server makes 5 different predictions with different random seeds for each target and then submits the prediction with the lowest interchain PAE. If there are multiple ligands in a given target, the interchain PAE is calculated over all ligands in the target.

We ran the RFAA CAMEO server (Server 2) from 04/08/2023 to 09/02/2023. Ligand pose baselines were introduced from 08/12/2023 onward (`https://beta.cameo3d.org/cameong_servers/`). The CAMEO organizers returns ligand RMSD poses scores that we then plot for our evaluations. We do not compute our own RMSD metrics for the CAMEO challenge.

The CAMEO server does not provide stoichiometry information for either protein or ligand in a given protein ligand complex. In order to determine protein stoichiometry, we check the symmetry group of each template of the input protein. We make a list of the top 20 most similar templates to the input protein by sequence similarity, and then filter that list based on a coverage threshold of 0.75 and a match score of 0.95. For the passing templates, if at least half of them form a dimeric or a trimeric complex, we then attempt to predict a dimeric or trimeric complex with the input protein, respectively. If the predicted alignment error between the symmetric subunits is less than 10 and the computed clash score between the subunits is less than 1, we model the protein as either a dimer or a trimer. Otherwise, we default to modeling the protein as monomeric.

We only make predictions with a single copy of each ligand in each target and do not duplicate ligands in the case of dimer or trimer-forming protein chains. We found it difficult to distinguish between cases where there are two binding pockets for two different copies of the ligand in a dimer compared to a single copy of the ligand sitting at the dimeric interface. We also do not model symmetry groups above C3 because they often do not fit in GPU memory on the allocated GPUs.

#### 7.3 Recent PDB Evaluations

##### 7.3.1 Protein Nucleic Acid Complexes

We evaluate RFAA on Protein Nucleic Acid complexes using a dataset of recently deposited PDBs curated in [7]. RFAA and RFNA are evaluated using the same MSAs and templates using the default parameters described. For this benchmark, all small molecule and noncanonical amino acid/base context is ignored.

We repredicted a small subset of predictions where the protein-NA interface was predicted accurately and there was small molecule also bound near the NA interface (an example shown in Fig 2D). We did not rigorously test the model's ability to model ternary complexes and expect that adding them into training (perhaps by fine-tuning) would significantly improve accuracy.

##### 7.3.2 Small Molecule Protein Complexes

We curate a list of protein-ligand complexes deposited in the PDB from 2021 and onwards. For each ligand in every entry in the PDB past this deposition date, we make a dataset of items defined by the procedure outlined in 1.3. We add the additional constraint that every ligand in this evaluation set must be designated as a "subject of investigation", a label which indicates that the authors who deposited the crystal structure believe the ligand to have some significance rather than being, for example, a solvent molecule. We note that this is not a perfect filter: certain PDB entries have ligands marked as subject of evaluation that are not referenced in the main text of the associated papers.

In order to remove redundancy from this evaluation set, we clustered the primary protein partners of each item down to 30% sequence identity at 80% coverage using MMSeqs2 [10]. For each sequence cluster, we compute the set of unique query ligands in that cluster and select, uniformly at random, a single protein-ligand complex for each unique query ligand. This clustering process ensures that we do not bias our evaluations by having many repeated copies of similar protein ligand binding pockets. For each item in the dataset, which consists of a query ligand and its immediate protein and non-polymer binding partners, we compute the total length (number of residues + number of atoms) of the assembly and then filter out all items of length $> 1000$.

The final held-out evaluation dataset consists of 5421 items. Of these, 897 consist of ligands that are covalently bonded to the protein and 622 of them are metal ions.

###### 7.3.3 PoseBusters

The PoseBusters dataset carefully curated subset of the protein-ligand complexes in the PDB released on or after 2021 designed for evaluating the performance of docking methods [29]. Each item in the PoseBusters dataset consists of a PDB entry and a ligand identifier specifying the ligand to be docked. There are 428 protein-ligand complexes with non-redundant protein chains and non-redundant, drug-like ligands.

We pre-process the data in largely the same way as we described in Section 1.3, with a few distinct changes. First, the posebusters dataset pre-specifies the *query ligand* of interest. Second, we narrow the definition of contacting protein chain to be any protein that has at least one atom within 10Å of the query ligand. Third, we define any contacting cofactors to the query ligand as any cofactor that has at least one atom within 5Å of the query ligand. Finally, if there are multiple identical cofactors within the context of the query ligand, we keep only one such copy. This slightly narrowed definition of query ligand context allows the network to focus more on docking the query ligand, which is the purpose of the PoseBusters evaluation.

For each item in the PoseBusters evaluation set, we predict the entire complex - query ligand, contacting protein partners, and contacting cofactors - simultaneously. We superimpose the primary protein partner onto the crystal structure and extract the crystal pose of the query ligand. We then run the posebusters suite on the predicted pose of the molecule to determine chemically validity of the predicted ligand pose. We also use the posebusters suite to compute ligand RMSD between crystal and predicted ligand pose after protein superimposition, and to compute validity metrics between predicted ligand pose and *predicted* protein structure (and predicted cofactors).

We note that our model is not a docking method and does not take as input the crystal structure of the protein nor the binding site of the query ligand, making our task (prediction from sequence information alone) strictly harder than blind docking or local pose estimation. This implies that it does not make sense to compute validity metrics, such as minumum distance clashes, between predicted ligand pose and *crystal* protein structure, which is why we use the model's predicted protein structure instead.

Finally, we also note that although our model is only trained on data from the PDB up to April 2020 and the PoseBusters dataset is only defined on data from 2021 onwards, our training data is defined on the deposition date of the PDB entries, while the PoseBusters dataset is defined on the entry release date. In almost all cases the dates of deposition and release are similar enough that the items in the PoseBusters evaluation set are not in our training set. However, a single entry (6VTA, AKN) was deposited before April 2020 but released in 2021. We exclude this item from the evaluation set for all methods.

###### 7.3.4 Covalent Modifications

Our evaluation set of covalent modification consists of the 897 entries described in Section 7.3.2. These items underwent the same filtering described in Section 1.3.3. We excluded all metal ions that have *covale* headers in the PDB because these are trivial modeling cases. During evaluations, we split the predictions into three categories based on their putative biological functions shown in Table 13. We used the PDB 3-letter codes to identify cases from our evaluations in each set. For multi-residue ligands with covalent bonds, we included them in a section if any of the residues were present in any of the categories. There were no multi-residue ligands where multiple residues were in multiple categories.

Supplementary Methods Table 13: Types of Covalent Modifications

| **Type of Modification** | **PDB 3-letter Codes** |
|---|---|
| Glycosylation | BGC,MAN,NAG,FUC,BMA,GLC,RIB,<br>NGC,GAL,BDP,A2G,JIW |
| Enzyme Cofactors | PLP,HEC,F3S,SAH,ADP,ATP,UDP,<br>FAD,AMP,FMN,ANP,HEM |

| | |
|---|---|
| Other | UVT, 24N, V4B, 2GI, 9JT, 2I5, 21I, V48, TJB, V3Q, 1XZ, WLP, TJ8, 2I8, V4N, TKK, SGM, HC4, S7Q, ZFG, S7N, RET, 9IX, N36, V0G, S08, DWZ, UZM, 2RG, WV0, S0Q, UQN, MW0, UZS, RZZ, PNS, IM2, RY2, CYC, ID1, 2LJ, ISS, UZV, RET, 4D6, 30W, FDX, N2Q, NXL, RXW, RET, 1S6, CMC, QS8, ZZ7, UPQ, MER, MWC, UZY, L9U, HH8, L8X, IIY, IJI, IK3, HNU, 82Q, LW1, L9C, 8NB, LBI, V7G, VR4, V46, YHI, XTM, YHJ, 2RG, UHS, UJ1, DWZ, XV4, VEV, 9JT, Q8H, PLM, USA, VEG, VEJ, XTP, RW8, XTJ, 0WN, CHL, VEM, UHY, DWZ, ZL7, UPD, VEP, 9JT, 2RG, ZJ1, Q8E, USD, UHV, VEY, YCV, UPJ, XC4, W48, UED, Y48, UJ4, EKZ, UUK, O1K, US8, 9ZG, 2IE, USZ, 2IJ, QVR, UQW, YWJ, 5YZ, URK, AMI, USH, SUU, R1L, UED, O1R, UUW, 1LE, 5ZB, ALD, A0U, O10, RN2, B1S, 7VB, QNC, AG7, 4W8, IFO, 91Z, NNA, SIN, U5G, 8GW, 7YW, I8H, SV6, PXQ, SFW, 90X, I1W, 8T6, I68, I71, 7VQ, 7W5, 4WI, 90U, R8H, 7YB, E9H, RBL, IRR, 7YZ, Q56, 8ZI, H60, H63, MYC, I70, EOF, 4IT, 2XI, HC1, IS5, 99W, TG3, H6L, 800, 7VW, 90I, HF0, I80, 7XK, C7A, HER, IRZ, 7VI, I54, NEN, 90H, G7L, 9SW, CB1, 8UI, K36, 2RI, ALD, 4YG, GKF, S9E, 7YQ, 8I0, 8I7, RN2, 8H9, 81L, T8M, SGH, IYB, 8G9, H6F, ISG, S8H, GQU, 91I, S8E, NOL, R7Q, 8H3, 7XT, I1Z, I69, S4L, 9JT, 7V2, EW9, G7O, IY8, H6R, NB2, 7YI, 7Y2, JMF, WHL, WHL, V08, WHL, 9AI, E3J, WHL, JVF, 9JT, MYR, WHL, WHL, S5K, AAC, WHL, QOK, WHL, WHL, 03S, FAR, DEP, I1J, 3CN, QNN, QN8, 6SI, 6BI, YB4, PLM, F5L, FWI, FV5, VKH, CLR, EIB, F8C, N63, T3K, PEB, |

| | |
|---|---|
| Other (continued) | Z41, 2DA, FEY, HYR, FHS, FZI, ACM, FVE, 1RG, BOV, U88, 1S7, R28, WFD, M1V, PNS, 90F, VOY, HHL, VO7, WF7, 5JR, NXL, 70I, BVX, AR6, RQT, RQZ, DTT, R1W, NFF, R2E, WZG, ZN, UOT, P5N, UPK, MHH, UON, 8I4, VX5, UFH, TJ8, RET, P8K, TCK, PLM, VLD, RET, X2S, MU4, YS7, X2G, DUT, YY3, DYF, X2J, PLR, Y37, 8BS, C9D, 4KZ, THR, I8K, S1N, MUB, RET, U8W, UT8, RET, 7J8, FP6, RET, DPM, U8Z, USW, RET, UST, NZ6, H40, PXQ, RFT, J2C, 8Z6, FAR, J3X, 88T, JRA, H2T, H0O, I6T, VU1, QTU, VU4, J50, V4T, ACE, ZGV, VLE, NZX, I7H, FHR, W6X, QPE, MYR, BZ2, G7F, PLM, QTU, QPB, Q5W, G7L, CLR, MTX, MEZ, CA, HNO, YJG, G5U, PLM, G8C, BFB, OCA, DON, UVB, H2S, V3N, V48, NEN, UWK, V2E, UVH, V2W, V3W, V32, V1H, V3Z, UXN, V3K, V2N, UVN, CPS, V3T, UWQ, V0T, V1E, V42, V2T, 4Y8, V3B, V1Q, VEE, RVW, UWH, V4K, V2K, V4H, UUB, V1K, V2Q, V0W, PAM, 9S5, 9TC, 7ID, CYC, 9Y6, NAP, S1K, S1B, M1V, VL5, CR8, PEB, ZXQ, DBV, VMM, PMS, WHL, FFQ, M1V, 9SS, GIT, S0Z, CYC, S1E, CYC, SQW, Q5O, 7TC, NW3, CEF, TSL, J00, ZXQ, Q5X, PLR, Q8F, SRQ, SRH, LBX, 6LN, OC0, O2O, LD0, 85Q, 4D6, R0B, Q33, Q2X, FVC, IM2, O0U, HJ9, TBE, TVE, AIX, 5YW, Q4C, ODN, D6M, PN7, DON, DTT, 06D, AG7, J6O, XSA, 05Y, UKS, KK9, BO2, 0G6, PEX, BJ3, CXS, BQG, GBS, RET, 0GJ, MYR, MYR, ME7, TA1, Y2D, UFY, SJK, G7M, CYC, 5GP, VHV, 5TI, XQV, PEB, UGH, XM2, GHX, LBV, VHP, MG, VHJ, JQI, GDP, XAG, ZOY, 81T, T5Q, TZK, PBW, VHM, T5Z, W6Z, VF2, 323, BG3, HP6, T5W, H9M, LNK, H9P, |

| | |
|---|---|
| Other (continued) | PLM, D12, W6X, SPH, RET, RET, H8S, H8V, T5N, V0Q, PNS, QE8, T6H, T4Z, D10, T6W, UHZ, T4W, OEH, T4Q, 868 |

###### 7.3.5 Metal Ions

622 of the entries in Section 7.3.2 are metal ions. As in our training data, we filter out examples that are not in Table 2.

##### 7.4 Computing Sequence Similarity to Training Set

For each item in our evaluation set, we BLAST [30] (with the `-qcov_hsp_perc 50` option and otherwise default parameters) the primary protein partner against all protein chains seen in our training set and compute the maximum sequence identity percentage to any protein chain seen during training. We consider a validation protein to have a training set match if it there exist a protein seen during training that has $> 30\%$ sequence identity at 50% coverage.

##### 7.5 Computing Ligand Similarity to Training Set

To compute similarity of ligands, we download idealized coordinates of all ligands in the PDB in sdf format (see `https://www.rcsb.org/downloads/ligands`). We use OpenBabel to compute the Morgan fingerprint for each ligand. We compute, for every ligand in the evaluation set, the bitwise similarity between the Morgan fingerprints for that ligand and every ligands deposited in the PDB, which we henceforth refer to as the Tanimoto similarity. We consider a ligand "similar" to a ligand seen in training if it has a Tanimoto similarity $> 0.5$ to any ligand seen during training.

##### 7.6 Evaluating effects of protein similarity and ligand similarity to training set on accuracy

In order to investigate whether or not RFAA can generalize to examples beyond those seen during training, we consider two notions of "similarity" to the training set: protein sequence similarity and the aforementioned ligand similarity.

To evaluate the dependence of protein similarity on accuracy, we clustered the sequences curated in Section 7.3.2 using MMSeqs2 (since we took every unique protein-ligand pair, certain protein sequence clusters were overrepresented). To determine whether it was possible for the network to produce accurate predictions with that sequence cluster, we extracted the lowest ligand RMSD prediction in each cluster. We then BLAST all the cluster representatives against the training set to determine the maximum similarity to any training example.

We followed a similar procedure to measure the impact of ligand similarity on accuracy. We computed an all by all matrix of Tanimoto similarity in our test set. We remove multi-residue ligands from this evaluation since the PDB does not provide ideal coordinates with the entire bonded ligand (only each residue) and the construction of the files can affect the Morgan fingerprint calculation. We then performed agglomerative clustering [31] with a threshold of 0.5 and selected the lowest RMSD representative from each cluster. All of the PDB 3-letter codes from the cluster representatives are mapped back against all the 3-letter codes used during training to measure the maximum Tanimoto similarity between each test set item and each training set item.

#### 7.7 Evaluating Correlation between RFAA Accuracy and Native Complex energies

We expected that a network that has learned general principles about protein-small molecule binding would make more accurate predictions for tighter binding interfaces. We performed Rosetta energy calculations on the native complexes from our test set (described in 7.3.2). Since small bond length and angle deviations in native structures can cause large energy differences, we choose to run a short minimization protocol to equilibrate the structures into the Rosetta forcefield. We use a recent protocol described in [32] where Rosetta's small molecule docking method, GALigandDock, is used in "eval" mode to evaluate a pose. In this mode, the ligand and any residues within a heavy atom contact distance of less than 8Å can move. The energy of the conformation is minimized using GALigandDock's generalized ligand potential and a harmonic constraint on the starting coordinates to prevent large movement. The pose is then scored using the generalized ligand potential in Rosetta. We remove cases where Rosetta does not have appropriate atom-typing or where the ligand moves more than 0.5Å from the starting position since those do not accurately score the native binding complex.

### 8 Training Details for RFdiffusionAA

RFDiffusionAA was trained on protein monomer structures in the PDB used for RFAA training 50% of the time and protein monomer/small-molecule complexes 50% of the time. Two small changes were made to the training set for small molecules: first, all cases with small molecules with unresolved atoms were removed and second, chains longer than 384 residues were removed (following RFDiffusion). As in RFDiffusion, training examples consist of the unconditional task 20% of the time and a motif-conditional task 80% of the time. The motif-conditional task comprises three distinct tasks:

- Middle motif [40%]: A contiguous set of residues
- Terminal motif [40%]: Two contiguous sets of residues at the N and C terminus of the monomer.
- Sparse contacts [20%]:
  - *For proteins*: A random set of 3 residues all $> 10$ residues apart in sequence space but with pairwise $C_\beta$–$C_\beta$ distances $< 6$Å is selected to form a model "active site". These 3 residues are included in the motif, and for each, there is a 50% chance of including one flanking residue. If no such triad is found in the monomer the task would fall back to Middle motif or Terminal motif with equal probability.
  - *For protein:small-molecule complexes*: A number of residues $n$ is selected from $\mathcal{U}(1,7)$. A candidate set of residues is then constructed from the set of the $n+2$ closest residues to the small molecule union the set of any residues within 2Å of the $n^{th}$ closest residue. $n$ residues are then selected at uniform from this candidate set.

During training, the structure module blocks are not allowed to update the positions of the motif residues.

**Losses:** RFDiffusionAA was trained with a loss comprising two terms that closely follows the loss used in RFdiffusion:

$$\mathcal{L}_{\text{Diffusion}} = \mathcal{L}_{\text{Frame}} + w_{\text{2D}}\mathcal{L}_{\text{2D}},$$

$\mathcal{L}_{\text{2D}}$ is the same as that used in Section 5.4, with the modification that motif residues are not included in its calculation.

Where $\mathcal{L}_{\text{Frame}}$ is exponentially weighted over the intermediate structure module outputs, increasing towards the end of the network:

$$\mathcal{L}_{\text{Frame}} = \frac{1}{\sum_{i=0}^{I-1} \gamma^i} \sum_{i=1}^{I} \gamma^{I-i} d_{\text{Frame}}(x^{(0)}, \hat{x}^{(0),i})^2$$

where $\hat{x}^{(0),i}$ is the $i^{\text{th}}$ structure block output and $d_{\text{Frame}}$ is a weighted mean squared error which includes clamping on displacement.

$$d_{\text{Frame}}(x^{(0)}, \hat{x}^{(0)}) = \sqrt{\frac{1}{L}\sum_{l=1}^{L}\Big(w_{\text{trans}}\min(\|z_l^{(0)} - \hat{z}_l^{(0)}\|_2, d_{\text{clamp}})^2 + w_{\text{rot}}\|I_3 - \hat{r}_l^{(0)\top} r_l^{(0)}\|_F^2\Big)},$$

where $w_{\text{trans}}$ and $w_{\text{rot}}$ are weights on the rotation and translation distances, and $d_{\text{clamp}}$ is a maximum distance above which translation distances are clamped. Note that the translation distance is only clamped $p_{\text{clamp}}$ of the time.

The forward noising process uses the same functional form as RFdiffusion with the same parameter set. For further details on the noise schedule see [33].

**Training time:** RFDiffusionAA trained to convergence when initialized from RFAA weights in 10 epochs. This took 8 days on 8 NVIDIA A6000 GPUs.

| **Parameter name** | **Value** |
|---|---|
| Crop size | 256 |
| Pseudo-batch size | 48 |
| $w_{\text{trans}}$ | 0.5 |
| $w_{\text{rot}}$ | 1.0 |
| $w_{\text{2D}}$ | 1.0 |
| $d_{\text{clamp}}$ | 10 |
| $p_{\text{clamp}}$ | 0.9 |
| Structure block iteration decay rate $\gamma$ | 0.99 |
| Learning rate | 0.0005, No warm-up. Decay learning rate by 0.95 after every 10000 optimization steps. |
| Examples per epoch | 25600 |
| Number of diffusion timesteps (T) | 200 |
| Variance schedule for translations | $\beta^{(t)} = \beta^z_{min} + (\frac{t}{T})(\beta^z_{max} - \beta^z_{min})$ with $\beta^z_{min} = 0.01$ and $\beta^z_{max} = 0.07$. |
| Variance schedule for rotations | $\sigma_t = \sigma_{min} + \frac{t}{T}\beta^r_{min} + \frac{1}{2}(\frac{t}{T})^2(\beta^r_{max} - \beta^r_{min})$, with $\sigma_{min} = 0.02$, $\beta^r_{min} = 1.06$, and $\beta^r_{max} = 1.77$ |
| Probability of motif being contiguous or discontiguous | 0.5 |
| Probability of providing self-conditioning information | 0.5 |
| Coordinate scaling | 0.25 |

Supplementary Methods Table 14: RFDiffusionAA training hyperparameters.

### 9 *In Silico* Design Methods with RFdiffusionAA

In this section, we will describe the *in silico* methods we used to benchmark the ability of RF*diffusion* AA to generate small molecule binders and the methods used to design the proteins that were characterized experimentally.

#### 9.1 Ligand Contact Potential

In RFdiffusion, it was shown that using a contact potential which biased trajectories towards forming interactions between protein chains improved success rates for generation of symmetric oligomers. We implemented a similar potential in RFdiffusionAA. These heuristic potentials update the $C_\alpha$ coordinates after each denoising step: $x^{(t-1)} = x^{(t-1)} + g(t)\nabla_{x^{(t)}} P(x^{(t)})$. Further motivating details can be found in [33]. The ligand contact potential we use is described below.

Denoting the coordinates of a ligand with $K$ atoms by $s = \{s_k\}_{k=1}^{K}$ and the C coordinates of a protein by $z = [z_1, \ldots, z_L]$:

$$P_{\text{contact}}(z, s) = \sum_{1 \leq l \leq L} \text{Switch}(\min_{1 \leq k \leq K} \|z_l - s_k\|_2^2)$$

This potential is then multiplied by a time-dependent guide scale $g(t)$. In the *in silico* benchmark and bilin binder design cases a linear guide scale is used $g(t) = \frac{t}{T}$.

#### 9.2 Small Molecule Binder Design Benchmark

We chose four small molecule targets to benchmark based on the following criteria: 1) the molecules are diverse in size and topology with respect to each other 2) two molecules are found in the training set and two molecules are novel to the training set ($<$ 0.5 Tanimoto similarity to closest ligand in training; deposited after our training date cutoff). The chosen molecules are (by PDB 3-letter codes) FAD, SAM, IAI and OQO (conformations from PDB IDs: 7bkc, 7c7m, 5sdv, 7v11). For each target, we generated 400 distinct diffusion trajectories, only providing the small molecule conformation and no privileged information about the backbone or any side chains in the native structure. For each generated backbone, we assign 8 sequences using LigandMPNN and predict all 8 sequences using AF2 in single sequence mode. We measure the RMSD of the AF2 predicted backbones compared to the design models to evaluate the probability of the backbone forming.

#### 9.3 Rosetta Calculations on Diffused Structures

GALigandDock was run in "eval" mode (as done in Section 7.7), to indicate that the ligand and any residues within a heavy atom contact distance of less than 8Å can move. We initialize the structure with the designed scaffold after the sequence has been assigned with LigandMPNN. LigandMPNN also predicts sidechain orientations which are used to initialize the sidechains for our energy calculations. The energy of the conformation is minimized using GALigandDock's generalized ligand potential and a harmonic constraint on the starting coordinates to prevent large movement. The pose is then scored using the generalized ligand potential in Rosetta.

#### 9.4 Rosetta Ligand-Aware Relax

Rosetta metrics such as "ddG" and "contact molecular surface" to computationally score protein and ligand binding were applied for the designs generated with RFdiffusionAA and following sequence design. The generated protein and ligand complex models were relaxed using Rosetta prior to scoring with the

Rosetta generic potential [32]. Harmonic potential on the protein Ca coordinates and the distances of selected pairs of protein backbone and ligand atoms were added as restraints to the generic scoring function. The estimated binding free energy “ddG” was calculated by taking the difference of *holo* and *apo* state Rosetta scores. The area of the target small-molecule packed by protein atoms was calculated using the Rosetta “contact molecular surface” metric [34]. This metric uses the contact distance to re-weight the contacting surfaces.

#### 9.5 Assessing Diversity of Designs

Diversity was assessed by generating 100 designs for the four *in silico* benchmark ligands using RFdiffusionAA without use of a potential. In addition, 100 designs were generated with RFdiffusion for the unconditional case as well as for scaffolding a 20 residue motif from PDB ID 5TRV used as a benchmark case in [33]. All designs were 150 residues in length. For each of the six design cases, 100x100 pairwise TMAligns were performed. Agglomerative clustering with the distance metric $1 - TMscore$ was performed using the complete linkage criterion for various distance thresholds using $scikit-learn$ [31]. This ensures that each member of any given cluster has a TMScore to every other member of the cluster at least as high as the clustering threshold, which was swept from 0 to 1 in 400 evenly spaced increments. The results are shown in Fig S10: left shifted curves correspond to higher diversity.

#### 9.6 Assessing Novelty of Designs

The 400 designs (without potential) for ligands FAD and SAM were TMAligned to the training dataset, and for each design, the highest scoring hit, the highest scoring hit with the same ligand, and the mean TMScore are recorded. The results are shown in figure S11. As expected, on average, designs are more similar to training data examples with the same ligand due to certain conserved binding modes (such as the Rossman fold for FAD). Yet, the distribution of Max TM-scores in S11.B shows that the designs are not merely memorized examples from the training dataset for the target ligand, as the maximum TM scores to the training set are higher than TM scores to the training set containing the same ligand. The median maximum TM score of designs to a training example containing the same ligand is 0.61 (FAD) and 0.62 (SAM). The other two molecules in our benchmark were not present in the training set.

#### 9.7 Bilin Binders

2776 designs across lengths 100, 150, and 200 were generated possessing the bilin-CARD motif at random locations at least 10 residues from either terminus. Half used a contact potential with linear decay. From those designs, 328 unique backbones were selected on the basis of having AF2 RMSD $< 2$, AF2 motif RMSD $< 1$, AF2 PAE $< 5$, Rosetta ddG $<$ -7.5, and Contact Molecular Surface $> 300$ [34]. These 328 designs were then clustered into 100 groups to maximize the minimum intra-cluster TMScore and the design with the lowest AF2 RMSD was selected from each cluster.

#### 9.8 Heme Binders

**Heme-substrate model preparation**

The heme model used for the design of proteins with an open substrate pocket represents a transition state of the C-H abstraction reaction by the ferryl oxygen from the methoxy group of anisole (i.e., anisole *O*-demethylase activity). While the subsequent design methods are not necessarily expected to yield heme enzymes with that particular activity, the chosen model represents a chemically sound heme-substrate complex with a generic reactant structure shape and size. An additional *para*-phenyl group was added to anisole to increase the likelihood of generating protein backbones with surface-accessible substrate binding pockets.

The heme model was built by locating the *para*-phenylanisole methoxy C-H abstraction transition state based on an axially methanethiolate-ligated (as a mimic for cysteine) heme ferryl intermediate in quartet state.All calculations were performed using Gaussian 16 software.[35] Structural optimizations and frequency calculations were performed with B3LYP-D3 method along with 6-31G(d) basis set and the SDD ECP on Fe atom. D3 dispersion correction was applied using the Becke-Johnson damping function.[36] Solvent effects of water were included using the CPCM solvation model during optimization. Frequency calculations were performed to confirm whether the structure is a minimum or a transition state. Intrinsic reaction coordinate (IRC) analysis was used to confirm that the obtained transition states connect the correct minima.

Conformer library of the transition state was created based on sampling the dihedral angles within the two propionic acid groups, the rotation of substrate above the heme plane, and the two ether C-O bonds. Conformational diversity of the transition states was sampled using a frozen coordinate conformer sampling script(`https://github.com/ikalvet/frozen-conf-xtb.git`) using the GFN2-XTB semiempirical QM method[37] for energy evaluation. 5000 conformers were saved.

The generated conformers were initially saved as XYZ files that were subsequently converted to MOLfiles using OpenBabel [12]. The bonding information in the MOLfile was manually inspected to ensure that the entire structure is represented as a single fragment, and edited, if necessary. Thereafter, mol2params.py script, available within Rosetta, was used to convert the MOLfile to a Rosetta-compatible .params file. The partial charges of the carboxylate oxygen atoms of the propionate groups were adjusted in the params files from -0.74 to -1.24 to increase the likelihood of H-bonds being created with these atoms during Rosetta design and relax.

**Ligand model selection for diffusion**

A subset of 55 conformers from the thousands of generated ligand conformers were used as inputs for RFdiffusionAA. The selected heme-substrate complex models are intended to serve the purpose of guiding the diffusion trajectories towards creating a heme binding pocket with more of a top-open substrate access, akin to unspecific peroxygenases and P450's. The conformers were selected based on three criteria:
1) Clustering the conformers based on structural similarity within 0.5 Å RMSD and selecting representative examples from each cluster (Supplementary Information Figure 2A).
2) Excluding conformers with the substrate pointed towards the carboxylate groups of heme. This was done to avoid creating an opening for a relatively hydrophobic substrate near possibly the most polar part of the heme binding site. Conformers with the C28-FE1-O5-C47 absolute dihedral angle greater than 50° were selected (Supplementary Information Figure 2B).
3) Excluding conformers where the substrate lies close to parallel against the heme plane. This was done to avoid creating heme binding sites with limited vertical space. Conformers with the FE1-O5-C47 angle greater than 140° were selected (Supplementary Information Figure 2C).

**Heme binding site design**

To generate input structures for RFdiffusionAA, the selected conformers were aligned to the HEM ligand in the crystal structures of cytochrome P450 peroxygenase CYP152K6 (PDB: 6fyj), and unspecific peroxygenase *Hsp*UPO from *Hypoxylon sp. EC38* (PDB: 7o2g). Four orientations of heme were sampled based on the rotation around the S-Fe bond at 90° intervals, and PDB files were created representing

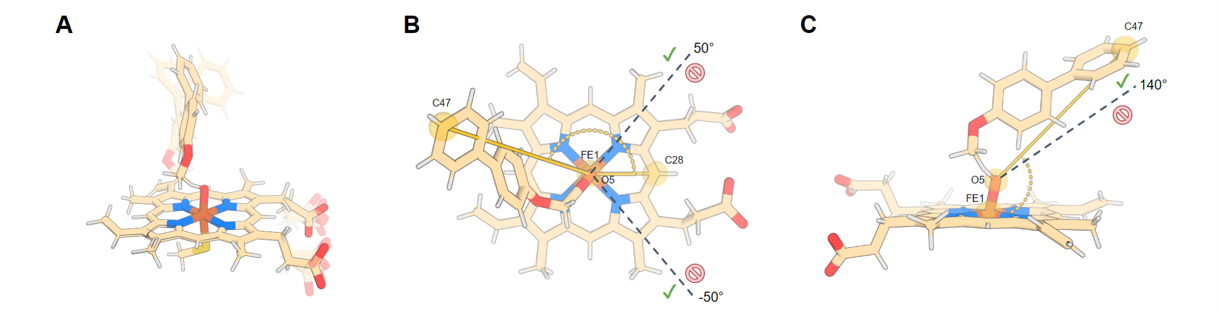

Supplementary Information Figure 2: Selection of heme-substrate complex conformers as inputs for RFdiffusionAA. A. Representative conformers picked from clusters of similar conformers. B. Excluding rotamers where the substrate molecule is close to the carboxylate groups of heme. C. Excluding rotamers where the substrate molecule lies too close to parallel with the heme plane.

each heme-substrate conformation in each of the sampled orientations. The heme-coordinating cysteine residues from these proteins were used as input motifs for RFdiffusionAA (CYS-15 for UPO and CYS-365 for P450). Furthermore, the selected motif residues were required to be outside of the first and the last 30 positions of any generated protein scaffolds. Protein backbones were generated with 150-210 amino acids in length. The resulting input contig string based on the UPO motif is "30-150,A15-15,30-150".

From each unique input conformer, 20-40 diffusion trajectories were spawned. A substrate contact potential was applied following either a quadratic or cubic decay to guide the trajectories towards creating proteins where the heme-binding site is sufficiently buried. With no ligand contact potential applied, a vast majority of diffused protein backbones lacked a well-buried heme binding site. On the other hand, using linear decay of the guiding potential led to significant clashes between the ligand and the scaffold. A total of ~10000 backbones were generated through the combination of the different input motifs, conformers and contact potentials. From the generated protein backbones we filtered out globular proteins with desirable binding pocket qualities based on features describing ligand burial, substrate exposedness, lack of clashes between the ligand and protein backbone, loop content, distance of termini from the ligand, radius of gyration, and the length of the longest helix.

As part of the scaffold generation process, we also explored further diversification of the selected backbones using partial diffusion (fixed length global backbone remodeling)[38] or RoseTTAFold joint inpainting (variable length loop remodeling)[39]. Partial diffusion allows subsampling the structural space roughly defined by the input scaffold. With RoseTTAFold joint inpainting (RF*joint*) we re-generated loop regions of the diffused backbones while allowing variable length loop substitutions. The loop regions were selected based on geometric distance from the ligand heavyatoms and the motif residue. Sequence design was enabled on all non-motif positions during the inpainting process. For each input scaffold, 5 diversified backbones were generated with both of these methods.

ProteinMPNN[40] was thereafter used to generate amino acid sequences optimal for each of the selected backbones. With the sequence design we reasoned that due to a significant degree of flexibility in the heme-substrate complex it would be beneficial to first identify sequences that are more likely to fold correctly before focusing on designing the protein-ligand interactions. Therefore, we then used ProteinMPNN to generate 15 sequences for each of the backbones while keeping the coordinating CYS fixed. Three sampling temperatures (0.1, 0.2, 0.3) were used, with 5 sequences generated at each, and introduction of additional cysteines was disallowed. Generated sequences were analyzed using single-sequence AlphaFold2 prediction (model 4 with 3 recycles) to identify the sequences that do fold into the desired shape. Successful sequences were selected based on AF2 metrics of plDDT $>$ 87.0, C$\alpha$

RMSD $<$ 1.5 Å, and CYS sidechain RMSD $<$ 1.2 Å. The selected backbones were then subjected to ligand binding site design by using either Rosetta FastDesign[22], or iterative application of ligandMPNN and Rosetta FastRelax[41]. The designable positions were selected based on distance from the heme-substrate complex heavyatoms, with residues with C$\alpha$ atom within 8 Å from any ligand heavyatom considered for design. Geometric constraints were applied to the heme-CYS interaction using the Rosetta AddOrRemoveMatchCsts mover. Successful designs were selected based on metrics describing protein-ligand contacts (Rosetta ddG, contact molecular surface, SASA, H-bonds to acceptors). An additional round of ligandMPNN sequence design was performed at the positions flanking the binding pocket, with only those in the window of 8-12 Å from the ligand enabled for redesign. The structures of the resulting sequences were predicted using AlphaFold2 (with tighter cutoffs of C$\alpha$ RMSD $<$ 1.2 Å and CYS sidechain RMSD $<$ 1.0 Å applied). Finally, the heme-substrate model was re-aligned into the AF2-predicted model structure, and relaxed with Rosetta FastRelax. The relaxed models were re-evaluated based on the same protein-ligand interaction metrics as used before. Upon final manual inspection of the generated designs, 168 were selected for experimental testing.

#### 9.9 Digoxigenin Binders

To demonstrate the capability of RFdiffusionAA in designing *de novo* binders for small molecules with the ligand information alone, we designed digoxigenin-binding proteins. We used a model conformation of the target ligand digoxigenin as an input to generate 25,000 diffused backbones using RFdiffusionAA. The diffusion outputs were subsequently filtered based on the radius of gyration, secondary structure content, solvent accessible surface area (SASA) of the small molecule, and the number of contacts between the protein backbone and ligand atoms. The selected backbones underwent ligand pose-aware sequence design by applying iterative cycles of LigandMPNN and Rosetta FastRelax ([22], [41]). Eight sequences were sampled per backbone with LigandMPNN, and AF2 was used to predict the protein structures. The selected designs with high AF2 pLDDT ($>$80) and low backbone-RMSD ($<$1.5Å) were used to re-dock the ligand digoxigenin using ChemNet. We reasoned that by employing ChemNet, a deep-learning fixed backbone ligand docking method, updated ligand positions would contribute in sampling more sequences from LigandMPNN. We used the docking outputs with high ChemNet confidence ($>$80 ChemNet pLDDT; to filter out poor predictions from ChemNet) to perform a second round of sequence design with LigandMPNN. Finally, we conducted filtering of the designs based on AlphaFold2 and Rosetta metrics. The designs chosen for experimental testing exhibited high AF2 pLDDT ($>$80), low backbone-RMSD between design model and AF2 prediction ($<$2 Å), low Rosetta ddG ($<$-30), and formed at least one hydrogen bond with the ligand. Given that our experimental screen involved digoxigenin covalently linked to biotin, we also excluded designs where the digoxigenin-linker atoms were deeply buried within the protein (Rosetta atomic_depth $<$ 30). This resulted in 4,416 designs to experimentally screen.

### 10 Experimental Methods

#### 10.1 Bilin Binders

Synthetic genes encoding the RF Diffusion designs were ordered as eBlocks gene fragments in a 96-well plate format (Integrated DNA Technologies) and ligated into modified pET-29b as in [[40]]. The resulting plasmid arrays were transformed into *E. coli* BL21(DE3) harbouring pCOLADuet-*cpcEF*-*pebS*-HO1 ([42]). This plasmid encodes the biosynthetic machinery required to synthesize phycoerythrobilin (PEB) as well as a bilin-protein lyase, CpcEF, which is required to attach PEB to the CXRD bilin binding motif included in all the protein designs. PEB-binding positives in the 96-well plate were easily identified by their pink color, and were further characterized by scanning the plate using UV-Vis absorption (250-800 nm - Fluostar Omega) and fluorescence (excitation: 460, 520, and 630 nm; Emission filters: 530BP30, 605DF50, and 695DF55 - Amersham Imager 600).

Positives from the initial screening were scaled up in 500 ml LB broth cultures in 2L baffled Erlenmeyer flasks at 37°C and 180 rpm in the presence of 50 $\mu$g/ml and 100 $\mu$g/ml, kanamycin and ampicillin, respectively. At an optical density at 600 nm ($OD_{600}$) of $\sim$ 0.6, IPTG was added to a final concentration of 1 mM to induce protein production and the cultures were switched to 18°C for 16 h. *E. coli* cell pellets were resuspended in 50 mM HEPES, 500 mM NaCl, 20 mM Imidazole (pH 7.6), disrupted by sonication, and the His-tagged biliproteins were purified from the cell extracts via immobilized metal affinity chromatography (IMAC) and subsequent size-exclusion chromatography (SEC).

Samples were diluted to OD $\sim$ 0.1 for recording fluorescence emission spectra (Horiba Fluorolog-3), using a 10 nm FWHM excitation source. An absorptance (1-T) spectrum was calculated for each sample (Agilent Cary 60), and the ratio of absorptance to fluorescence (areas under the graph) were used to estimate the relative fluorescence yield for each biliprotein, with CpcA-PEB arbitrarily set to 100%.

#### 10.2 Heme Binders

**Construction of pET29b(+) plasmids encoding heme binding protein variants**

Double-stranded DNA fragments encoding the designs and any variants thereof (codon-optimized for bacterial expression) were purchased from Integrated DNA Technologies (IDT) as eBlocks™ Gene Fragments. Following the Golden Gate cloning protocol using T4 DNA ligase and BsaI-HFv2 restriction enzyme (master mix #E1601, NEB),[43] the DNA fragments encoding design sequences and including overhangs suitable for a BsaI restriction digest were cloned into a custom pET29b(+) target vector containing lethal ccdb gene, and C-terminal SNAC[44] and hexahistidine tags (#191551, Addgene). This yielded final expressed sequences as: MSG <design> GSGSHHWGSTHHHHHH.

**Small-scale screen for expression**

Assembled plasmids containing the designs were transformed into chemically competent *E. coli* BL21(DE3) cells by heat shock. DNA was incubated on ice with competent cells for 30 minutes, followed by 30 second heat shock at 42 °C, and 2 minute incubation on ice. 100 $\mu$L rich medium (super optimal broth with catabolite repression, SOC) was added to transformed cells and samples were incubated at 37 °C, 1050 r.p.m. on a shaking platform for 1 hour. The cells were subsequently transferred to 900 $\mu$L of LB medium containing 50 $\mu$g/mL kanamycin, and incubated on a shaking platform (1050 r.p.m) at 37 °C for 16 hours. Thereafter, 100 $\mu$L of the starter culture was transferred to 900 $\mu$L of TB-II medium containing 50 $\mu$g/mL kanamycin and the cultures were grown at 37 °C for 4 hours. Protein expression was induced by the addition of 2 mM IPTG and the cultures were incubated at 37 °C for 2 hours. Meanwhile, glycerol stocks were prepared by mixing 100 $\mu$L of the starter culture in LB media with 100

$\mu$L of 50% glycerol and stored at -80 °C. The cell pellets were harvested by centrifugation at 4,000 g for 10 minutes and lysed with BugBuster® lysis reagent containing 0.01 mg/mL deoxyribonuclease and 0.1 mg/mL lysozyme, using a ultrasonication in a microplate horn (40% amplitude with 10 s on, 10 s off for a total of 4 min on-time). Lysate was collected by centrifugation at 4,000 xg for 20 minutes and analyzed by SDS-PAGE to identify solubly expressed protein variants. The variants not showing a band of over-expressed protein on the SDS-PAGE gel may have done so for a number of reasons: very low level of expression; insoluble protein; errors in the cloned DNA fragment. The precise modes of failure were not investigated at this stage of screening.

**Small-scale heme-binding screen**

To clarified cell lysates containing overexpressed protein was added 9 $\mu$L of hemin solution (250 $\mu$M in 0.5 M aq. NaOH) to reach a final hemin concentration of 10 $\mu$M. The lysates were then applied to Ni-NTA resin (50 $\mu$L) that was equilibrated with wash buffer (50 mM KPi, 200 mM NaCl, 25 mM imidazole, pH 7.4). The resin was washed with 25 column volumes (CV) of wash buffer. Protein was eluted with 200 $\mu$L of elution buffer (50 mM KPi, 200 mM NaCl, 300 mM imidazole, pH 8.0). To remove non-specifically bound heme and most of the imidazole, 130 $\mu$L of the eluted protein solutions were thereafter loaded onto a 96-well PD MultiTrap G-25 desalting plate (Cytiva) equilibrated with a buffer containing 50 mM KPi and 200 mM NaCl at pH 7.2, and eluted by centrifugation at 800 xg for 2 minutes. As a control, the IMAC elution buffer alone was also eluted through the desalting column, and the resulting solution used as a background in the subsequent UV-Vis spectroscopic analysis. UV-Vis absorbance spectra of the desalted heme-loaded protein solutions were collected in half-area UV-STAR microplates (Greiner) using a platereader (BioTek Synergy Neo2) in the 250-700 nm range. Heme-binding was qualitatively assessed based on the wavelength and the intensity of the Soret maximum of heme. A Soret maximum at 420-425 nm is indicative of CYS-ligated heme-binding in a hexacoordinate low-spin state (with the 6th coordination site most likely filled by endogenous imidazole), whereas a Soret maximum at 370-390 nm, along with a charge transfer band at  640 nm is indicative of a CYS-ligated pentacoordinate high-spin heme state.[45] The collected spectra are presented in Supplementary Information Figures 3, 4 and 5.

**Larger-scale expression and purification of heme-binding proteins**

45 designs were selected based on the results of the small-scale heme-binding assay for expression scale-up and further characterization. Clonal variants of the designs were obtained by spreading stabs from the polyclonal glycerol stocks on LB-agar plates containing 100 $\mu$g/mL kanamycin and incubating the plates at 37 °C for 16 hours. Single colonies were picked, and the DNA fragments encoding the designs were amplified following a colonyPCR protocol using GoTaq® Green DNA polymerase master mix (#M7122; Promega) and T7 reverse and forward primers. The PCR products identified to contain DNA of appropriate size based on agarose gel (1.2%) electrophoresis with SybrSafe dye were sent to Sanger sequencing (GeneWiz/Azenta) for sequence-verification. Single colonies containing the correct design sequences were grown up in 5 mL LB media containing 50 $\mu$g/mL kanamycin, over 16 hours at 37 °C. 2 mL of the starter culture was used to inoculate 40 mL TB-II media containing 50 $\mu$g/mL kanamycin and the rest used for plasmid extraction following the Qiagen QIAprep MiniPrep protocol. The 40 mL cultures were grown at 37 °C for 4 hours, after which protein expression was induced with the addition of 1 mM IPTG, and the cultures were incubated at 37 °C for 2 hours. Pellets were harvested by centrifugation at 4,198 g for 8 minutes and resuspended in a lysis buffer containing 50 mM KPi, 200 mM NaCl, 25 mM imidazole, 0.01 mg/mL DNAse, 0.1 mg/mL lysozyme, and a Pierce protease inhibitor tablet. 200 $\mu$M hemin (from 12 mM stock in 0.5 M aq. NaOH) was added to the resuspended cells. Lysis was immediately performed by ultrasonication (13 mm probe, 2.5 mins, 10s on, 10s off, 65% amplitude). Lysate was collected by centrifugation at 15,000 xg for 20 minutes and applied to Ni-NTA resin that

was equilibrated with wash buffer (50 mM KPi, 200 mM NaCl, 25 mM imidazole, pH 7.4). The resin was washed extensively with 50 column volumes (CV) of wash buffer. Protein was eluted with 1.2 CV of elution buffer (50 mM KPi, 200 mM NaCl, 300 mM imidazole, pH 8.0) and further purified via size exclusion chromatography (SEC) using a Superdex Increase 75 10/300 GL column (GE Healthcare) on ÄKTAxpress (GE Healthcare) instrument at 0.8 mL min-1 flow rate using a running buffer containing 50 mM KPi, 200 mM NaCl pH 7.2. The monomeric or smallest oligomeric fractions of each run were collected. The obtained chromatograms are presented in Supplementary Information Figures 6 and 7. The Cys/Ala knockout mutants of the designs were produced by following the aforementioned Golden Gate assembly protocol, and transformed into *E. coli* BL21(DE3) cells as described above. After incubation in SOC media, the cells were spread on LB-agar plates containing 100 $\mu$g/mL kanamycin, incubated at 37 °C for 16 hours. Single colonies were picked and sequence-verified, and the larger-scale expression protocol was followed in most part. These proteins were purified without the addition of hemin prior to the lysis step, and were isolated as apo proteins.

**Mass spectrometry analysis**

MS data for the designed proteins were acquired on an Agilent 1200series LC G6230B TOF LC-MS with an AdvanceBio RP-Desalting column (A: H2O with 0.1% Formic Acid, B: Acetonitrile with 0.1% Formic Acid). The final protein concentrations were adjusted to 1-2 mg/mL in 50 mM KPi, 200 mM NaCl, pH 7.2. Subsequent data deconvolution was performed in Bioconfirm using a total entropy algorithm. All data are presented in Table 15.

**Variable temperature spectrophotometric measurements**

To observe changes in the spectral properties of bound heme at increasing temperatures, UV/Vis spectra were measured of in vitro loaded holo-proteins using the Jasco Spec V750 spectrophotometer and a 10 mm pathlength cuvette. Spectra in the 230-700 nm range were collected at every 10 °C intervals between 25 °C and 95 °C. Temperature was increased at the rate of 5 °C min$^{-1}$, and spectra were acquired after the temperature had stabilized to within 0.5 °C of target temperature for 5 seconds. Measurements were performed with 20 $\mu$M solutions of purified holoprotein in KPi buffer (50 mM KPi, 200 mM NaCl, pH 7.2).

**Circular dichroism spectroscopy**

To determine secondary structure and thermostability of the designs, far-ultraviolet circular dichroism (CD) measurements were carried out on a JASCO J-1500 instrument using a 1 mm pathlength cuvette. Samples of purified protein were prepared at 0.3-1.0 mg/mL in 50 mM KPi, 20 mM NaCl, pH 7.2. The temperature of the sample was scanned from 25 °C to 95 °C with full spectrum scans from 190 nm to 260 nm performed after each 10 degree increment. Protein concentrations were determined by absorbance at 280 nm, measured using a NanoDrop spectrophotometer (Thermo Scientific) using predicted extinction coefficients.[46]

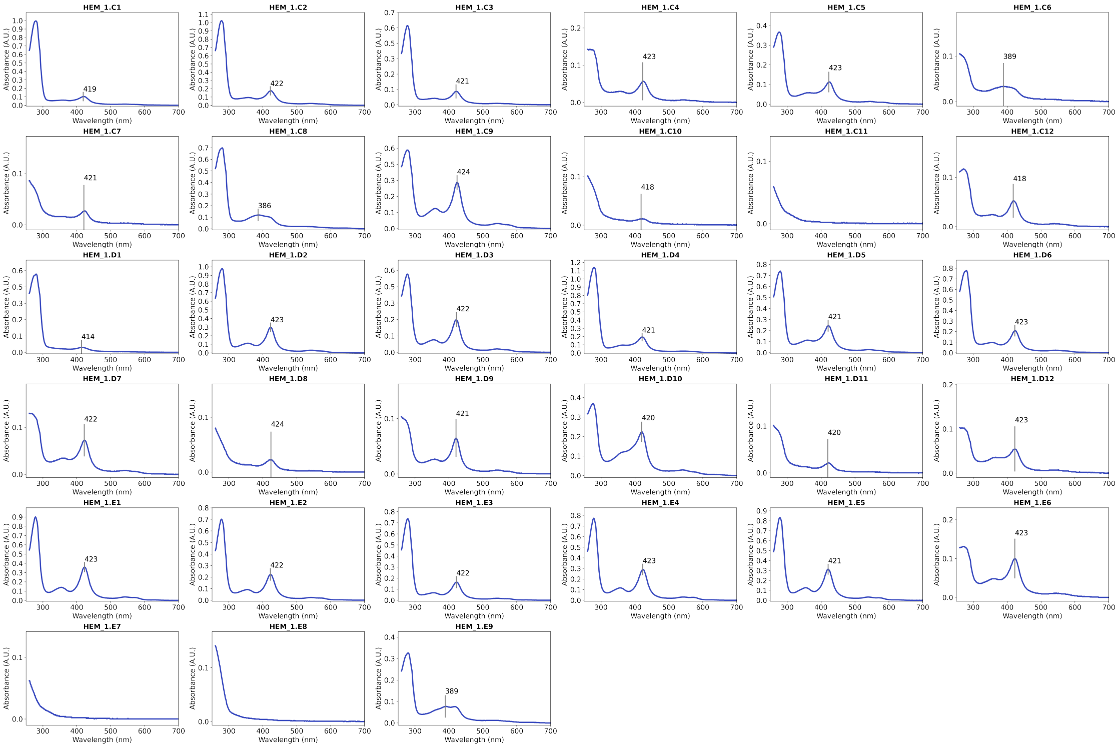

Supplementary Information Figure 3: UVVis spectra collected from small-scale screening for heme binding, HEM_1.C1 to HEM_1.E9. Samples were prepared by adding 10 $\mu$M hemin to clarified cell lysates (grown from 1 mL cell cultures), purifying the proteins with Ni-NTA affinity chromatography, and eluting through PD MultiTrap G-25 desalting column.

#### 10.3 Digoxigenin Binders

The yeast transformation was performed by ordering the selected designs as synthetic oligonucleotides and transforming into *S. cerevisiae* EBY100 strain as a pooled library, using a previously described protocol [34]. EBY100 cultures were grown in C-Trp-Ura medium supplemented with 2% (w/v) glucose (CTUG). For induction of expression, yeast cells initially grown in CTUG were transferred to SGCAA medium supplemented with 0.2% (w/v) glucose and induced at 30 °C for 16–24 h. Cells were washed with PBSF (PBS with 0.1% (w/v) BSA) and incubated with a solution containing 2 $\mu$M of biotinylated digoxigenin, streptavidin conjugated to PE (SA-PE, Invitrogen), and anti-myc antibody conjugated to FITC (Immunology Consultants Laboratory) for 40 minutes at room temperature. After incubation time, cells were washed with PBSF and resuspended before cell sorting. We performed fluorescent activated cell sorting (FACS) to collect cells with PE-signal which represents binding to biotinylated digoxigenin. We performed a second round of cell sorting with three different conditions of incubation, which were prepared by having different biotinylated digoxigenin concentrations. We identified three designs that showed enrichment for binding to biotinylated digoxigenin and SA-PE. To characterize the binding affinity of the designs *in vitro*, we used Golden Gate assembly reaction with BsaI-HFv2 restrction enzyme (NEB) to clone the gene fragments of the potential hit sequences into a custom pET29b(+) target vector including a BsaI restriction site, lethal ccdb gene, and N terminal histidine tag for protein expression in *E. coli* [43]. This resulted in the final expressed sequence being MSHHHHHHSG-design-GS. We transformed the cloned plasmid into BL21(DE3) competent cells (NEB), and the *E. coli* cells were incubated for growth and expression in autoinduction medium for 16 hrs at 37 °C. The expressed proteins were purified using His-tag and Nickel affinity with IMAC, and further purification was achieved by using size-exclusion chromatography (SEC) in phoshpate buffered saline with 137 mM NaCl, 2.7 mM

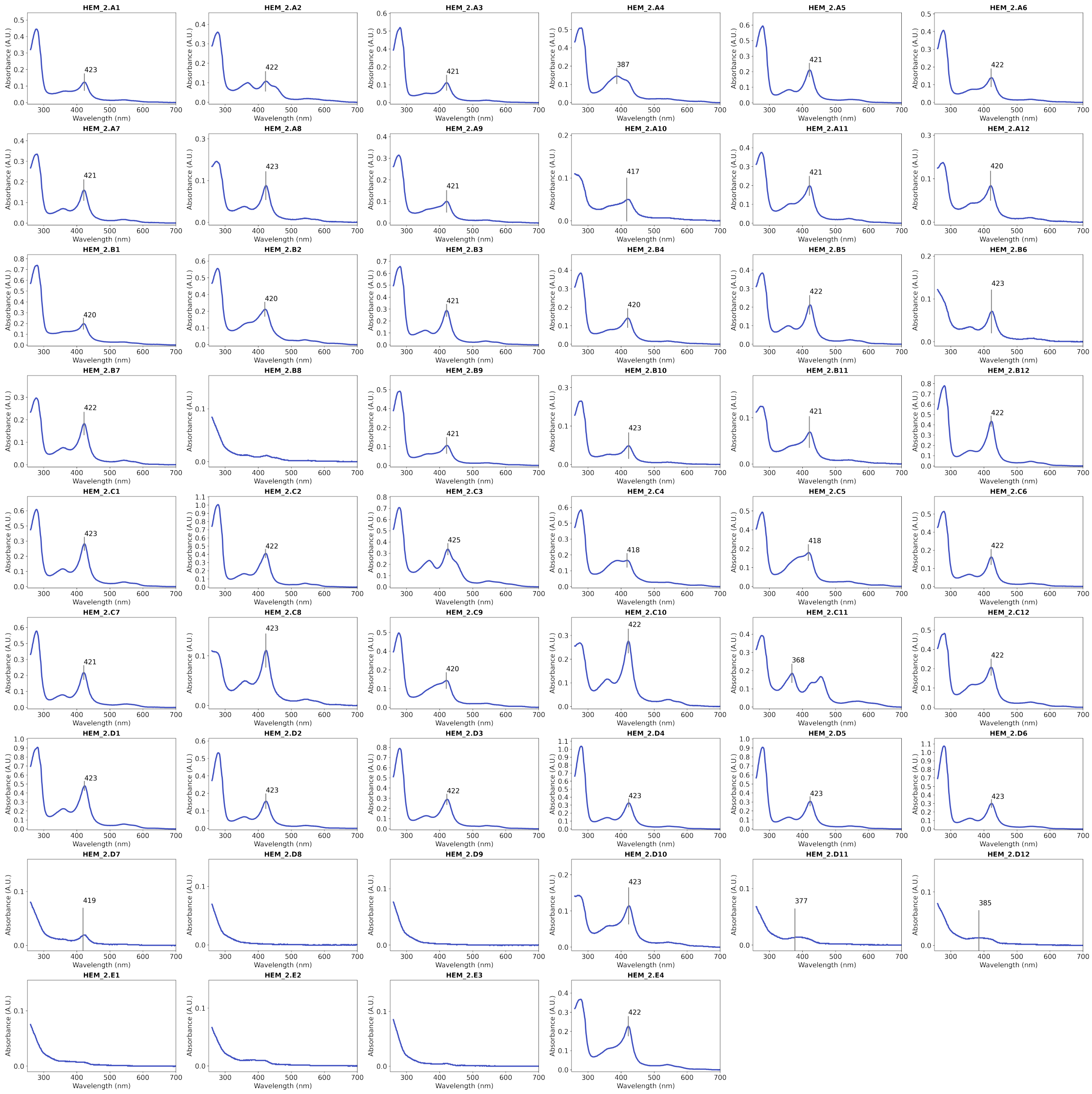

Supplementary Information Figure 4: UVVis spectra collected from small-scale screening for heme binding, HEM_2.A1 to HEM_2.E4. Samples were prepared by adding 10 $\mu$M hemin to clarified cell lysates (grown from 1 mL cell cultures), purifying the proteins with Ni-NTA affinity chromatography, and eluting through PD MultiTrap G-25 desalting column.

KCl and 11.9 mM phosphates (PBS, Fisher). Superdex Increase 75 10/300 GL column (GE Healthcare) was used with ÄKTAxpress (GE Healthcare) for SEC and we collected monodisperse fractions to further determine the binding affinity to digoxigenin. Fluorescence polarization (FP) experiment was performed by measuring fluorescence polarization with decreasing protein concentration when the concentration of AlexaFluor488-labeled digoxigenin was fixed as 5nM in phosphate buffered saline (PBS) as done previously [47]. Isothermal titration calorimetry (ITC) was carried out by injecting 140.8 $\mu$M label-free digoxigenin to 27.3 $\mu$M binder protein in PBS with 0.5% DMSO using the 19 injections with rinsing protocol of MicroCal PEAQ-ITC (Malvern Panalytical). ITC curve shown in Supplementary Information Figure 12.

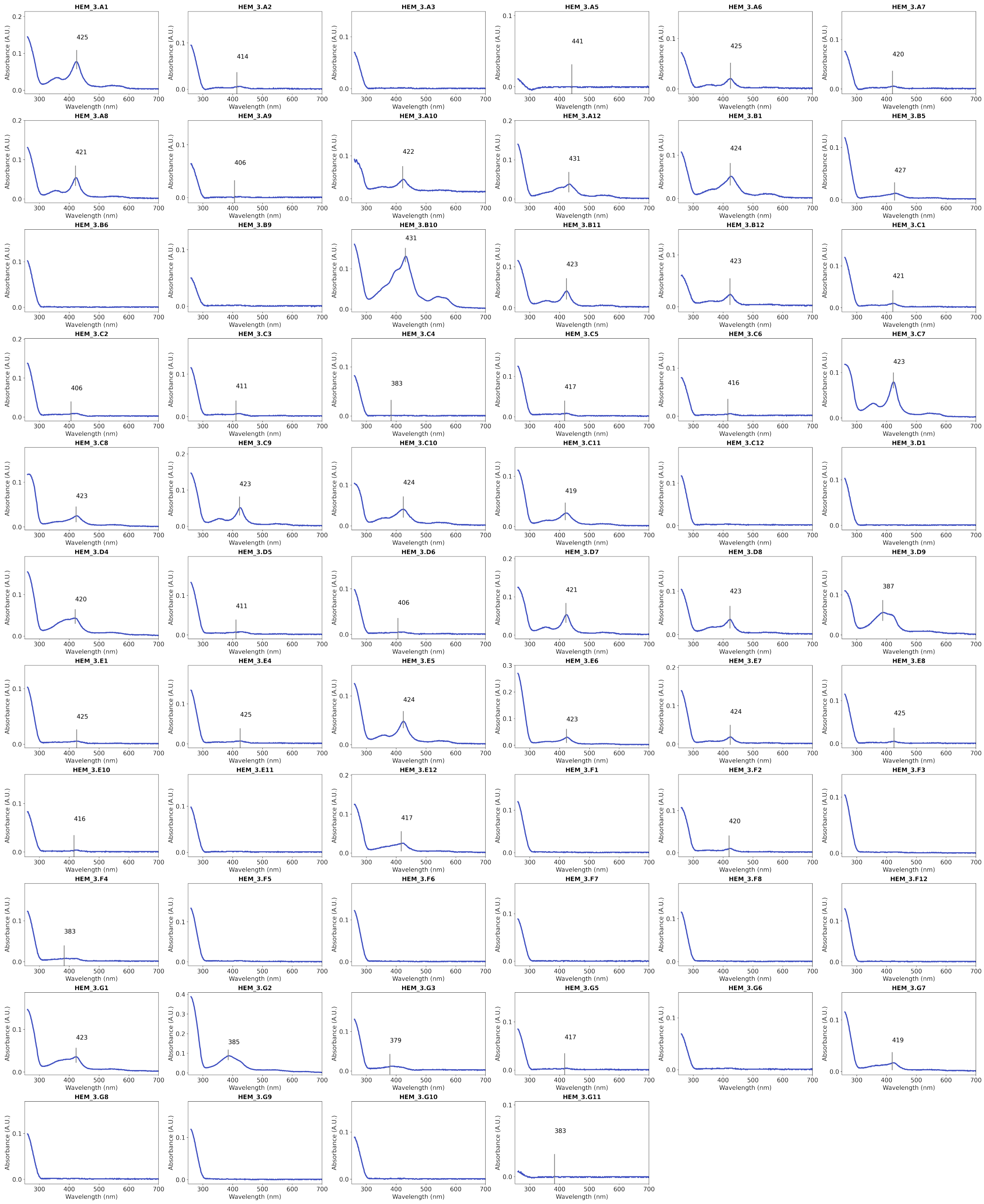

Supplementary Information Figure 5: UVVis spectra collected from small-scale screening for heme binding HEM_3.A1 to HEM_3.G11. Samples were prepared by adding 10 $\mu$M hemin to clarified cell lysates (grown from 1 mL cell cultures), purifying the proteins with Ni-NTA affinity chromatography, and eluting through PD MultiTrap G-25 desalting column.

### 11 Figures and Statistics

Figures were generated using matplotlib[48] and seaborn[49] and appropriate statistical tests were performed with Scipy[50] when noted in figure legends. Outliers were removed from boxplots for clarity.

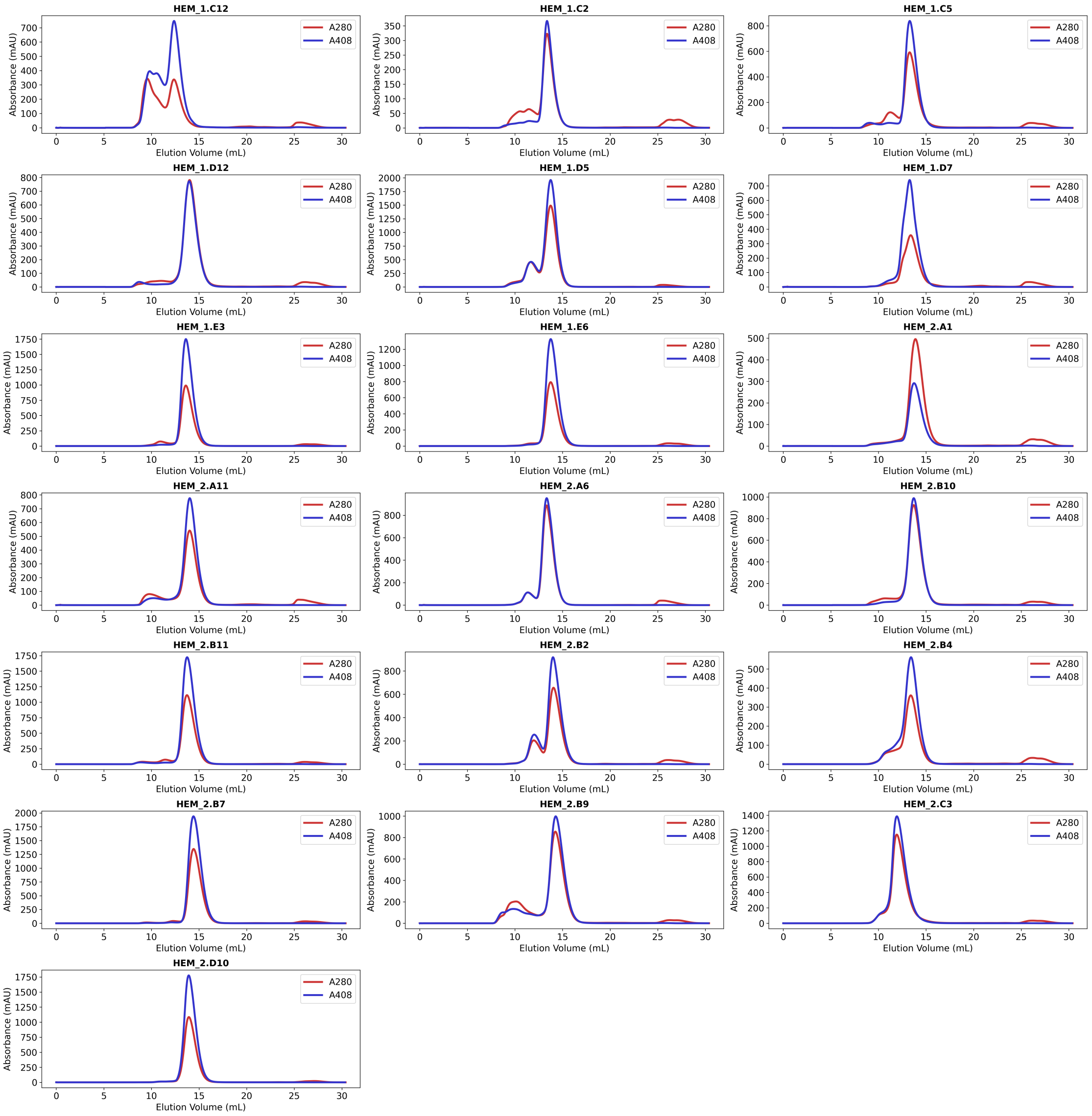

Supplementary Information Figure 6: Size-exclusion chromatograms of heme-loaded proteins, HEM_1.C2 to HEM_2.D10. Data were collected using a Superdex Increase 75 10300 GL column (GE Healthcare) in a buffer containing 50 mM KPi and 200 mM NaCl at pH 7.2. Void volume of the column is 8.5 mL. Blue chromatograms were obtained by following the absorbance at 408 nm, indicating elution of heme-containing species. Red chromatograms were obtained from absorbance at 280 nm.

### 12 Supplementary Results

#### 12.1 Ligand-Aware Protein Structure Prediction

In order to investigate whether or not the additional ligand context helps the RFAA model make better predictions of protein structure (Fig. S3), we filtered the dataset from Section 7.3.2 down to only those items that RFAA predicts confidently (inter-chain PAE $< 10.0$). We hypothesize that in the cases that RFAA gets the ligand dock correct, it may predict more accurate structures of the protein binding pocket. In those cases in which RFAA fails to dock the ligand correctly, it is unlikely that the added ligand context will aid in protein structure prediction.

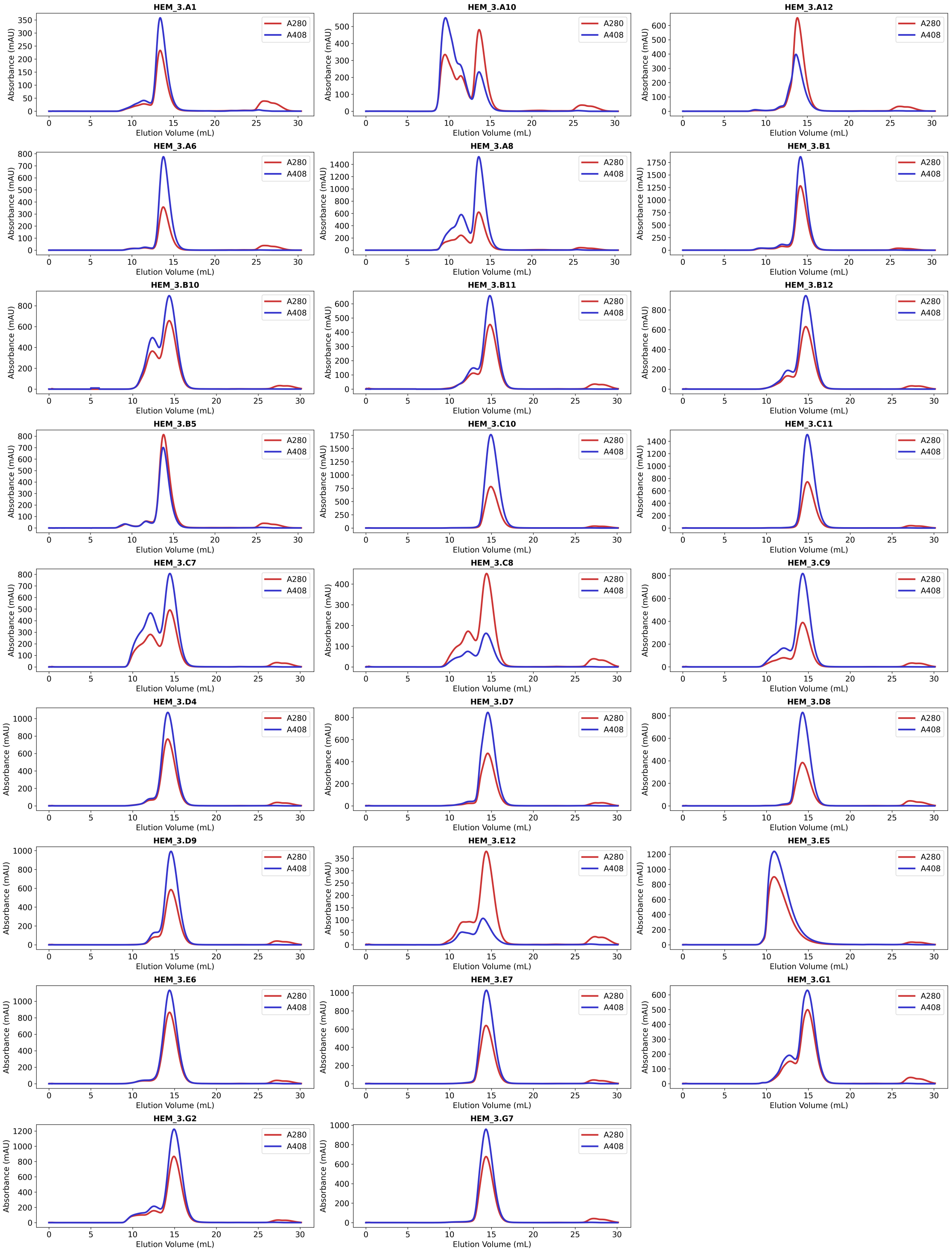

Supplementary Information Figure 7: Size-exclusion chromatograms of heme-loaded proteins, HEM_3.A1 to HEM_3.G7. Data were collected using a Superdex Increase 75 10300 GL column (GE Healthcare) in a buffer containing 50 mM KPi and 200 mM NaCl at pH 7.2. Void volume of the column is 8.5 mL. Blue chromatograms were obtained by following the absorbance at 408 nm, indicating elution of heme-containing species. Red chromatograms were obtained from absorbance at 280 nm.

##### 12.1.1 Comparing against RF2

We predict the same protein chains (those that RFAA predicts with high confidence) with RF2, which has no way to represent the bound ligand. This a safe comparison because the validation clusters between

Supplementary Methods Table 15: Mass spectrometry data for diffused heme-binding proteins. In all cases the observed mass corresponded to the loss of N-terminal methionine (-130 Da).

| **Variant** | **Expected Mass** | **Observed Mass** | | **Variant** | **Expected Mass** | **Observed Mass** |
|---|---|---|---|---|---|---|
| HEM_1.C12 | 20524 | 20393 | | HEM_3.A8 | 25128 | 24997 |
| HEM_1.C2 | 27185 | 27054 | | HEM_3.B1 | 19349 | 19218 |
| HEM_1.C5 | 26735 | 26604 | | HEM_3.B10 | 23298 | 23168 |
| HEM_1.D12 | 21959 | 21828 | | HEM_3.B11 | 21344 | 21213 |
| HEM_1.D5 | 20016 | 19885 | | HEM_3.B12 | 21761 | 21630 |
| HEM_1.D7 | 22406 | 22275 | | HEM_3.B5 | 24682 | 24551 |
| HEM_1.E3 | 22940 | 22809 | | HEM_3.C10 | 23580 | 23449 |
| HEM_1.E6 | 22722 | 22591 | | HEM_3.C11 | 23542 | 23411 |
| HEM_2.A1 | 23983 | 23852 | | HEM_3.C7 | 23187 | 23056 |
| HEM_2.A11 | 22449 | 22319 | | HEM_3.C8 | 23218 | 23087 |
| HEM_2.A6 | 23630 | 23499 | | HEM_3.C9 | 23179 | 23048 |
| HEM_2.B10 | 17123 | 16992 | | HEM_3.D4 | 21383 | 21252 |
| HEM_2.B11 | 22889 | 22758 | | HEM_3.D7 | 24155 | 24024 |
| HEM_2.B2 | 22750 | 22619 | | HEM_3.D8 | 21197 | 21066 |
| HEM_2.B4 | 22449 | 22318 | | HEM_3.D9 | 21326 | 21195 |
| HEM_2.B7 | 17507 | 17377 | | HEM_3.E12 | 23057 | 22926 |
| HEM_2.B9 | 16938 | 16807 | | HEM_3.E5 | 21084 | 20953 |
| HEM_2.C3 | 19173 | 19042 | | HEM_3.E6 | 23364 | 23233 |
| HEM_2.D10 | 22007 | 21876 | | HEM_3.E7 | 23226 | 23095 |
| HEM_3.A1 | 24975 | 24844 | | HEM_3.G1 | 22702 | 22571 |
| HEM_3.A10 | 24457 | 24326 | | HEM_3.G2 | 23108 | 22977 |
| HEM_3.A12 | 21607 | 21476 | | HEM_3.G7 | 22482 | 22351 |
| HEM_3.A6 | 24773 | 24642 | | | | |

RFAA and RF2 are identical so the only large differences are the architectural differences described above and the ability to explicitly model ligands. We expect that comparisons to other structure prediction networks will be harder because of training dataset bias. We further filter the evaluation set by confident prediction from RF2 (protein pLDDT > 80), and end up with 594 unique items in an evaluation set that are predicted confidently by both RFAA and RF2. For each item, we make 3 predictions from both RFAA and RF2 to remove any artifacts that come from random seed, and pick the prediction that either minimizes inter-chain PAE or maximizes pLDDT, respectively.

We measure the all-atom RMSD of the predicted protein structure relative to the crystal after kabsch alignment on the backbone atoms. We also measure the *ligand pocket* RMSD, where the ligand pocket is defined as the set of residues that have at least one atom within 5Å of the ligand in the crystal structure. The ligand pocket RMSD is then computed as the all-atom RMSD of said residues after kabsch alignment on the backbone atoms of the same residues.

We observe a statistically significant difference (paired t-test) between the RMSD values from RFAA and RF2 (Fig. S3A). We depict several examples of structures that RFAA predicts better than RF2, likely due to the added ligand context (Fig. S3B-D). One illustrative example is the PDB entry 7rjj in our test set. For this target, RFAA generates an accurate prediction while RF2 predicts an incorrect, "open" conformation of a helix forming the ligand-binding pocket (Fig. S3D, pink structure). This open conformation is present in the most sequence-similar example in the training set (Fig. S3D, yellow structure). However, a different example in the training set, which has lower sequence similarity to 7rjj, has the helix in a "closed" conformation in the presence of a ligand (Fig. S3D, orange structure). We hypothesize that RFAA uses the presence of the ligand to better disambiguate alternate conformations

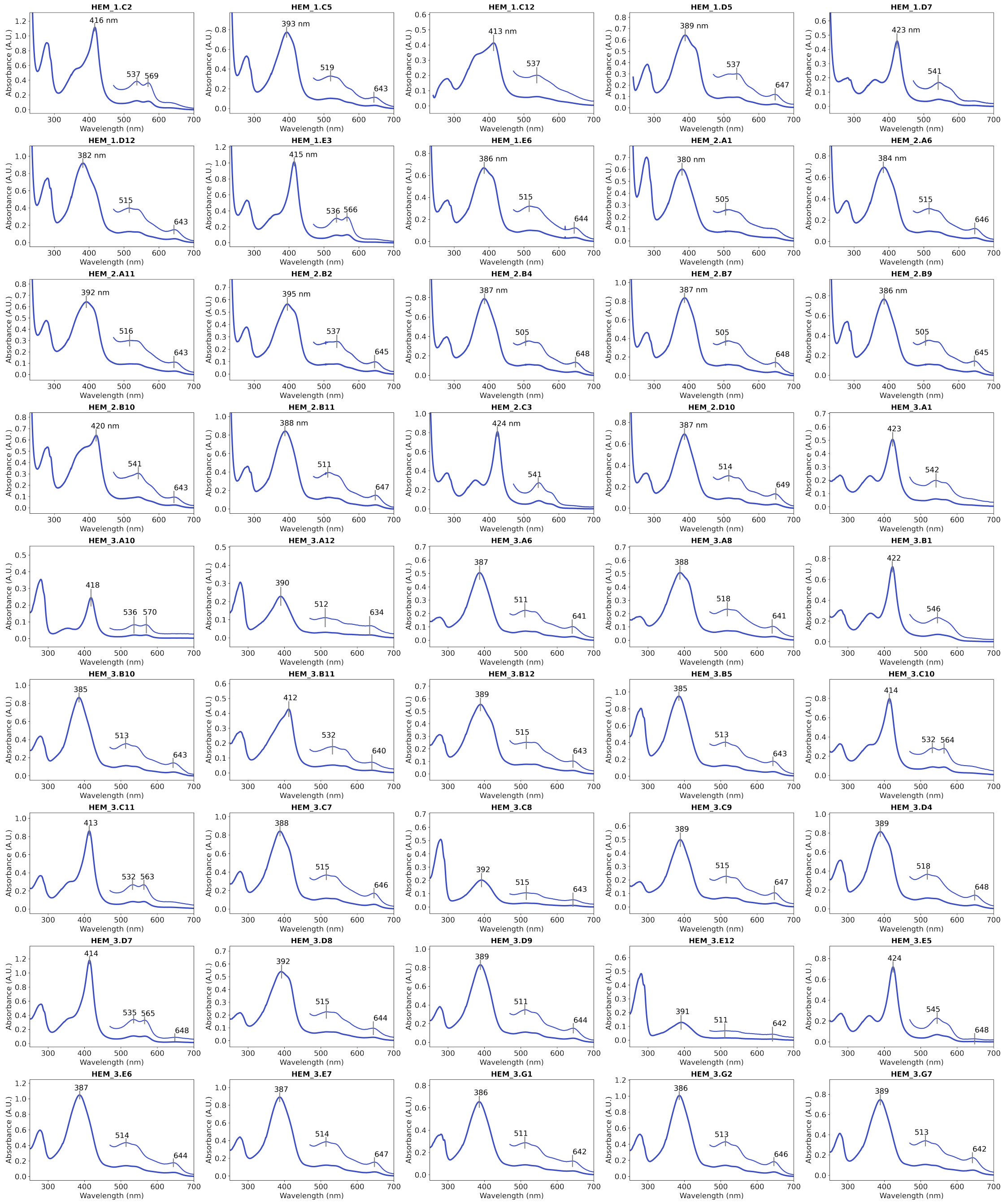

Supplementary Information Figure 8: UVVis spectra of heme-loaded proteins. Inset shows the visible region at 3x magnification. Spectroscopic data of most designs is in agreement with CYS-ligated heme binding (either Soret maximum at ∼420 nm and Q band features at 540570 nm for hexacoordinate low spin state, or Soret maximum at 370-390 nm and Q band features at 510/540 and charge transfer band at 640 nm for pentacoordinate high spin state). Spectra were recorded in a buffer containing 50 mM KPi and 200 mM NaCl at pH 7.2.

of similar proteins seen during training.

##### 12.1.2 Predicting Structures With and Without Ligands

A natural question that arises is whether or not RFAA can predict conformational shifts in a protein with and without a ligand partner present. For those same set of 594 items that are confidently predicted by RFAA and RF2, we make predictions of the protein structure without the added ligand context

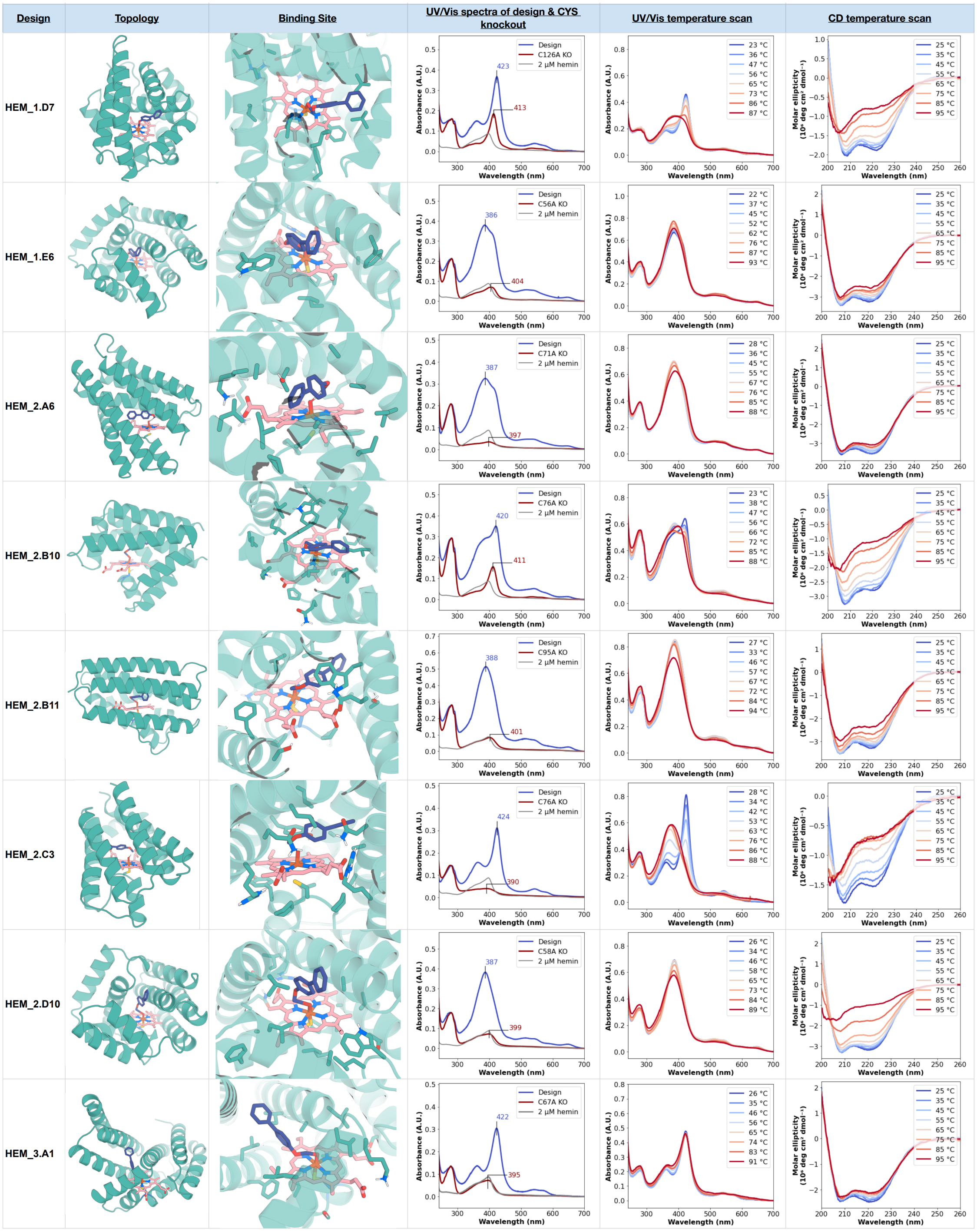

Supplementary Information Figure 9: Characterization of heme-binding proteins obtained with RFdiffusionAA, HEM_1.D7 to HEM_3.A1. The 4th column shows the UVVis spectra of the purified heme-loaded protein (blue), and the putative axial CysAla mutant at 10 $\mu$M, mixed with 2 $\mu$M hemin (red trace), along with free hemin at 2 $\mu$M (gray). The 5th column shows changes in the UVVis spectra, upon heating the protein sample to above 86 °C. The last column shows the CD spectra at increasing temperatures up to 95 °C.

using the RFAA model (best of three predictions by PLDDT) and plot the binding site RMSD between predictions with and without the ligand in figure (Figure S3E). We observe some differences between predictions made with and without ligand partner present - in particular, RFAA is capable of predicting conformational shifts for which both apo and holo states are well-represented in its training set. However,

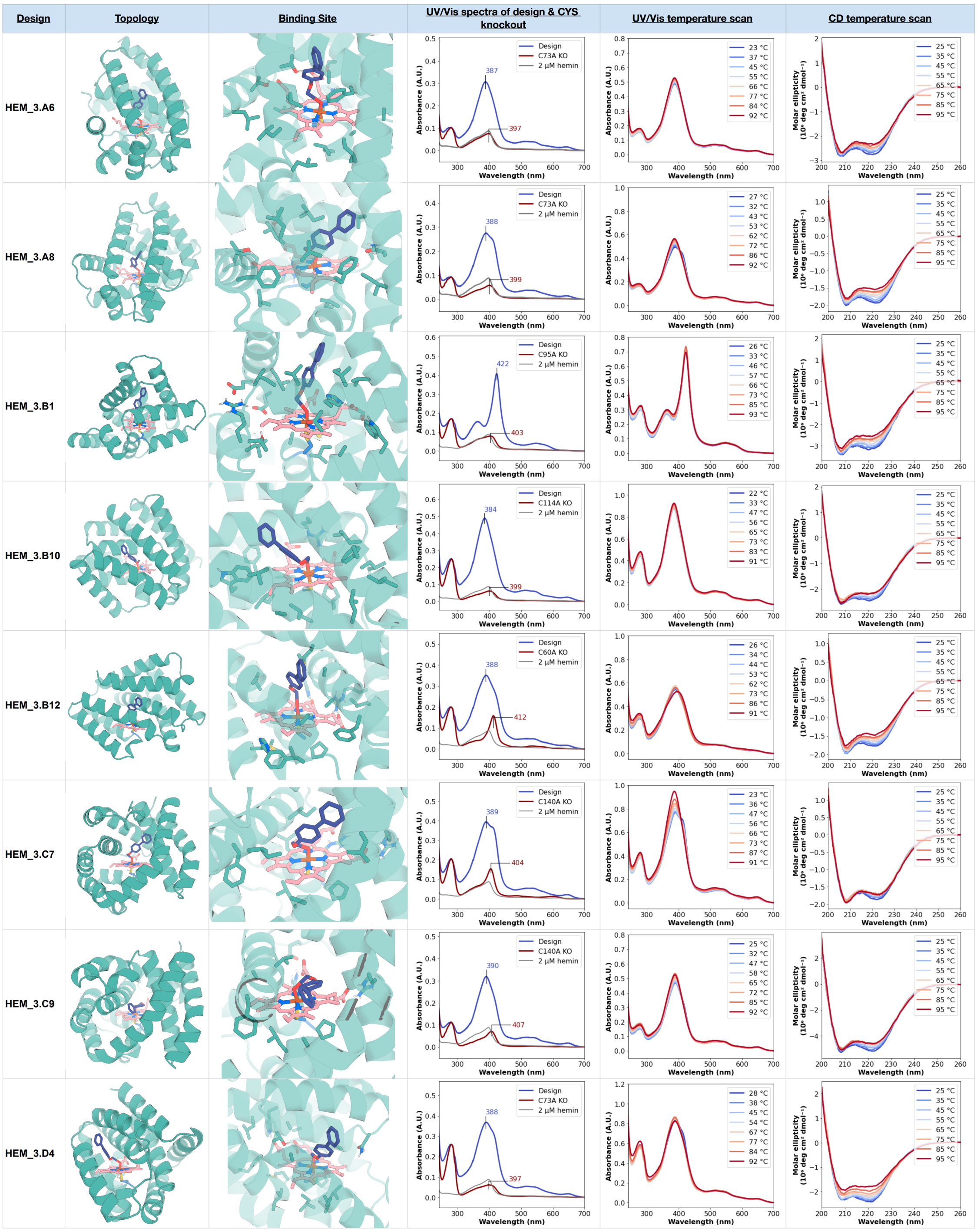

Supplementary Information Figure 10: Characterization of heme-binding proteins obtained with RFdiffusionAA, HEM_3.A6 to HEM_3.D4. The 4th column shows the UVVis spectra of the purified heme-loaded protein (blue), and the putative axial CysAla mutant at 10 $\mu$M, mixed with 2 $\mu$M hemin (red trace), along with free hemin at 2 $\mu$M (gray). The 5th column shows changes in the UVVis spectra, upon heating the protein sample to above 86 °C. The last column shows the CD spectra at increasing temperatures up to 95 °C.

we note that it is difficult in general to evaluate how well RFAA changes to a protein upon binding and do not expect the model to generalize to completely novel conformational changes. We expect that future work will make intentional train/test splits that exclude specific conformational changes to assess whether the network can generalize to novel conformational shifts but consider that outside the scope of

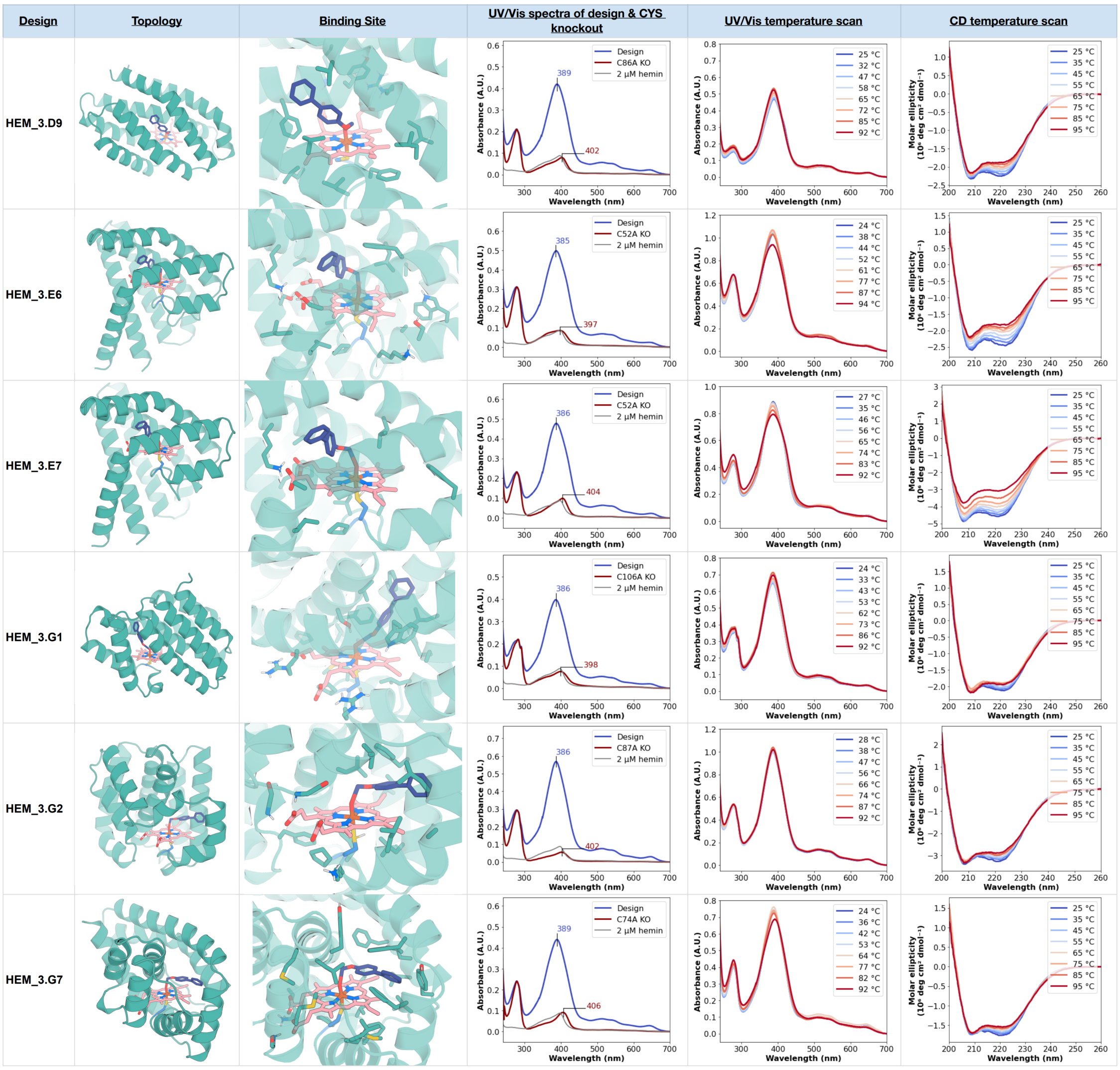

Supplementary Information Figure 11: Characterization of heme-binding proteins obtained with RFdiffusionAA, HEM_3.D9 to HEM_3.G7. The 4th column shows the UVVis spectra of the purified heme-loaded protein (blue), and the putative axial CysAla mutant at 10 $\mu$M, mixed with 2 $\mu$M hemin (red trace), along with free hemin at 2 $\mu$M (gray). The 5th column shows changes in the UVVis spectra, upon heating the protein sample to above 86 °C. The last column shows the CD spectra at increasing temperatures up to 95 °C.

the work presented in this manuscript.

#### 12.2 CAMEO Baseline Servers

In Figure S2A-B, we present a per-ligand breakdown of the targets predicted in the CAMEO challenge by both the RFAA server and the CAMEO baselines based on classical docking methods from weeks 08/12/2023 to 09/02/2023. The CAMEO baselines first predict the structure of the protein chain(s) using the SWISS-MODEL software and then dock the ligands using AutoDock Vina sequentially with a bounding box that covers the volume of the entire protein [51, 52]. The only difference between between the "Vina" and "AD4" baselines is the scoring function used when running AutoDock Vina (vina or AutoDock 4).

We show some common failure modes of the network in Figure S2D. The most common failure mode

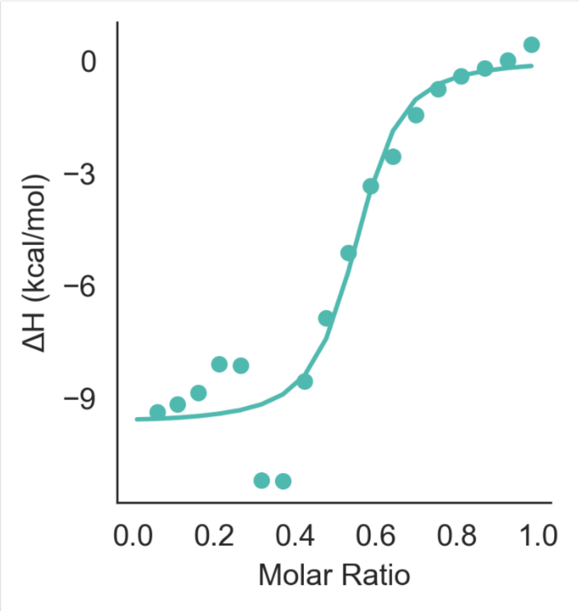

Supplementary Information Figure 12: Secondary binding affinity dataset using ITC. $K_\mathrm{d}$ estimate: 154 nM +− 111 nM.

is placing the molecule in a different orientation than the crystallized pose. These orientations usually also have shape and chemical complementarity to the protein structure but there is another potentially "lower energy" dock that is not sampled. Other failure modes we noticed in CAMEO is that the network predicts regions that are unresolved in the true structure which change the geometry of the binding pocket and thus change the dock of the small molecule. The final common failure mode is choosing the predict more buried positions for complexes that interact on the protein surface.

### References

[1] Minkyung Baek, Ivan Anishchenko, Ian R Humphreys, Qian Cong, David Baker, and Frank DiMaio. Efficient and accurate prediction of protein structure using RoseTTAFold2. May 2023.

[2] Michael Remmert, Andreas Biegert, Andreas Hauser, and Johannes Söding. HHblits: lightning-fast iterative protein sequence searching by HMM-HMM alignment. *Nat. Methods*, 9(2):173–175, December 2011.

[3] Martin Steinegger, Markus Meier, Milot Mirdita, Harald Vöhringer, Stephan J Haunsberger, and Johannes Söding. HH-suite3 for fast remote homology detection and deep protein annotation. *BMC Bioinformatics*, 20(1):473, September 2019.

[4] Helen M Berman, John Westbrook, Zukang Feng, Gary Gilliland, T N Bhat, Helge Weissig, Ilya N Shindyalov, and Philip E Bourne. The protein data bank. *Nucleic Acids Res.*, 28(1):235–242, January 2000.

[5] John Jumper, Richard Evans, Alexander Pritzel, Tim Green, Michael Figurnov, Olaf Ronneberger, Kathryn Tunyasuvunakool, Russ Bates, Augustin Žídek, Anna Potapenko, Alex Bridgland, Clemens Meyer, Simon A A Kohl, Andrew J Ballard, Andrew Cowie, Bernardino Romera-Paredes, Stanislav Nikolov, Rishub Jain, Jonas Adler, Trevor Back, Stig Petersen, David Reiman, Ellen Clancy, Michal Zielinski, Martin Steinegger, Michalina Pacholska, Tamas Berghammer, Sebastian Bodenstein, David Silver, Oriol Vinyals, Andrew W Senior, Koray Kavukcuoglu, Pushmeet Kohli, and Demis Hassabis. Highly accurate protein structure prediction with AlphaFold. *Nature*, 596(7873): 583–589, August 2021.

[6] Chloe Hsu, Robert Verkuil, Jason Liu, Zeming Lin, Brian Hie, Tom Sercu, Adam Lerer, and Alexander Rives. Learning inverse folding from millions of predicted structures. In Kamalika Chaudhuri, Stefanie Jegelka, Le Song, Csaba Szepesvari, Gang Niu, and Sivan Sabato, editors, *Proceedings of the 39th International Conference on Machine Learning*, volume 162 of *Proceedings of Machine Learning Research*, pages 8946–8970. PMLR, 2022.

[7] Minkyung Baek, Ryan McHugh, Ivan Anishchenko, David Baker, and Frank DiMaio. Accurate prediction of nucleic acid and protein-nucleic acid complexes using RoseTTAFoldNA. September 2022.

[13] Minkyung Baek, Frank DiMaio, Ivan Anishchenko, Justas Dauparas, Sergey Ovchinnikov, Gyu Rie Lee, Jue Wang, Qian Cong, Lisa N Kinch, R Dustin Schaeffer, Claudia Millán, Hahnbeom Park, Carson Adams, Caleb R Glassman, Andy DeGiovanni, Jose H Pereira, Andria V Rodrigues, Alberdina A van Dijk, Ana C Ebrecht, Diederik J Opperman, Theo Sagmeister, Christoph Buhlheller, Tea Pavkov-Keller, Manoj K Rathinaswamy, Udit Dalwadi, Calvin K Yip, John E Burke, K Christopher Garcia, Nick V Grishin, Paul D Adams, Randy J Read, and David Baker. Accurate prediction of protein structures and interactions using a three-track neural network. *Science*, 373(6557):871–876, August 2021.

[14] Roshan M Rao, Jason Liu, Robert Verkuil, Joshua Meier, John Canny, Pieter Abbeel, Tom Sercu, and Alexander Rives. MSA transformer. In Marina Meila and Tong Zhang, editors, *Proceedings of the 38th International Conference on Machine Learning*, volume 139 of *Proceedings of Machine Learning Research*, pages 8844–8856. PMLR, 2021.

[15] Nick Bhattacharya, Neil Thomas, Roshan Rao, Justas Daupras, Peter K Koo, David Baker, Yun S Song, and Sergey Ovchinnikov. Single layers of attention suffice to predict protein contacts. October 2020.

[16] Fabian B Fuchs, Daniel E Worrall, Volker Fischer, and Max Welling. SE(3)-Transformers: 3D Roto-Translation equivariant attention networks. June 2020.

[17] Nathaniel Thomas, Tess Smidt, Steven Kearnes, Lusann Yang, Li Li, Kai Kohlhoff, and Patrick Riley. Tensor field networks: Rotation- and translation-equivariant neural networks for 3D point clouds. February 2018.

[18] Adam Paszke, Sam Gross, Francisco Massa, Adam Lerer, James Bradbury, Gregory Chanan, Trevor Killeen, Zeming Lin, Natalia Gimelshein, Luca Antiga, Alban Desmaison, Andreas Köpf, Edward Yang, Zach DeVito, Martin Raison, Alykhan Tejani, Sasank Chilamkurthy, Benoit Steiner, Lu Fang, Junjie Bai, and Soumith Chintala. PyTorch: An imperative style, High-Performance deep learning library. December 2019.

[19] Jacob Devlin, Ming-Wei Chang, Kenton Lee, and Kristina Toutanova. BERT: Pre-training of deep bidirectional transformers for language understanding. In *Proceedings of the 2019 Conference of the North American Chapter of the Association for Computational Linguistics: Human Language Technologies, Volume 1 (Long and Short Papers)*, pages 4171–4186, Minneapolis, Minnesota, June 2019. Association for Computational Linguistics.

[20] Jianyi Yang, Ivan Anishchenko, Hahnbeom Park, Zhenling Peng, Sergey Ovchinnikov, and David Baker. Improved protein structure prediction using predicted interresidue orientations. *Proceedings of the National Academy of Sciences*, 117(3):1496–1503, 2020.

[21] Hannes Stärk, Octavian-Eugen Ganea, Lagnajit Pattanaik, Regina Barzilay, and Tommi Jaakkola. EquiBind: Geometric deep learning for drug binding structure prediction. February 2022.

[22] Rebecca F Alford, Andrew Leaver-Fay, Jeliazko R Jeliazkov, Matthew J O'Meara, Frank P DiMaio, Hahnbeom Park, Maxim V Shapovalov, P Douglas Renfrew, Vikram K Mulligan, Kalli Kappel, Jason W Labonte, Michael S Pacella, Richard Bonneau, Philip Bradley, Roland L Dunbrack, Jr, Rhiju Das, David Baker, Brian Kuhlman, Tanja Kortemme, and Jeffrey J Gray. The rosetta All-Atom energy function for macromolecular modeling and design. *J. Chem. Theory Comput.*, 13(6): 3031–3048, June 2017.

[23] A K Rappe, C J Casewit, K S Colwell, W A Goddard, Iii, and W M Skiff. UFF, a full periodic table force field for molecular mechanics and molecular dynamics simulations. *J. Am. Chem. Soc.*, 114(25):10024–10035, December 1992.

[24] Yang Zhang and Jeffrey Skolnick. Scoring function for automated assessment of protein structure template quality. *Proteins*, 57(4):702–710, December 2004.

[25] Ivan Anishchenko, Minkyung Baek, Hahnbeom Park, Naozumi Hiranuma, David E Kim, Justas Dauparas, Sanaa Mansoor, Ian R Humphreys, and David Baker. Protein tertiary structure prediction and refinement using deep learning and rosetta in CASP14. *Proteins*, 89(12):1722–1733, December 2021.

[26] UniProt: the universal protein knowledgebase in 2023. *Nucleic Acids Res.*, 51(D1):D523–D531, 2023.

[27] I-Min A Chen, Victor M Markowitz, Ken Chu, Krishna Palaniappan, Ernest Szeto, Manoj Pillay, Anna Ratner, Jinghua Huang, Evan Andersen, Marcel Huntemann, Neha Varghese, Michalis Hadjithomas, Kristin Tennessen, Torben Nielsen, Natalia N Ivanova, and Nikos C Kyrpides. IMG/M: integrated genome and metagenome comparative data analysis system. *Nucleic Acids Res.*, 45(D1): D507–D516, January 2017.

[28] Sean R Eddy. Accelerated profile HMM searches. *PLoS Comput. Biol.*, 7(10):e1002195, October 2011.

[29] Martin Buttenschoen, Garrett M Morris, and Charlotte M Deane. PoseBusters: AI-based docking methods fail to generate physically valid poses or generalise to novel sequences. August 2023.

[30] Christiam Camacho, George Coulouris, Vahram Avagyan, Ning Ma, Jason Papadopoulos, Kevin Bealer, and Thomas L Madden. BLAST+: architecture and applications. *BMC Bioinformatics*, 10: 421, December 2009.

[31] F. Pedregosa, G. Varoquaux, A. Gramfort, V. Michel, B. Thirion, O. Grisel, M. Blondel, P. Prettenhofer, R. Weiss, V. Dubourg, J. Vanderplas, A. Passos, D. Cournapeau, M. Brucher, M. Perrot, and E. Duchesnay. Scikit-learn: Machine learning in Python. *Journal of Machine Learning Research*, 12:2825–2830, 2011.

[32] Hahnbeom Park, Guangfeng Zhou, Minkyung Baek, David Baker, and Frank DiMaio. Force field optimization guided by small molecule crystal lattice data enables consistent Sub-Angstrom Protein–Ligand docking. *J. Chem. Theory Comput.*, 17(3):2000–2010, March 2021.

[33] Joseph L Watson, David Juergens, Nathaniel R Bennett, Brian L Trippe, Jason Yim, Helen E Eisenach, Woody Ahern, Andrew J Borst, Robert J Ragotte, Lukas F Milles, Basile I M Wicky, Nikita Hanikel, Samuel J Pellock, Alexis Courbet, William Sheffler, Jue Wang, Preetham Venkatesh, Isaac Sappington, Susana Vázquez Torres, Anna Lauko, Valentin De Bortoli, Emile Mathieu, Sergey Ovchinnikov, Regina Barzilay, Tommi S Jaakkola, Frank DiMaio, Minkyung Baek, and David Baker. De novo design of protein structure and function with RFdiffusion. *Nature*, 620(7976):1089–1100, August 2023.

[34] Longxing Cao, Brian Coventry, Inna Goreshnik, Buwei Huang, William Sheffler, Joon Sung Park, Kevin M Jude, Iva Marković, Rameshwar U Kadam, Koen H G Verschueren, Kenneth Verstraete, Scott Thomas Russell Walsh, Nathaniel Bennett, Ashish Phal, Aerin Yang, Lisa Kozodoy, Michelle DeWitt, Lora Picton, Lauren Miller, Eva-Maria Strauch, Nicholas D DeBouver, Allison Pires,

Asim K Bera, Samer Halabiya, Bradley Hammerson, Wei Yang, Steffen Bernard, Lance Stewart, Ian A Wilson, Hannele Ruohola-Baker, Joseph Schlessinger, Sangwon Lee, Savvas N Savvides, K Christopher Garcia, and David Baker. Design of protein-binding proteins from the target structure alone. *Nature*, 605(7910):551–560, May 2022.

[35] M. J. Frisch, G. W. Trucks, H. B. Schlegel, G. E. Scuseria, M. A. Robb, J. R. Cheeseman, G. Scalmani, V. Barone, G. A. Petersson, H. Nakatsuji, X. Li, M. Caricato, A. V. Marenich, J. Bloino, B. G. Janesko, R. Gomperts, B. Mennucci, H. P. Hratchian, J. V. Ortiz, A. F. Izmaylov, J. L. Sonnenberg, D. Williams-Young, F. Ding, F. Lipparini, F. Egidi, J. Goings, B. Peng, A. Petrone, T. Henderson, D. Ranasinghe, V. G. Zakrzewski, J. Gao, N. Rega, G. Zheng, W. Liang, M. Hada, M. Ehara, K. Toyota, R. Fukuda, J. Hasegawa, M. Ishida, T. Nakajima, Y. Honda, O. Kitao, H. Nakai, T. Vreven, K. Throssell, J. A. Montgomery, Jr., J. E. Peralta, F. Ogliaro, M. J. Bearpark, J. J. Heyd, E. N. Brothers, K. N. Kudin, V. N. Staroverov, T. A. Keith, R. Kobayashi, J. Normand, K. Raghavachari, A. P. Rendell, J. C. Burant, S. S. Iyengar, J. Tomasi, M. Cossi, J. M. Millam, M. Klene, C. Adamo, R. Cammi, J. W. Ochterski, R. L. Martin, K. Morokuma, O. Farkas, J. B. Foresman, and D. J. Fox. Gaussian~16 Revision C.01, 2016. Gaussian Inc. Wallingford CT.

[36] Stefan Grimme, Stephan Ehrlich, and Lars Goerigk. Effect of the damping function in dispersion corrected density functional theory. *Journal of Computational Chemistry*, 32(7):1456–1465, 2011. doi: https://doi.org/10.1002/jcc.21759. URL `https://onlinelibrary.wiley.com/doi/abs/10.1002/jcc.21759`.

[37] Christoph Bannwarth, Eike Caldeweyher, Sebastian Ehlert, Andreas Hansen, Philipp Pracht, Jakob Seibert, Sebastian Spicher, and Stefan Grimme. Extended tight-binding quantum chemistry methods. *WIREs Computational Molecular Science*, 11(2):e1493, 2021. doi: https://doi.org/10.1002/wcms.1493. URL `https://wires.onlinelibrary.wiley.com/doi/abs/10.1002/wcms.1493`.

[38] Susana Vázquez Torres, Philip J Y Leung, Isaac D Lutz, Preetham Venkatesh, Joseph L Watson, Fabian Hink, Huu-Hien Huynh, Andy Hsien-Wei Yeh, David Juergens, Nathaniel R Bennett, Andrew N Hoofnagle, Eric Huang, Michael J MacCoss, Marc Expòsit, Gyu Rie Lee, Paul M Levine, Xinting Li, Mila Lamb, Elif Nihal Korkmaz, Jeff Nivala, Lance Stewart, Joseph M Rogers, and David Baker. De novo design of high-affinity protein binders to bioactive helical peptides. December 2022.

[50] Pauli Virtanen, Ralf Gommers, Travis E. Oliphant, Matt Haberland, Tyler Reddy, David Cournapeau, Evgeni Burovski, Pearu Peterson, Warren Weckesser, Jonathan Bright, Stéfan J. van der Walt, Matthew Brett, Joshua Wilson, K. Jarrod Millman, Nikolay Mayorov, Andrew R. J. Nelson, Eric Jones, Robert Kern, Eric Larson, C J Carey, İlhan Polat, Yu Feng, Eric W. Moore, Jake VanderPlas, Denis Laxalde, Josef Perktold, Robert Cimrman, Ian Henriksen, E. A. Quintero, Charles R. Harris, Anne M. Archibald, Antônio H. Ribeiro, Fabian Pedregosa, Paul van Mulbregt, and SciPy 1.0 Contributors. SciPy 1.0: Fundamental Algorithms for Scientific Computing in Python. *Nature Methods*, 17:261–272, 2020. doi: 10.1038/s41592-019-0686-2.

[51] Aurelien Grosdidier, Vincent Zoete, and Olivier Michielin. Swissdock, a protein-small molecule docking web service based on eadock dss. *Nucleic acids research*, 39(suppl_2):W270–W277, 2011.

[52] Oleg Trott and Arthur J Olson. Autodock vina: improving the speed and accuracy of docking with a new scoring function, efficient optimization, and multithreading. *Journal of computational chemistry*, 31(2):455–461, 2010.
